# Tuning Interdomain Conjugation Toward *in situ* Population Modification in Yeast

**DOI:** 10.1101/2023.09.12.557379

**Authors:** Kevin R Stindt, Megan N McClean

## Abstract

The ability to modify and control natural and engineered microbiomes is essential for biotechnology and biomedicine. Fungi are critical members of most microbiomes, yet technology for modifying the fungal members of a microbiome has lagged far behind that for bacteria. Interdomain conjugation (IDC) is a promising approach, as DNA transfer from bacterial cells to yeast enables *in situ* modification. While such genetic transfers have been known to naturally occur in a wide range of eukaryotes, and are thought to contribute to their evolution, IDC has been understudied as a technique to control fungal or fungal-bacterial consortia. One major obstacle to widespread use of IDC is its limited efficiency. In this work, we utilize interactions between genetically tractable *Escherichia coli* and *Saccharomyces cerevisiae* to control the incidence of IDC. We test the landscape of population interactions between the bacterial donors and yeast recipients to find that bacterial commensalism leads to maximized IDC, both in culture and in mixed colonies. We demonstrate the capacity of cell-to-cell binding via mannoproteins to assist both IDC incidence and bacterial commensalism in culture, and model how these tunable controls can predictably yield a range of IDC outcomes. Further, we demonstrate that these lessons can be utilized to lastingly alter a recipient yeast population, by both “rescuing” a poor-growing recipient population and collapsing a stable population via a novel IDC-mediated CRISPR/Cas9 system.

## Introduction

Extraordinary advances have been made in recent years elucidating the composition and function of microbiome members, but the vast majority of this work has focused on bacterial species, and often overlooks fungal participants^1^. Though the number of fungal cells is typically dwarfed by bacterial cells, fungi play important roles in both human^2–6^ and environmental^7–10^ microbiomes. Many fungal pathogens live as commensals in humans before becoming infectious, whether due to hospital-derived nosocomial infections or auto-immune disorders, both of which are on the rise^5^. *Candida* species, which usually exist as commensals in the gut microbiome, cause many such nosocomial infections, resulting in a range of candidiasis symptoms that can lead to sepsis^2^. Common skin microbiome residents in the *Malassezia* genus^6^ have been implicated in Crohn’s disease^3^ and tumorigenisis^4^. Moreover, many fungi infect plant^7^ or other animal species^8^, e.g. bats^9^ and amphibians^10^, often causing significant agricultural or environmental loss. In addition to their natural roles, the unique metabolic capabilities of fungi make them important members in food production^11,12^ and engineered bioproduction^13,14^ and bioremediation^15^, often with other species in consortia, emulating the division of labor found in microbiomes. Thus, fungal microbes play important roles in human, plant, and environmental microbiomes, in addition to synthetic microbial consortia, and therefore tools for modifying the fungal microbiome are critical for advances in all these areas.

Modification of microbiome members is most frequently done by methods designed to reduce or kill off specific populations in a targeted or semi-targeted way^16^. Pre and probiotics, antibiotics and antifungals, microbial transplants, and phages, seek to promote or eliminate specific populations. In addition to focusing primarily on eliminating specific populations, few of these tools are targeting for modifying fungal microbiome members. In contrast, bacterial conjugation, a naturally occurring form of horizontal gene transfer (HGT)^17^ allows modification of bacterial populations instead of simple killing and has already been used for probiotics^18^, defense against antibiotic-resistant pathogens^19^, crop modification for desired traits^20^, control of undomesticated microbial species^21^, circuit-like control of synthetic consortia^22^, and *in situ* microbiome engineering^23,24^. Bacteria also conjugate with a variety of eukaryotic recipient cells, most commonly from bacterial donor *Agrobacterium tumefaciens* to plant cells^25^. And while *A. tumefaciens* is uniquely well-studied for performing interdomain conjugation (IDC) in the wild, highly genetically tractable bacteria, such as *Escherichia coli,* can be modified to perform IDC with diatoms^26–29^, mammalian cells^30^, and multiple yeast species^31–35^, offering a powerful opportunity for modifying fungi *in situ*. IDC has typically been referred to as transkingdom conjugation (TKC), though this nomenclature predates^31^ the domain designation of prokaryotes and eukaryotes proposed by Woese^36^, which further highlights the significance of this genetic transfer mechanism between cells of different domains.

Conjugative transfer of DNA occurs in multiple stages in the bacterial cell. First, a complex of proteins called the “relaxosome”, containing catalytic relaxases, nicks the conjugative plasmid at the origin of transfer (*ori^T^*), and transfers one strand of the plasmid DNA to the membrane-bound type IV secretion system (T4SS)^37^. The T4SS transports the relaxosome-DNA complex through both bacterial membranes and a pilus connecting the donor and recipient cells. For *E. coli* T4SS, the DNA re-circularizes in the recipient cell to recreate the original plasmid^38^. Conjugation can occur via either a *cis* mechanism, in which the plasmid carrying the relaxosome genes itself contains an *ori^T^* and thus is transferred to a recipient cell, or a *trans* mechanism, in which the *ori^T^* is on a separate plasmid, which gets transferred^39^ (Fig 1a).

**Figure 1:**
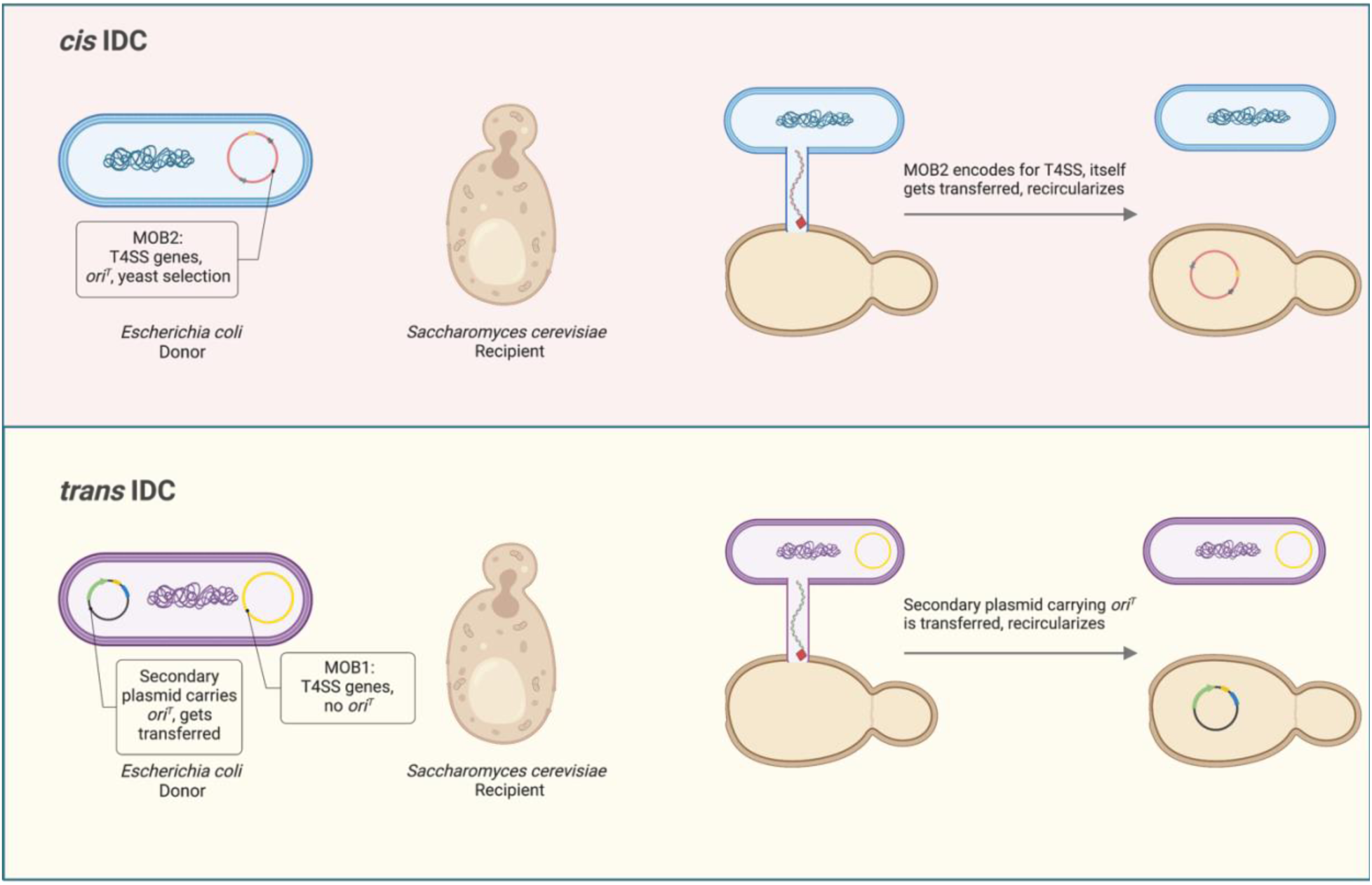
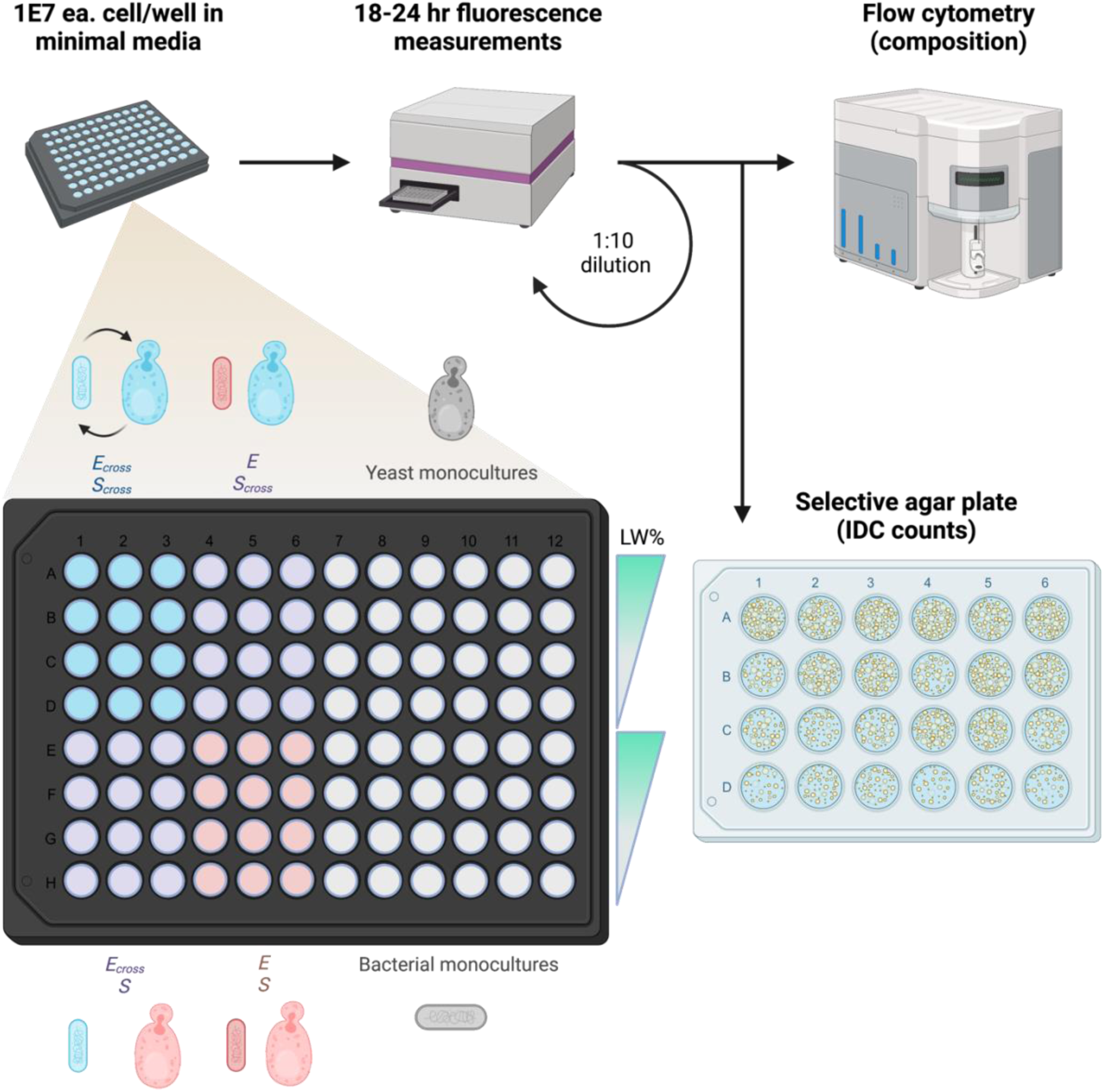
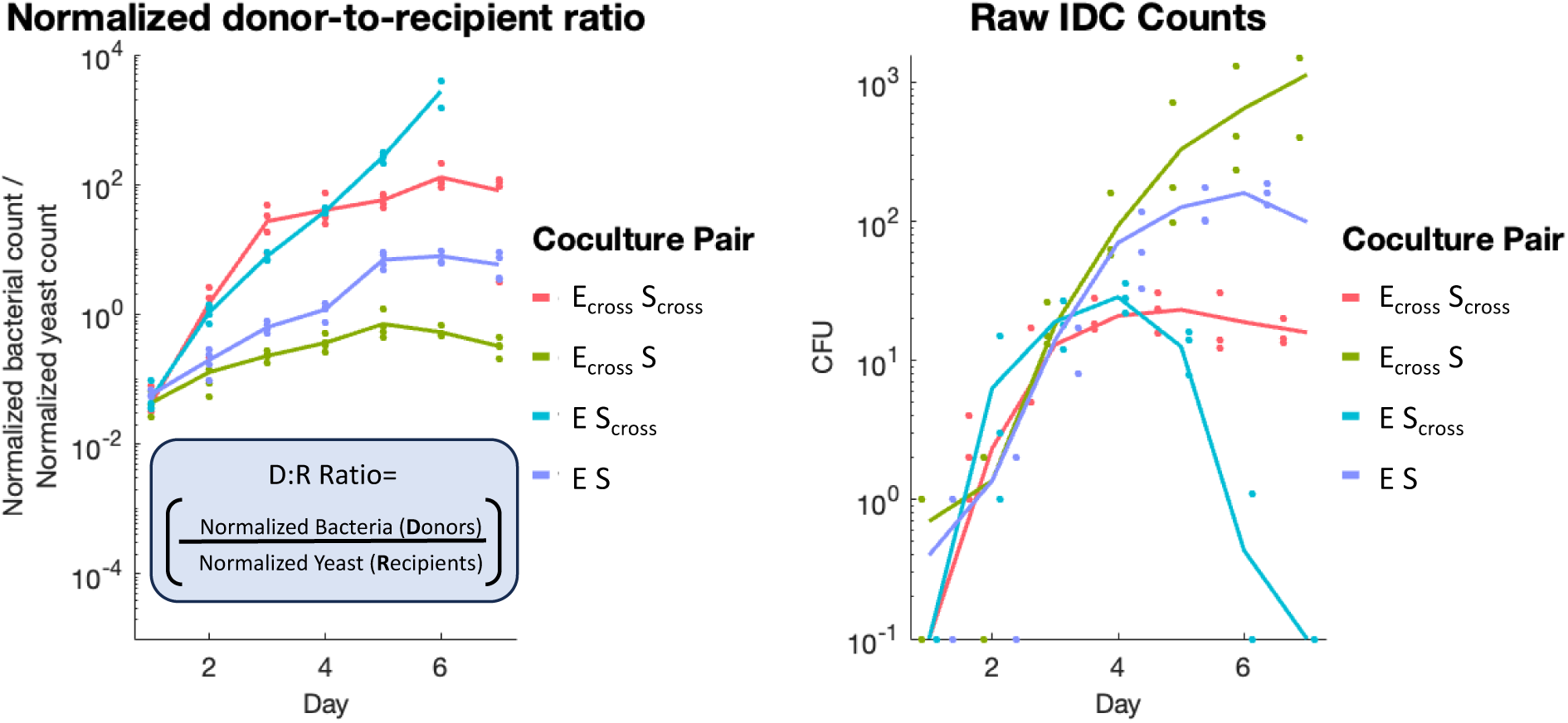
Setup of IDC cultures with metabolic crossfeeding. A. <u>Types of IDC transfer.</u> For this IncP T4SS system, DNA transfer to recipient yeasts can either occur in *cis* or in *trans*. For *cis*-IDC, the plasmid encoding the T4SS (pTA-Mob 2.0) also contains yeast centromeric DNA maintenance machinery, yeast selection genes HIS3 and URA3, and the transfer recognition sequence, oriT, at which the relaxosome nicks plasmid DNA and transfers it through the pilus to the recipient. In *trans*-IDC, the T4SS-encoding plasmid (pTA-Mob 1.0) lack the sequences for yeast maintenance, yeast selection, and the oriT sequence. Thus, a second plasmid is required for *trans*-IDC, which carries these elements and is transferred to recipients. B. <u>Experimental setup of batch cultures</u>. Cells are combined in a 96-well microplate with varying levels of leucine (L) and tryptophan (W) in a minimal media cocktail. Cocultures and monocultures are incubated at 30°C with continuous shaking and measured for fluorescence of each species every 15 minutes. After 18-24 hours of growth, cells are diluted 1:10 into new media to continue growing. Simultaneously, a 1:10 dilution of cells is prepared for flow cytometry, and an undiluted 100 uL is plated onto IDC-selective plates. C. <u>Example of culture composition measurements.</u> Traces of cells in culture, measured either by plate-reader fluorescence (left) or flow cytometry cell count (right), normalized by dividing each reading by the maximum for that measurement type after day 1. For example, mCherry (bacterial) measurements are all divided by the maximum mCherry reading for all samples during experiment after day 1. Cell counts are normalized per species, fluorescence per fluorophore. Red traces denote bacterial growth, brown traces, yeast growth. Solid lines represent cell growth in coculture (mean of four replicates), dotted lines growth in monoculture (mean of two replicates). Shaded regions are standard deviation from mean. Blue circles in the fluorescence plot denote times at which samples are diluted for batch culturing and additional measurements. Example shown is crossfeeding pair (E_cross_ S_cross_) at 15% Leu and Trp (15% LW).

IDC is currently limited as a tool for microbiome modification by its relatively low efficiency^40^. In fact, the vast majority of conjugation research has focused on lowering efficiency further^40,41^, in an effort to prevent the spread of antibiotic resistance, which occurs through conjugative transfer of resistance-coding genes^42^. Conjugation rates between *E. coli* and the genetically tractable yeast species *Saccharomyces cerevisiae* are typically below 1 in 1,000 yeast cells^43^ (vs. ∼1 in 100 efficiency per cell for 1 μg DNA in a LiAc transformation^44^), though recent work has succeeded in generating 10-fold higher efficiencies by selectively mutating the T4SS machinery^39^. Synthetic approaches also exist for increasing efficiency, such as colocalizing donor and recipient cells on beads, but have limited utility outside of laboratory settings^45^. Bacterial recipients of horizontal gene transfer can subsequently serve as conjugative donors, leading to an exponential increase in conjugated cells^45^. In contrast, IDC recipients are unable to subsequently serve as donors. This is ideal for biocontainment, but further limits efficiency. In the laboratory, it is easy to overcome low conjugation efficiencies by selecting specifically for transconjugants and subculturing them. However, whether IDC can be optimized to modify enough individuals to affect population level outcomes, as would be required for *in situ* modifications or control of engineering consortia, has not been tested.

In this work, we explore strategies for controlling IDC to affect population-level outcomes. One such strategy is to tune donor and recipient population ratios, since IDC rates are considered to be dependent on the frequency of donor-recipient interactions, and some work has demonstrated an increase in IDC for higher donor-to-recipient ratios, albeit on shorter time scales^43^. We thus use strains of *E. coli* and *S. cerevisiae* mutated to allow tunable population control via engineered crossfeeding between *E. coli* and *S. cerevisiae*, in which each species is auxotrophic for an essential amino acid that the other species overproduces. This approach also has implications in colony settings, which more closely match the dense biofilm environments in which most microbes naturally exist^46^. Here, conjugation events between two populations occurs along population boundaries^47^, and mutualism between cells can greatly increase intermixing of populations in both bacteria^48^ and yeast^49^, hypothetically creating more population boundaries along which IDC can occur. We also probe the spatial dynamics of supposedly well-mixed cultures, since *E. coli* and *S. cerevisiae* are known to form mixed cellular aggregates via mannoprotein binding of *E. coli*^50^, and determine how these short-range interactions affect both population dynamics and IDC. We model IDC in culture via a series of ordinary differential equations (ODEs), to predict both IDC transfer rates and conditions for which IDC is optimized. Finally, to verify whether these tunable population knobs can cause population-scale recipient changes via IDC, we apply them to control IDC-rescuing and IDC-killing of recipient yeast populations.

## Results

### Crossfeeding mixed cultures and automation enable control and measurement of population ratios

To enable tunable control of population ratios, we designed strains of *E. coli* and *S. cerevisiae* to be obligate mutualists when deprived of specific nutrients. We utilized a yeast strain^49^ that is auxotrophic for tryptophan (Trp^-^, *Δtrp2*, encoding anthranilate synthase, precursor to tryptophan synthesis^51^) and overproduces leucine (Leu^++^, *LEU4^FBR^*via feedback resistant mutation^52^), carrying a genomically-integrated constitutive ymCitrine fluorescent reporter (*his3Δ::prACT1-ymCitrine-tADH::HIS3MX6*). We also developed a corresponding leucine-auxotrophic, tryptophan-overproducing crossfeeder *E. coli* (Leu^-^, Trp^++^, via *ΔleuA*—responsible for leucine intermediate 2-isopropylmalate synthesis^53^—and *ΔtrpR*, the canonical Trp repressor) that constitutively expresses mCherry episomally. Along with corresponding strains of *S. cerevisiae* and *E. coli* unmodified for *LEU* and *TRP* (hereafter referred to as WT strains, “S” and “E”, respectively), these “crossfeeder” strains (“S_cross_” and “E_cross_”) allowed us to tune the concentrations of leucine and tryptophan in mixed culture to alter steady state cell ratios.

To track the dynamics of each strain in mixed culture in addition to IDC counts over several days, we employed a multifaceted measurement scheme (Fig 1b). Each cell pairing was batch cultured in a 96-well plate with various concentrations of leucine and tryptophan (“% LW”) and measured for fluorescence of each strain (ymCitrine for yeast, mCherry for bacteria) in 15 minute intervals via an automated plate handler and a fluorimeter, yielding strain-specific dynamic information. After ∼18-24 hours, batch cultures were diluted 10-fold into fresh media, but in most cases were also measured via flow cytometry for verification of cell counts, allowing us to utilize the difference in cell sizes between bacteria and yeasts to optically measure individual cells of each type in a high-throughput way. This dual-measurement scheme allowed dynamic growth information while also controlling for variation in fluorescence expression due to, e.g., growth phase (Fig 1c). Furthermore, we plated mixed cultures onto IDC-selective agar media after each day of batch culturing to get raw IDC counts. We measured population effects on IDC both in *cis*— with the self-transferring plasmid pTA-Mob 2.0—and in *trans*, via a two-plasmid system including the *ori^T^-*lacking pTA-Mob 1.0 and a separate, yeast-selectable transfer plasmid (Fig 1a). After screening for optimal coculture conditions (SF1), we batch cultured cells for six days in minimal media with a range of strain-dependent leucine and tryptophan concentrations. Thus, we showed we could dynamically track batch cultures of bacteria and yeast, while also measuring cell counts for bacteria, yeast, and transconjugants each day.

### Tuning population ratios in batch culture affects the number of IDC events

To determine if we can control IDC frequencies by tuning steady state population growth, we tested all combinations of crossfeeder *E. coli* (E_cross_) and *S. cerevisiae* (S_cross_), and their WT counterparts (E and S), over a range of leucine and tryptophan concentrations (% LW). In most cases, bacterial and yeast populations failed to establish stable crossfeeding with leucine and tryptophan fully removed from media, and interactions between species seemed primarily antagonistic, with one strain’s growth corresponding to the loss of the other. Crossfeeding bacteria temporarily survived via leucine secreted by crossfeeding yeast^52^ before the latter is outcompeted, driving down both populations; in contrast, auxotrophic yeasts did not benefit similarly from Trp-overproducing bacteria^54^. Moreover, auxotrophic E_cross_ bacteria survived from WT yeasts at 0% leucine and tryptophan (0% LW), in an apparent commensal relationship, suggesting either a low but significant level of basal leucine (or synthesis intermediate) secretion from S via an unknown mechanism^55^ or sufficient yeast lysate for E_cross_ survival. WT bacteria (“E”) did not provide a similar benefit for crossfeeding yeasts (Fig 2a, SF2).

**Figure 2:**
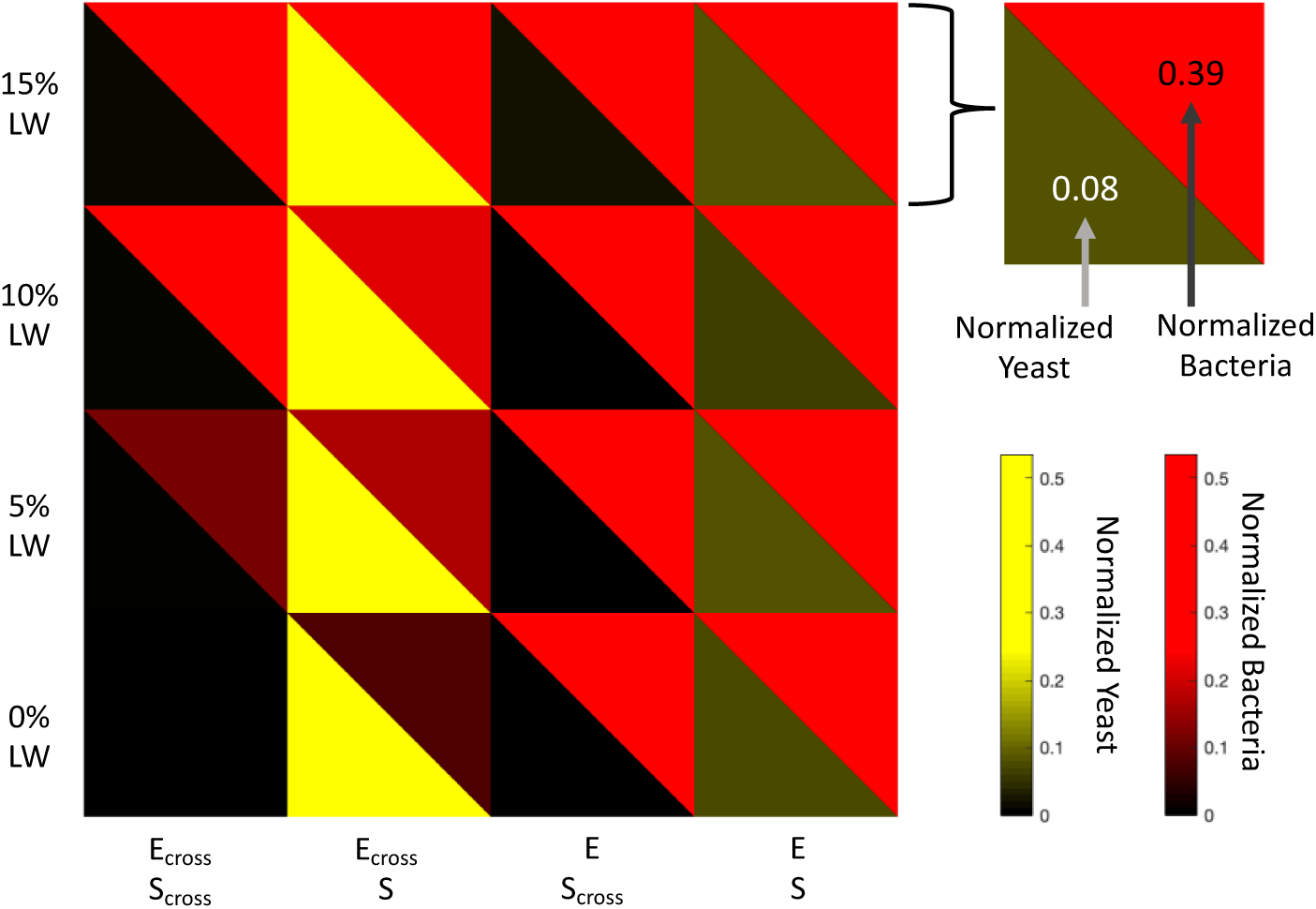
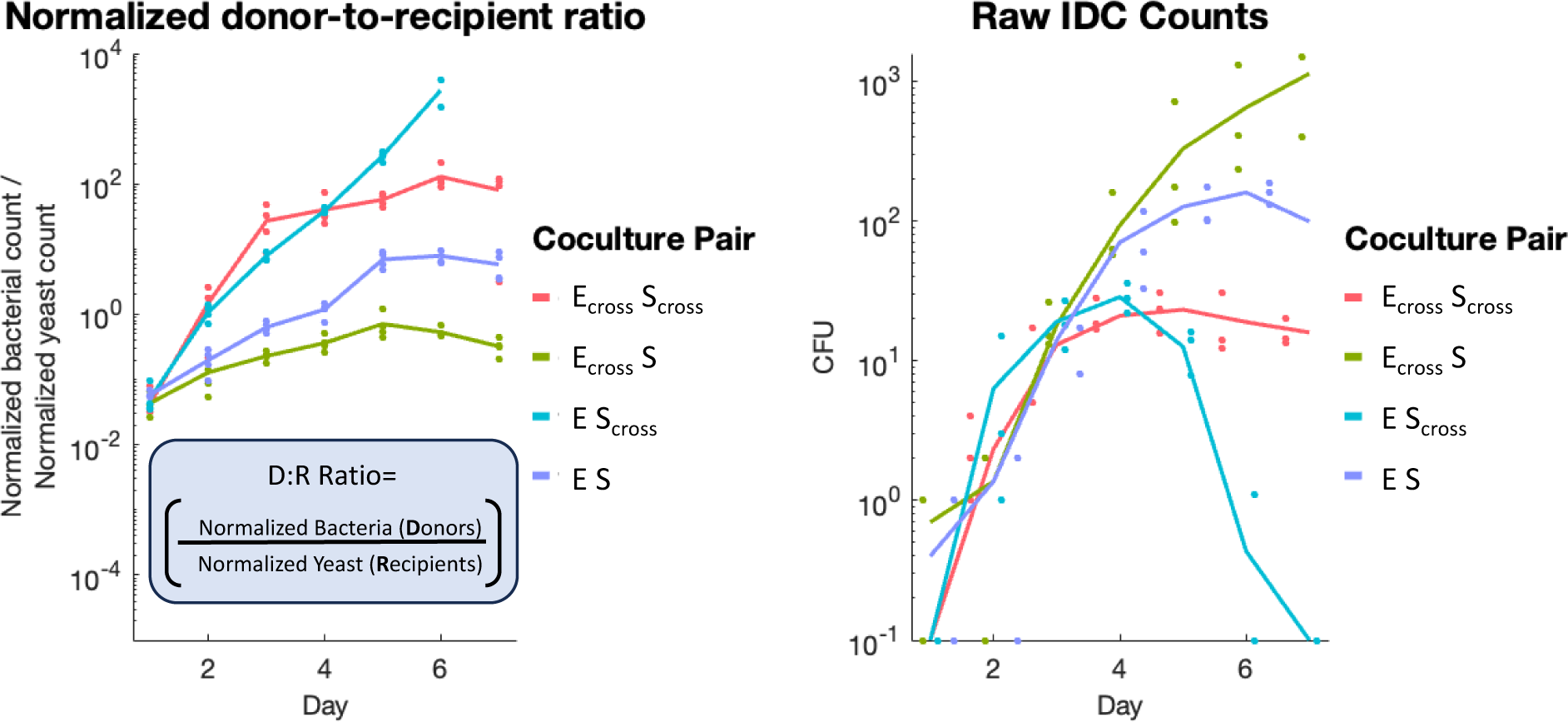
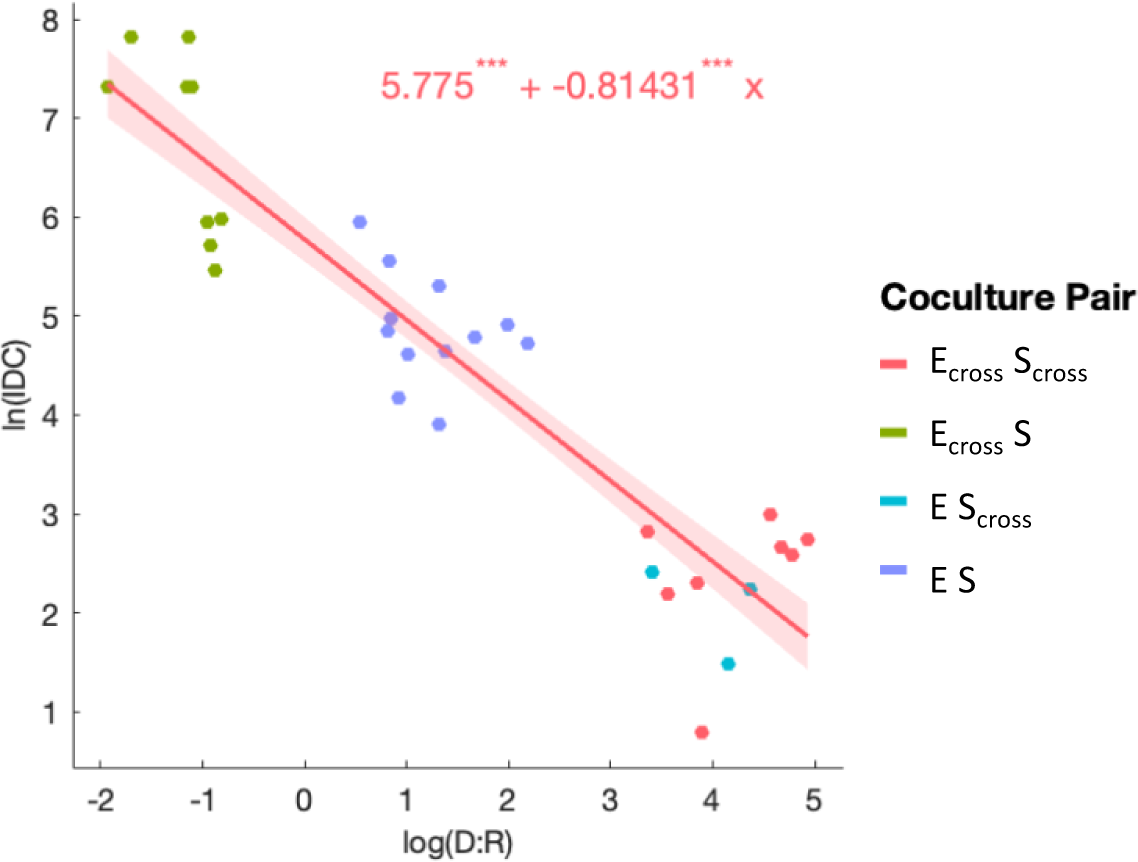
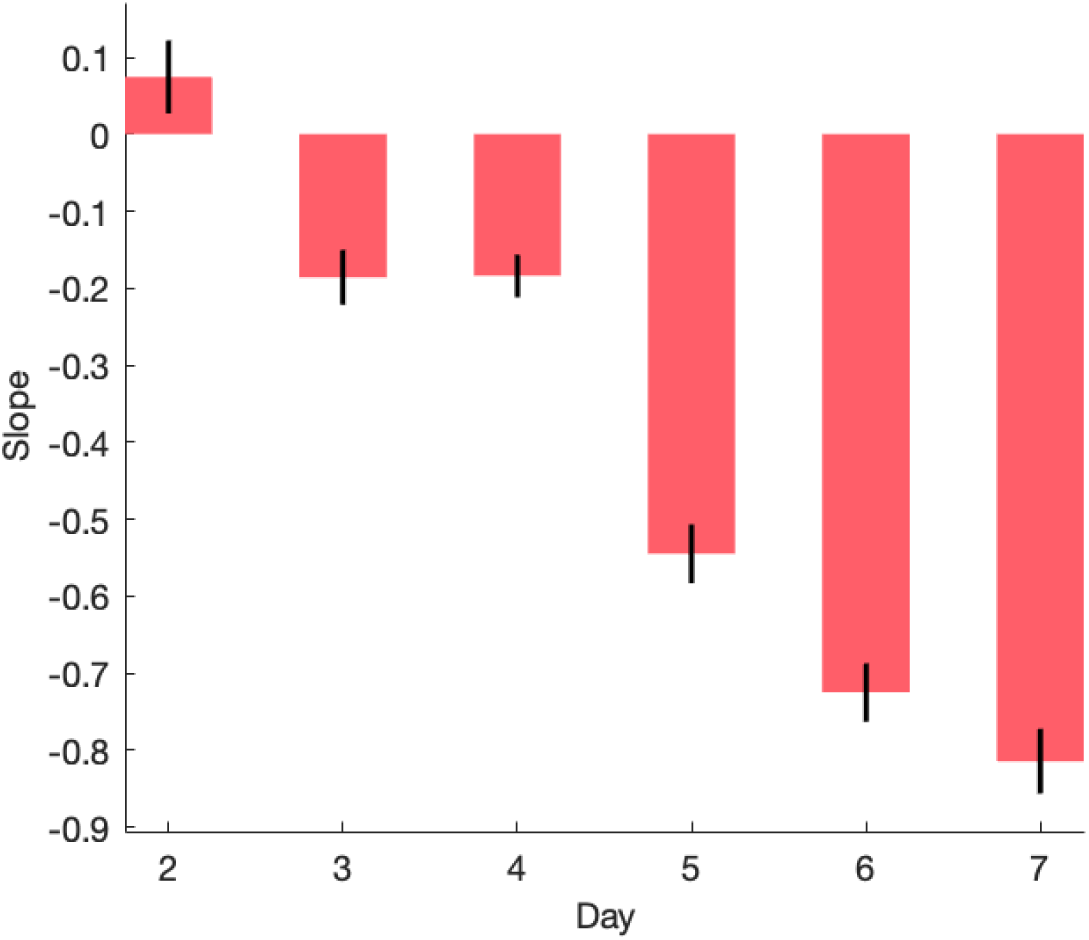
Batch culturing crossfeeding populations reveals relationships between population ratios and IDC. <u>A. Compositional outcomes of cocultures</u>. Split heatmap of bacterial cell counts (red) and yeast cell counts (yellow) for each coculture pairing (columns) over a range of leucine and tryptophan (LW) concentrations (rows), from flow cytometry of batch culture day 6 (mean of four replicates). Counts are normalized to max cell counts per species then multiplied uniformly to enhance color brightness, to better visualize low-growing populations. At 0% LW, crossfeeding pairs’ (E_cross_ S_cross_) growth is imperceptibly small, while the crossfeeder bacteria paired with WT yeast (E_cross_ S) shows bacterial commensalism, and the WT pair (E S) competitive exclusion of yeast. Experimental results are for *cis*-donors. Brightness is mean of four replicates. <u>B. Normalized donor-to-recipient ratios correspond inversely to IDC counts.</u> Ratios of normalized cell counts (cell count divided by maximum for that species across experiment) of bacterial donors and yeast recipients, calculated from flow cytometry data, plotted over time for each cell pairing, at 5% LW (left). Raw IDC counts from colony forming units (CFU) on selectable media for the same conditions and cell pairings (right). Lines represent means of four replicates (dots), colors, coculture pairings. Inset: determination of donor-to-recipient ratio (“D:R”). <u>C. Correlation between donor-to-recipient ratios and IDC is linearly inverse on log-log scale</u>. Log-log distribution of donor-to-recipient ratio and IDC counts for each cell pairing at day 7, for all LW%. IDC counts at detection limits (500 or 0.1 per 100 uL sampling, see methods) omitted. Generalized linear model fit, with normal distribution, shown in solid red line, with 95% CI in shaded region. Fit equation displayed with stars denoting *p-*value significance for each term. <u>D. Inverse correlation between D:R and IDC increases over time.</u> Slopes of log-log plot linear fits for all days of batch culture, showing decreasing slope over time. Black bars = standard error of mean.

As in previous work, we found markedly lower *trans* IDC rates relative to *cis* IDC (SF3)^56^. Contrary to previous work demonstrating higher IDC rates with more donor bacteria^43^, however, we found an inverse correlation between donor-to-recipient ratios and IDC counts over time (Fig 2b), especially for *cis* IDC (SF3). That is, more donor bacteria resulted in less conjugated yeast. This trend manifested as a linear fit on a log-log plot, with an increasingly negative slope over time (Fig 2c,d). Some features of this relationship were seemingly due to changes in recipient populations—e.g. S_cross_ eventually died off when paired with E, likely raising the ratio of donor bacteria to recipient yeast (labeled “D:R” in Figure) and lowering the IDC (blue dots, Figure 2c). However, if yeast density changes were the only cause of the inverse relationship in Fig 2c, we’d have expected the IDC-per-recipient frequency to remain constant, but we found that it increased over time (SF4), suggesting that density of *both* strains were required for this inverse relationship. These findings suggest that, despite the lack of stable crossfeeding at 0% LW, we can still control IDC by tuning populations, since steady-state ratios of donors-to-recipients are inversely correlated to IDC counts.

### Mannoprotein-based cell adhesion mediates IDC and affects bacterial commensalism

Since IDC depends on cell-cell collisions in culture, we explored how known adherence mechanisms between *E. coli* and *S. cerevisiae* affect IDC. Mannoproteins are ubiquitous in fungal cell walls^57^, and type I fimbriae in *E. coli* bind to these^58,59^, forming bacteria-yeast “clumps” that can affect crossfeeding dynamics^50^. We thus repeated our batch culture experiments for population dynamics and IDC with- and without mannose added to the media, which saturates bacterial mannose receptors and reduces clumping. These cultures were measured dynamically for fluorescence as per previous experiments (Fig 1b), but here were also imaged via fluorescence microscopy to optically assess the extent of clumping.

Fluorescence microscopy analysis replicated previous findings^50^ showing that adding mannose to growth media prevented most bacteria-yeast clumping (Fig 3a). Image analysis demonstrated that the size of yeast clumps—a proxy for number of yeast cells per clump—increased concurrent with the number of bacteria in a clump (“coincident bacteria”), implying that bacteria mediate cell clump formation (SF5, SF6). Interestingly, we found that mannose-infused media prevented nearly all IDC, with only a few samples yielding single-digit IDC counts by the end of a six-day time course, roughly 10-fold fewer than corresponding samples without mannose (Fig 3b). Moreover, mannose-supplemented samples showed fundamentally altered dynamics for E_cross_-S pairings, with auxotrophic E_cross_ cells unable to survive at 0% leucine, and with much lower growth at higher percentages of leucine relative to mannose-free samples (Fig 3c, SF7). Thus, mannose interrupted the commensal dynamics previously seen without mannose.

**Figure 3:**
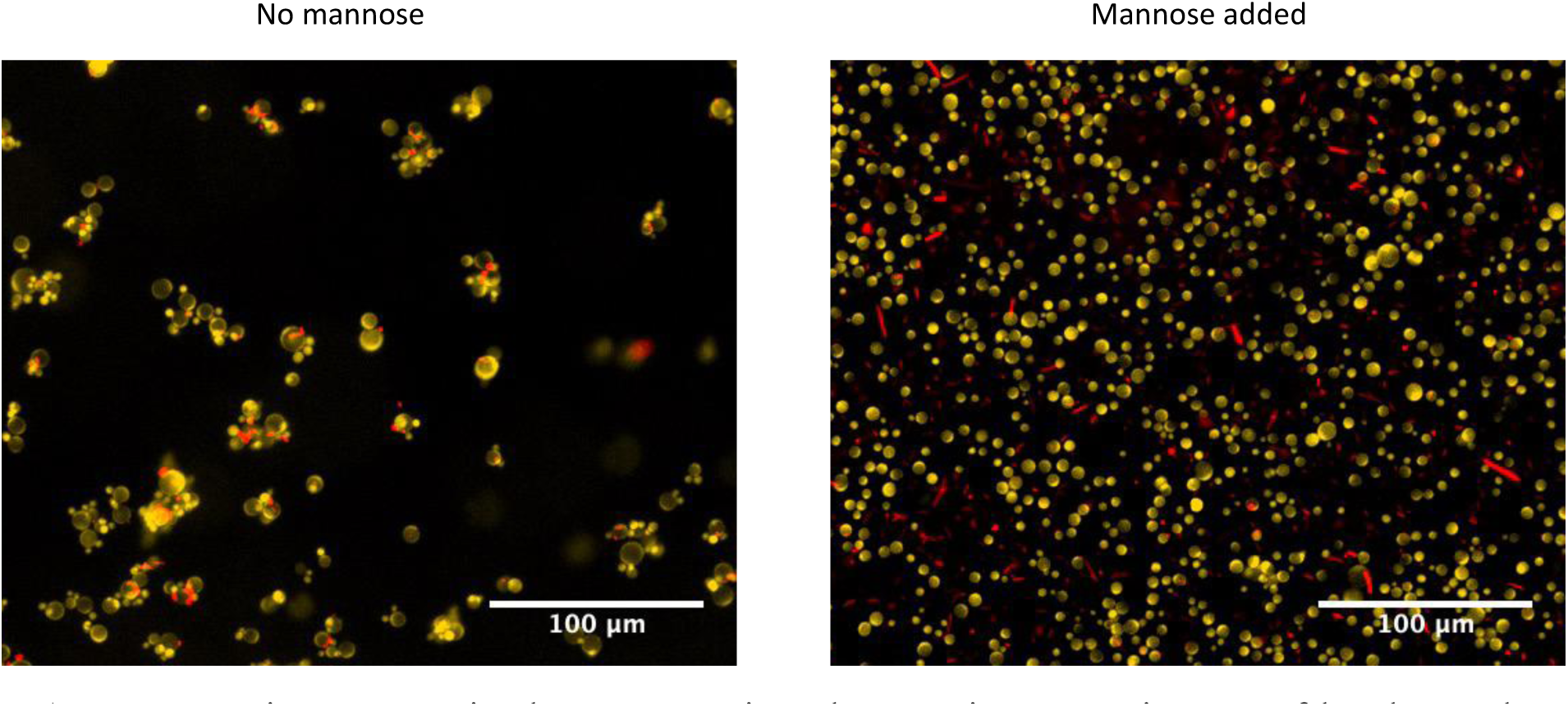
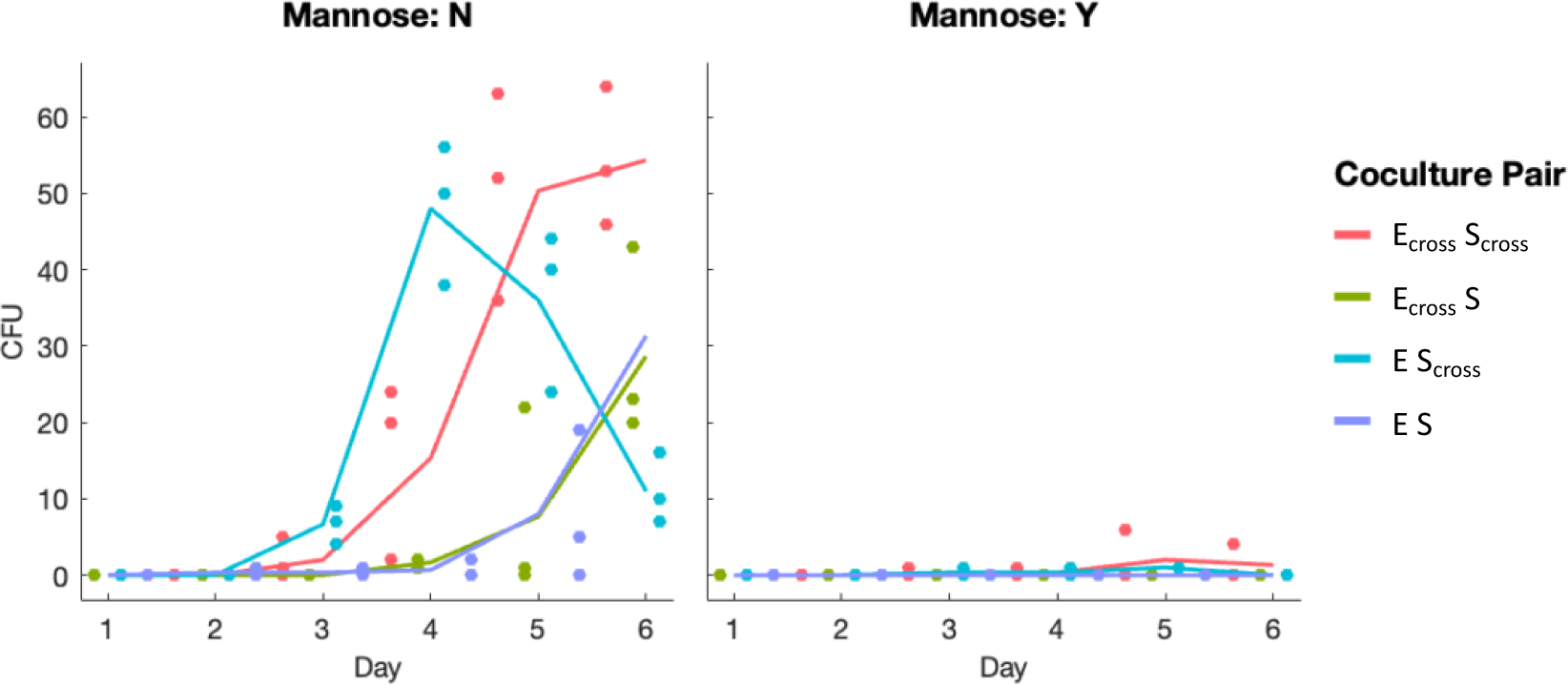
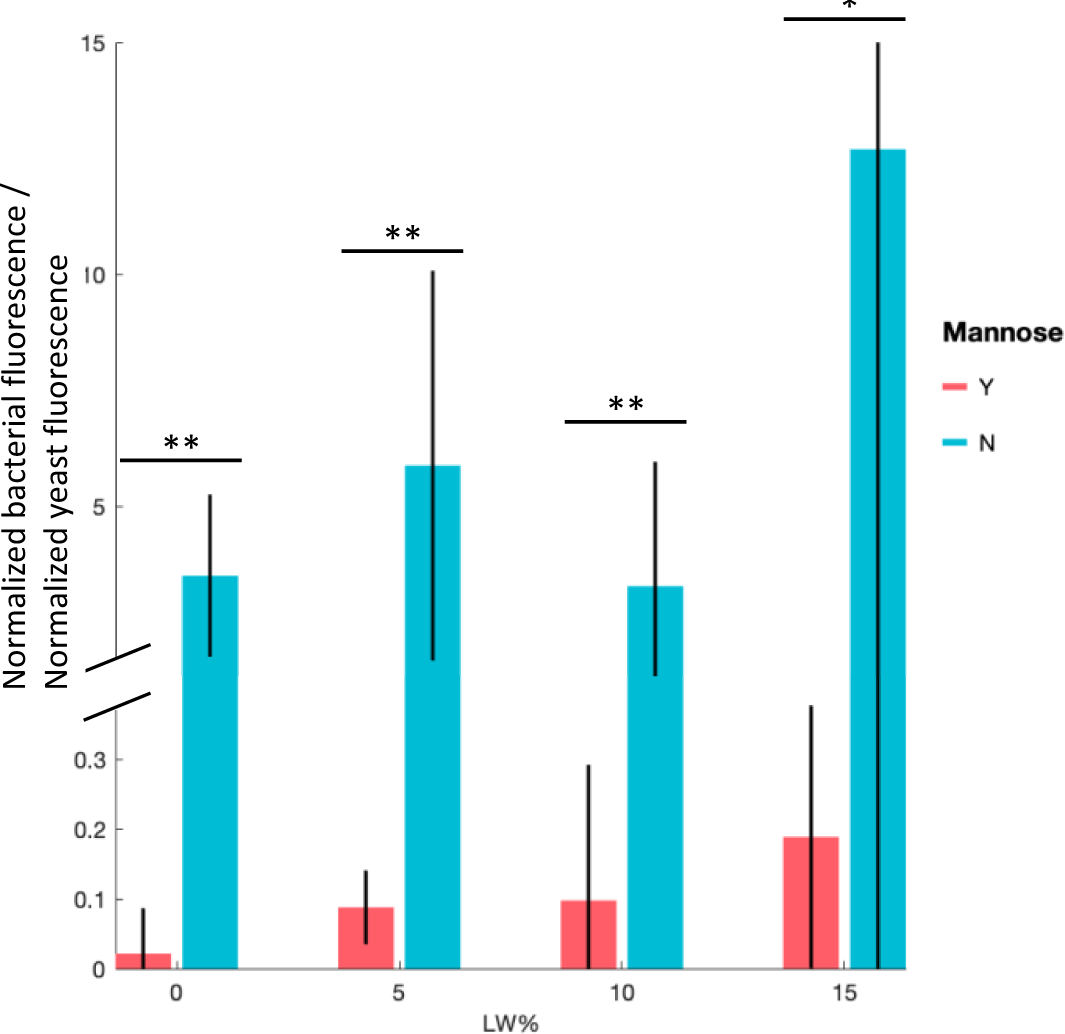
Mannose disruption of cell aggregates lowers IDC, interrupts bacterial commensalism. A. <u>Mannose interrupts mixed aggregates in culture.</u> Microscopy images of batch coculture after six days, either without (left) or with (right) mannose supplemented in media. Cells shown are *trans-*WT bacteria (E) and crossfeeding yeast (S_cross_) at 15% LW, chosen to exemplify differences in clumping. Samples were diluted 1:10, imaged with a 10x objective. Scale bar = 100 μm. B. <u>Interrupting clumps with mannose depresses IDC</u>. Raw IDC counts (CFU) for samples at 15% LW without mannose supplementation (left column) are ≥ 10x those with mannose supplementation (right column). Lines represent means of three replicates, colors, coculture pairings. C. <u>Interrupting clumps with mannose prevents commensalism for crossfeeding *E. coli*.</u> Normalized donor-to-recipient ratios for commensal E_cross_-S pairing across four different LW concentrations, after six days of batch culturing, calculated from fluorescence data. Mannose-minus (blue) and mannose-plus (red) samples show that clumping is required to sustain crossfeeding bacteria with WT yeast, especially at lower LW%. Bars represent mean of three replicates, error bars 95% CI, significance via two-sample *t* test. For 0% LW: *p* = 0.0010, *t* = 8.543, df = 4. 5% LW: *p* = 0.0041, *t* = 5.9229. 10% LW: *p* = 0.0071, *t* = 5.0709. 15% LW: *p* = 0.018, *t* = 3.8773. Degrees of freedom = 4 for all %LW.

### Deterministic models reveal boundaries of population control of IDC

To explore how the “knobs” of our system could be tuned to best affect population ratios and IDC, and to better understand the differences between clumping and non-clumping populations, we used a set of ordinary differential equations (ODEs) to deterministically model our experimental conditions (see SI discussion for more details), based on previous work modeling crossfeeding cocultures^60,61^. Moreover, to the best of our knowledge, no IDC rates have been reported in terms of reaction-diffusion equations for conjugative transfer first developed by Levin et. al.^62^, nor have any such models accounted for spatial heterogeneity in culture, so we sought to add to the body of literature on both of these fronts. We first fit the results from mannose-supplemented experiments to a system of two ODEs representing total bacteria and total yeast (including transconjugants). We ran Latin Hypercube Sampling iteratively to randomly sample all parameters within a predicted range and calculated total error between model outcomes and fluorescence data for bacteria and yeast. We then used this error to rank model parameters, which we adjusted and reran until key experimental results were demonstrated for each cell pairing: namely, susceptibility to amino acid supplementation, steady-state survival, and approximate donor-to-recipient ratio (see SI discussion, SF8-9).

Once we found best-fit approximations of parameters in bacterial and yeast growth equations, we tested a wide range of IDC rates (*γ*) against data from mannose-supplemented experiments, to determine order of magnitude for IDC given random cell collisions in culture. Though this plasmid transfer term has been previously approximated at 4*10^-3^ using a similar model for enteric bovine *E. coli*^63^, we found that γ would have to be significantly lower, roughly between 7*10^-6^ and 4*10^-^ _5_, to recapitulate our results in media containing mannose (Fig 4a, SF10).

**Figure 4:**
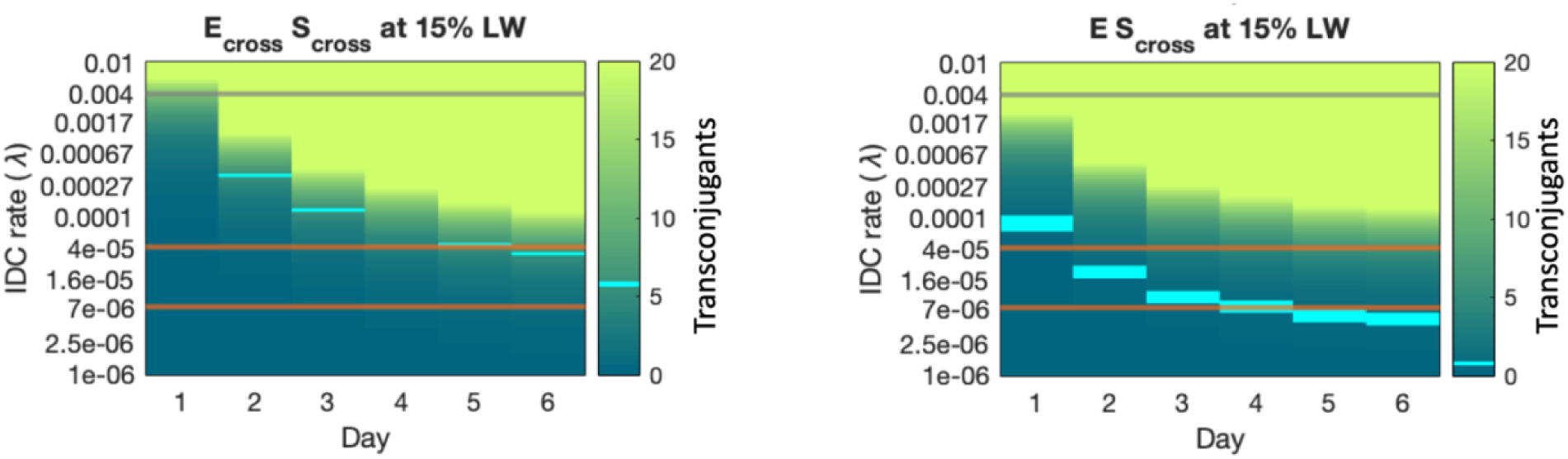
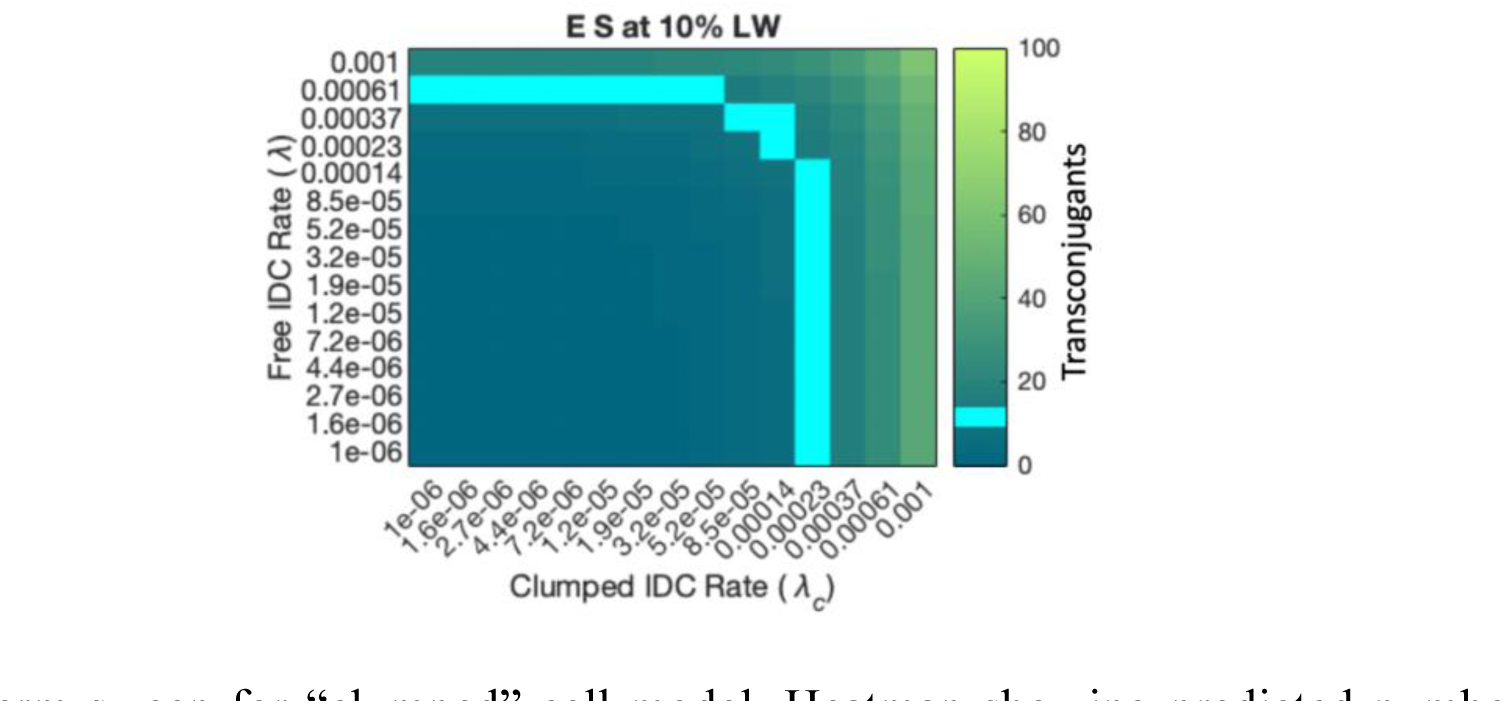
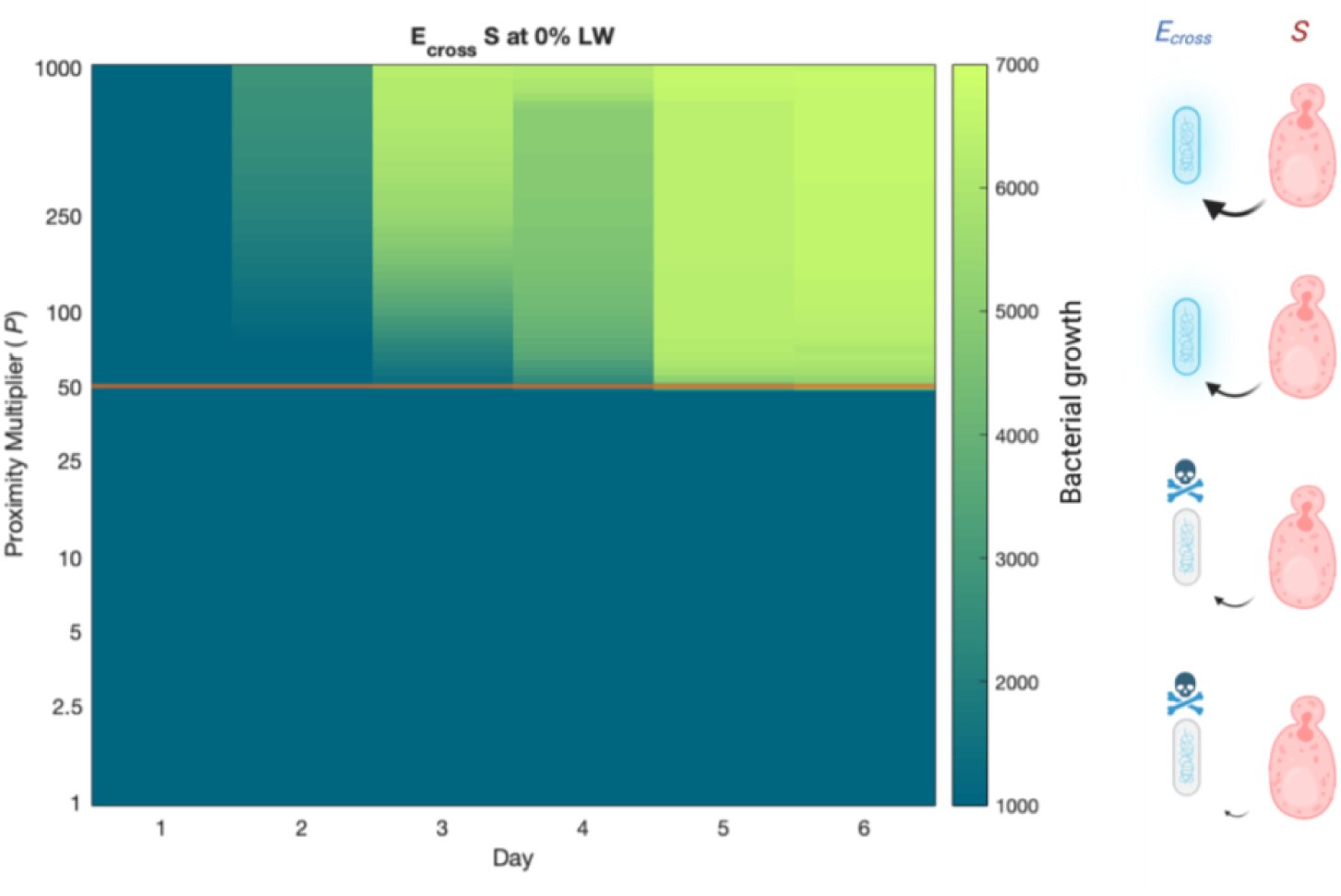
Deterministic modeling shows limits of IDC rate terms, proximity benefits, in and out of aggregates. A. <u>IDC rate-term sweep for “free” cell model.</u> Heatmaps showing predicted number of transconjugants (heatmap) for a range of IDC rates γ (y-axis) over six days of batch culturing, assuming cells are unable to clump, and thus conjugate via random collisions. Two of the four experimental conditions that yielded IDC counts > 0 with mannose are shown (mean experimental CFU of three replicates = cyan heat markers). Gray line at γ=0.004 represents literature prediction for enteric *E. coli* conjugation rate^63^, orange lines represent range of IDC-rate values matching experimental data, roughly between 7*10^-6^ and 4*10^-5^. B. <u>IDC rate-term sweep for “clumped” cell model.</u> Heatmap showing predicted number of transconjugants (color) for a range of “free” IDC rates γ (y-axis) and “clumped” IDC rates *γ_c_* (x-axis), for one representative experimental condition at day six of batch culturing. Cyan boxes represent mean experimental IDC count of 18.7 per 100 uL culture (three replicates),, with several values of *γ* and *γ_c_* resulting in this number of transconjugants. Two main conditions yield the experimental IDC results: low *γ_c_* with *γ* above 5*10^-4^, or low *γ* with *γ_c_* near 3*10^-4^. Because free-model results showed *γ* below 5*10^-5^, it’s likely that the latter case is true, with most conjugation resulting from clumped interactions. C. <u>Proximity term sweep for “clumped” model.</u> Heatmap showing predicted bacterial fluorescence signal (color) for range of proximity-benefit multiplier *P* (y-axis) over six days, for E_cross_-S pairing at 0% LW. Orange bar shows approximate *P* value matching experimental coculture data, i.e. a *P* value high enough (∼50) to allow E_cross_ growth solely from clumping to WT yeast. Based on the model structure, this implies that nutrient-dependent *E. coli* see ∼50x benefit from proximity to WT yeasts in terms of nutrient access from yeasts, though other mechanisms for this commensalism are possible.

We performed another round of parameterization against measurements of clumped cells growing in mannose-free media, using ODEs modified to include clumping. In this model, IDC was represented by two different rates: *γ* for free-cell collisions, as per previous fits, and *γ_c_* for clumped cells, and. The model fits (SF11) yielded two possibilities that recapitulated the data: low *γ_c_* with *γ* in the range of 5*10^-4^ – 1*10^-3^—higher than *γ* values found in the free-cell model, thus probably not representative—or low *γ* with *γ_c_* in the range of 2*10^-4^ – 4*10^-4^ (Fig 4b, SF12), only ∼10-fold lower than the conjugation rate predicted for intraspecies transfer. Together, these results predict that conjugation transfer rates for IDC in reaction-diffusion type models are near those found for intraspecies transfer, but only for cells that are clumped.

Additionally, a proximity term *P* was used in this model to account for changes in benefit arising from the proximity of clumped cells, which allows for E_cross_ survival with WT yeasts (S). *P* value sweeps showed an apparent amino-acid secretion increase on the order of 50x from WT yeasts (S), to allow E_cross_ cells to grow in 0% leucine (Fig 4c). While *P* mathematically served to multiply the amino acid secretion term in the model, it could just as plausibly have resulted from leucine (intermediate) in yeast cell lysate or some other mechanism of bacterial benefit. Both explanations had caveats, however. Roughly speaking, since nutrient concentrations diffuse exponentially^49^, we might’ve expected such a 50x nutrient benefit from secretion to correspond with ∼4x proximity of bacteria to yeasts (since ln(50)∼4), but there’s no evidence that WT yeasts secrete leucine (intermediates) basally^55^. On the other hand, yeast lysate-derived nutrients cut against higher IDC counts for these samples, which suggested a high number of surviving yeasts. Moreover, this model didn’t deconvolute whether mannoprotein binding led to both higher IDC and commensalism independently, or only higher commensalism which in turn increased IDC. Still, the model served both to add to our knowledge of how key features of this system function and allowed us to predict IDC outcomes for various experimental parameters, as we explore later.

### Mixed Colonies exhibit inverse D:R to IDC relationship, low spatial intermixing

Having thus far characterized IDC in well mixed liquid cocultures, we next sought to understand how population dynamics affect IDC in spatially constrained settings, to better predict IDC functionality in natural settings such as biofilms^46^. We emulated “expansion” assays^64^, which have previously demonstrated greater intermixing of mutualistic populations^48,49^ by repeating batch culture initial conditions on 2% agar minimal media plates, except with 10-fold fewer initial cells. We pipetted ≥ 18 2μL mixed-cell droplets onto plates and allowed them to grow continuously for six days. We imaged three colonies for 2D spatial distribution each day via wide-field fluorescence microscopy, and another three that were then scraped, washed, and diluted for composition and IDC measurements (Fig 5a,b).

**Figure 5:**
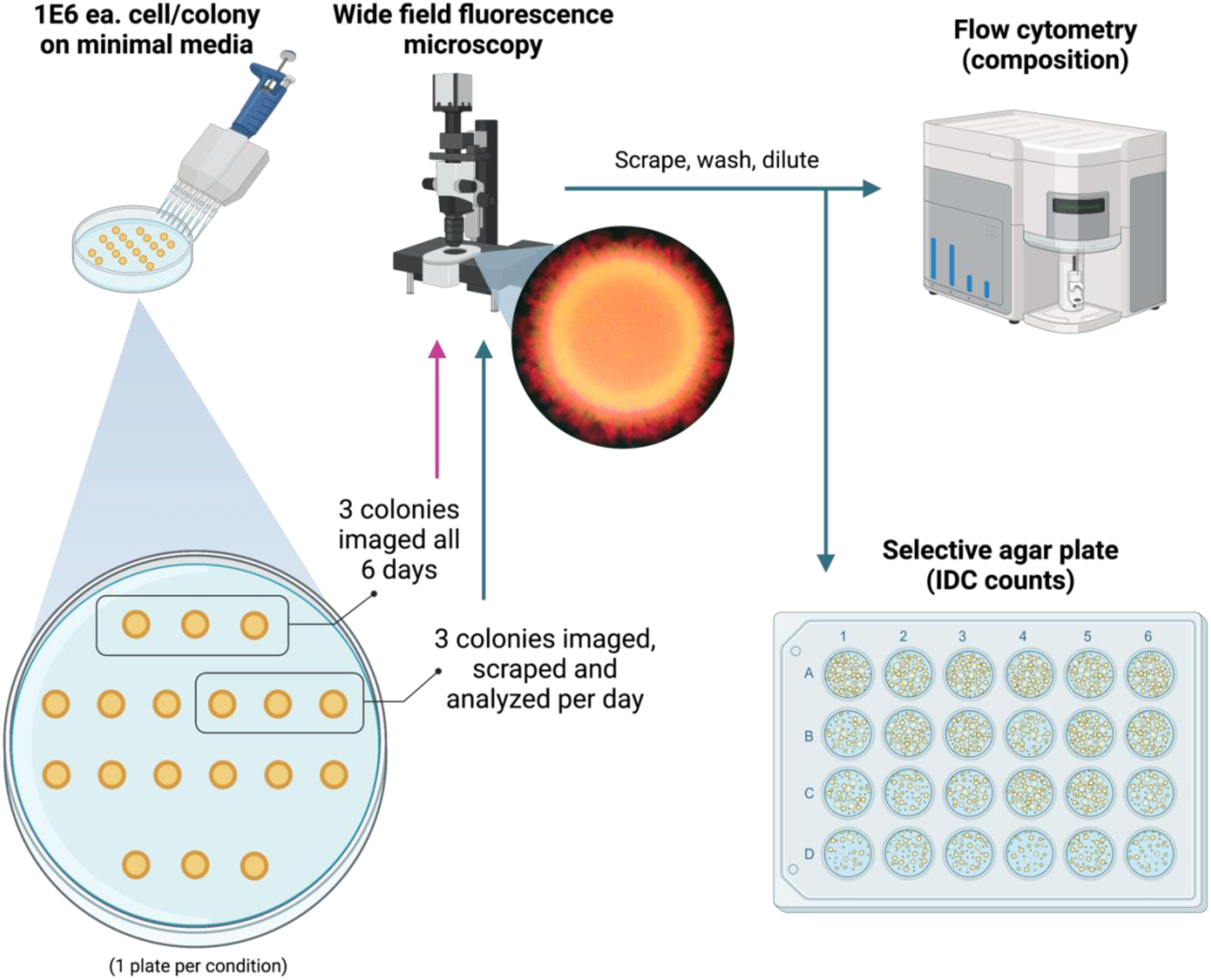
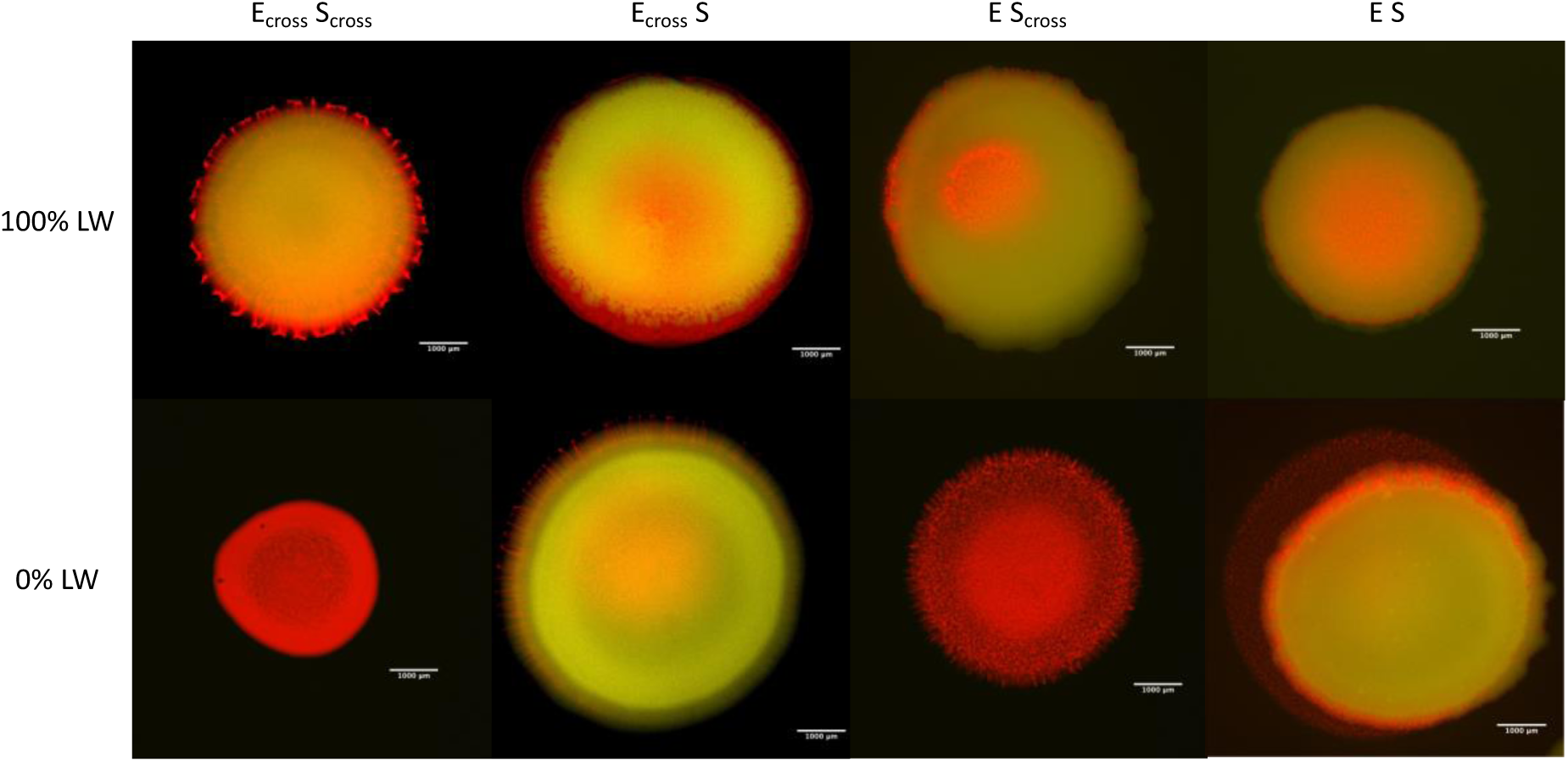
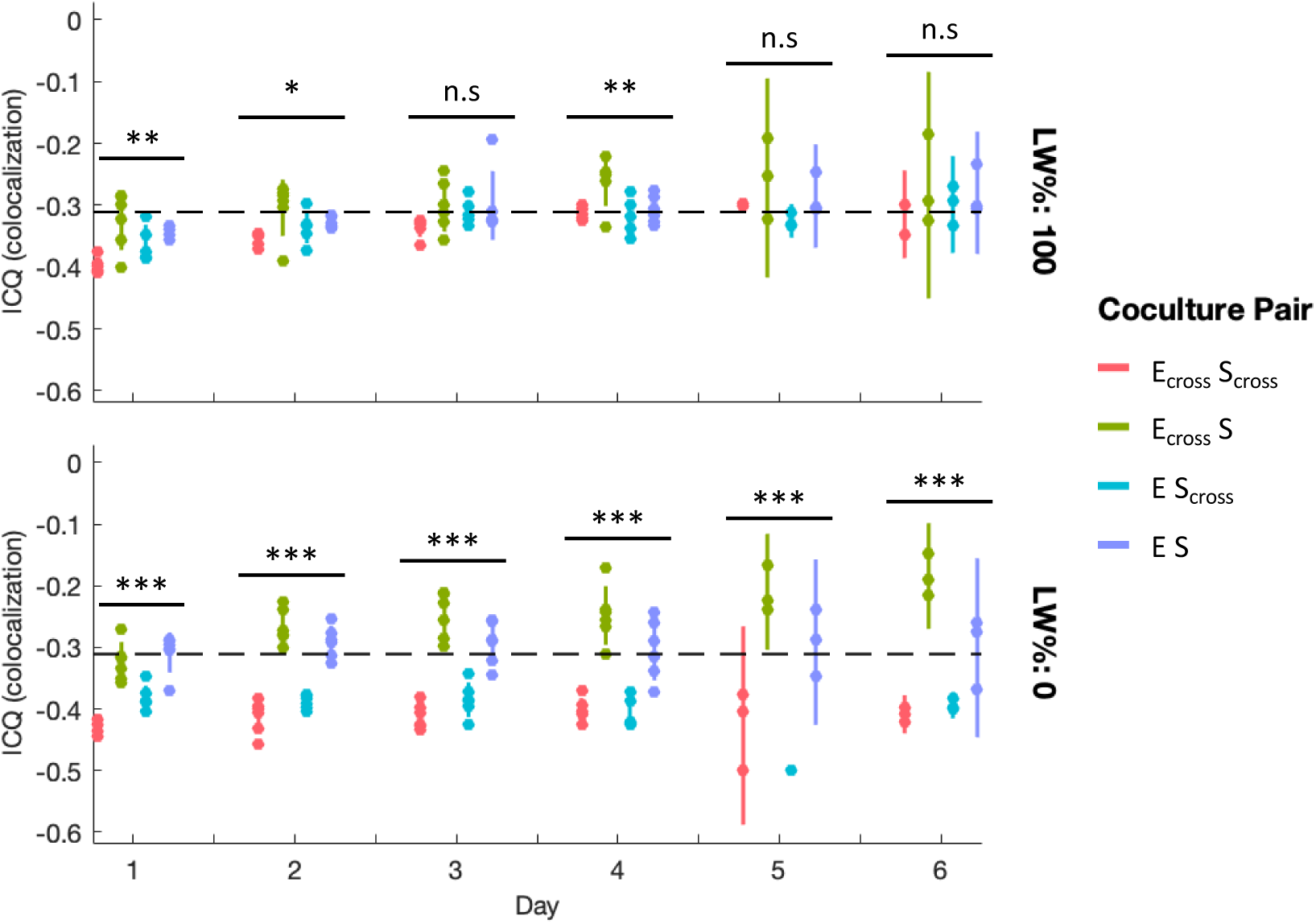
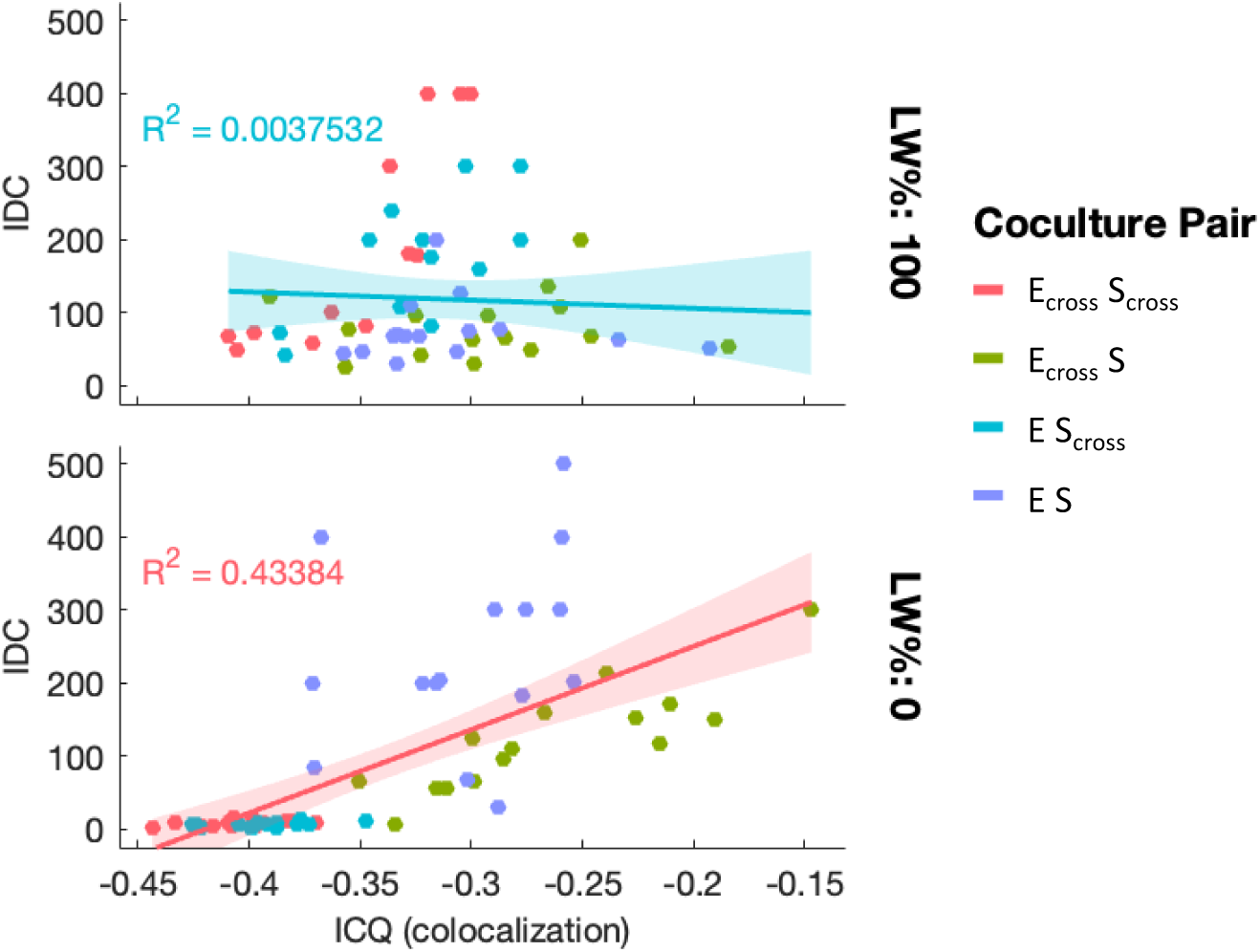
Mixed colonies follow similar dynamics to culture conditions, show increased IDC for more spatially mixed populations. A. <u>Experimental setup of colony assay</u>. Cells are combined and pipetted onto minimal media with 2% agar. Each plate contains ≥ 18 colony replicates of one cell pairing, and one amino acid concentration. After each day’s growth, 6 colonies are imaged with a wide-field fluorescence microscope. 3 of these continue to be imaged daily, while the other 3 are scraped, washed, and diluted for flow cytometry and IDC plating. B. <u>Example of mixed colony cell distribution.</u> Fluorescence microscopy images for each cell pairing at 100% LW (top row) and 0% LW (bottom row). All pairings here include *cis-* donors. Yeasts are displayed in yellow channel, bacteria in red. Channels are scaled for brightness to emphasize distribution, scale bar = 1000um. C. <u>Colocalization shows divergent intermixing at 0% amino acids, though antagonism drives</u> <u>spatial distribution overall.</u> Li colocalization analyses of colonies (ICQ = 0.5 is complete colocalization between channels, ICQ = -0.5 is complete spatial segregation) show range of outcomes for 0% LW colonies, less so for 100% LW colonies. All experimental ICQ values range from -0.1 – -0.5, implying antagonistic spatial segregation. Distributions shown are of *cis*-donor pairings. Calculated ICQ values for each replicate and condition represented by dots, 95% CI of the mean by vertical bars. Stars denote *p-*values from ANOVA 1-way test of 95% confidence between all 4 pairings at each day and % LW using sum of squares test (see Table S6 for F-values and degrees of freedom). Two-sample *t-*tests of day 6 ICQs (not displayed) show significant differences between 0% LW and 100% LW for E_cross_-S_cross_ pairing (*p =* 0.0064, *t* = -5.2247, df = 4) and E-S_cross_ pairing (*p* = 0.0073, *t* = -5.0285, df = 4), while E_cross_-S and E-S pairings do not show differences between LW% (*p* = 0.15, *t* = 1.770, df = 4; and *p* = 0.64, *t* = -0.5091, df = 4, respectively). D. <u>Colocalization correlates positively with IDC values.</u> ICQ values plotted against raw IDC (CFU) counts for *cis*-donor pairings, at 0% and 100% LW, for IDC≥2. For the smaller range of ICQ values at 100% LW, IDC counts slow little divergence, whereas at 0% LW, IDC correlates positively with ICQ.

As with batch cultures, there was an inverse correlation between donor-recipient ratios and IDC in most cases, though with greater noise (SF3). However, IDC-per-recipient rates remained relatively constant, unlike cultures (SF4). These differences from culture conditions might be due to “jackpot” populations, in which a genetic island of transconjugants finds a spatial niche among the stochastic colony front^65^, resulting in a wider range of IDC counts for each condition (see SI discussion, SF14). Because conjugation has been shown to occur along population boundaries^47^, we determined relative population mixing by calculating colocalization^66^ of bacterial and yeast fluorescence signals (see SI discussion). While colocalization did positively correlate with overall IDC values, most mixed colonies had very low colocalization, suggesting once more that antagonism dominates population dynamics between these species (Fig 5c,d).

### Population dynamics can be tuned to rescue a recipient population through IDC

To test whether population-control of IDC can be used to alter recipients at population scale (we know it can alter individual recipients), we next sought to “rescue” starved yeast cells with poor or non-existent growth, via genes carried on the transferred DNA. We first tested this with the *cis*-IDC plasmid pTA-Mob 2.0, which carries *HIS3* and *URA3* and allows transconjugants to grow in media deficient for uracil and histidine. IDC from WT donors mostly failed to rescue U or H-auxotrophic yeast recipients growing in low concentrations of uracil and histidine (% UH), as the bacteria competed the yeasts to collapse before sufficient transconjugant growth could establish (SF15).

Our previous results showed higher IDC for lower donor-to-recipient ratios, so to increase the likelihood of rescue, we used auxotrophic bacterial donors at 0% leucine. These donors can thus only survive if the paired yeasts metabolically support them. Remarkably, we found a drastic increase in IDC-rescue from E_cross_ donors, for both S_cross_ and S recipients, an effect that varied by uracil and histidine amounts (Fig 6a,b, SF15). E_cross_ rescued both recipient strains with greater speed and efficiency than E did in all cases, though at 0% UH, paired crossfeeder populations collapsed (Fig 6b). At intermediate concentrations of uracil and histidine—especially 5% UH— rescue showed high stochasticity, as some biological replicates were fully rescued while others collapsed (SF16). We also used our clumping model to predict the range of possible rescue outcomes for each cell pairing over a range of amino acid concentrations. With minimal alterations to account for experimental differences, the model recapitulated our experimental results: for most concentrations of U and H, and with [L] kept low, bacterial antagonism is minimized, and greater IDC is possible, allowing for the increased rescue of yeast seen in these experiments (Fig 6b,c, see SI discussion for model changes).

**Figure 6:**
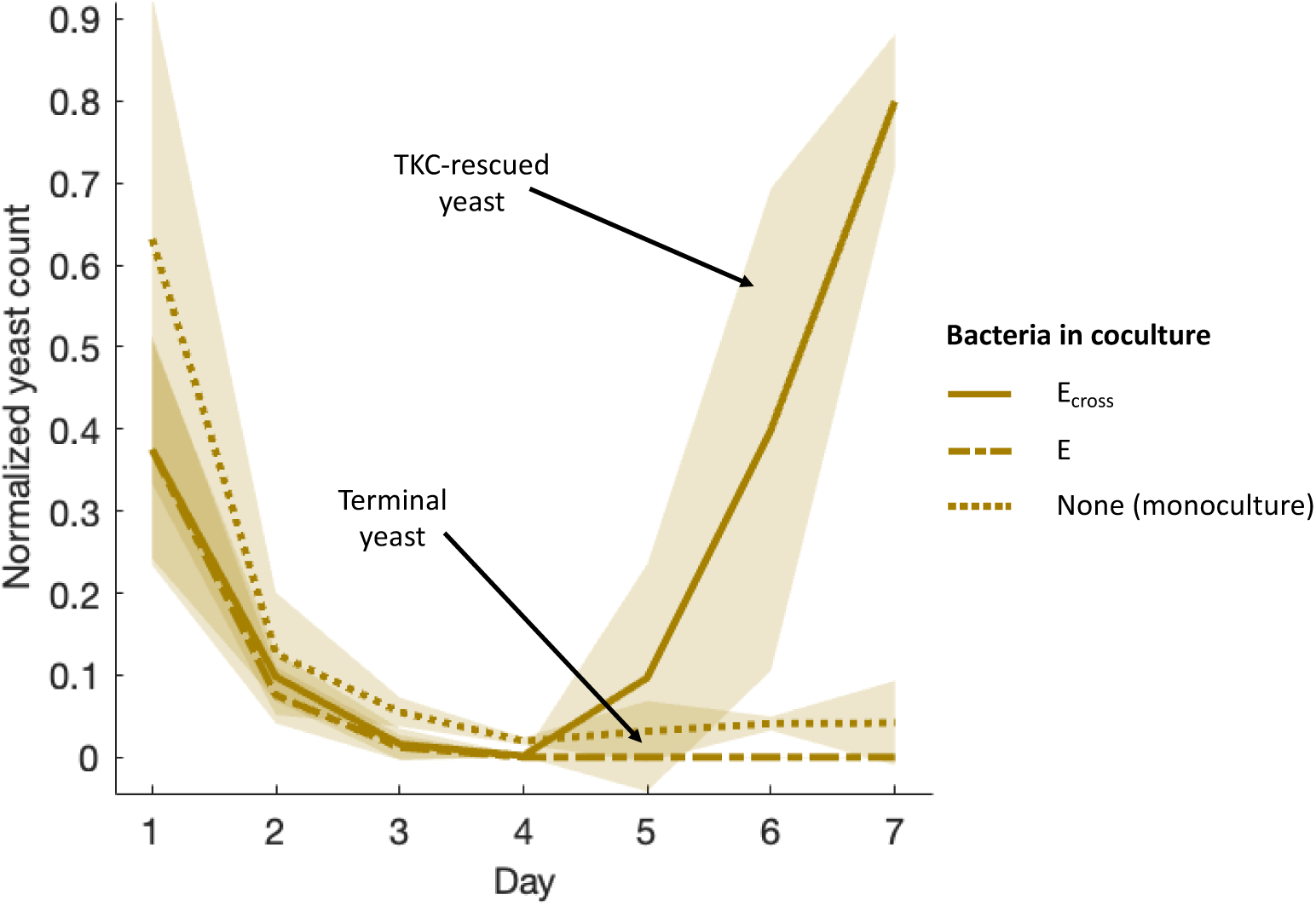
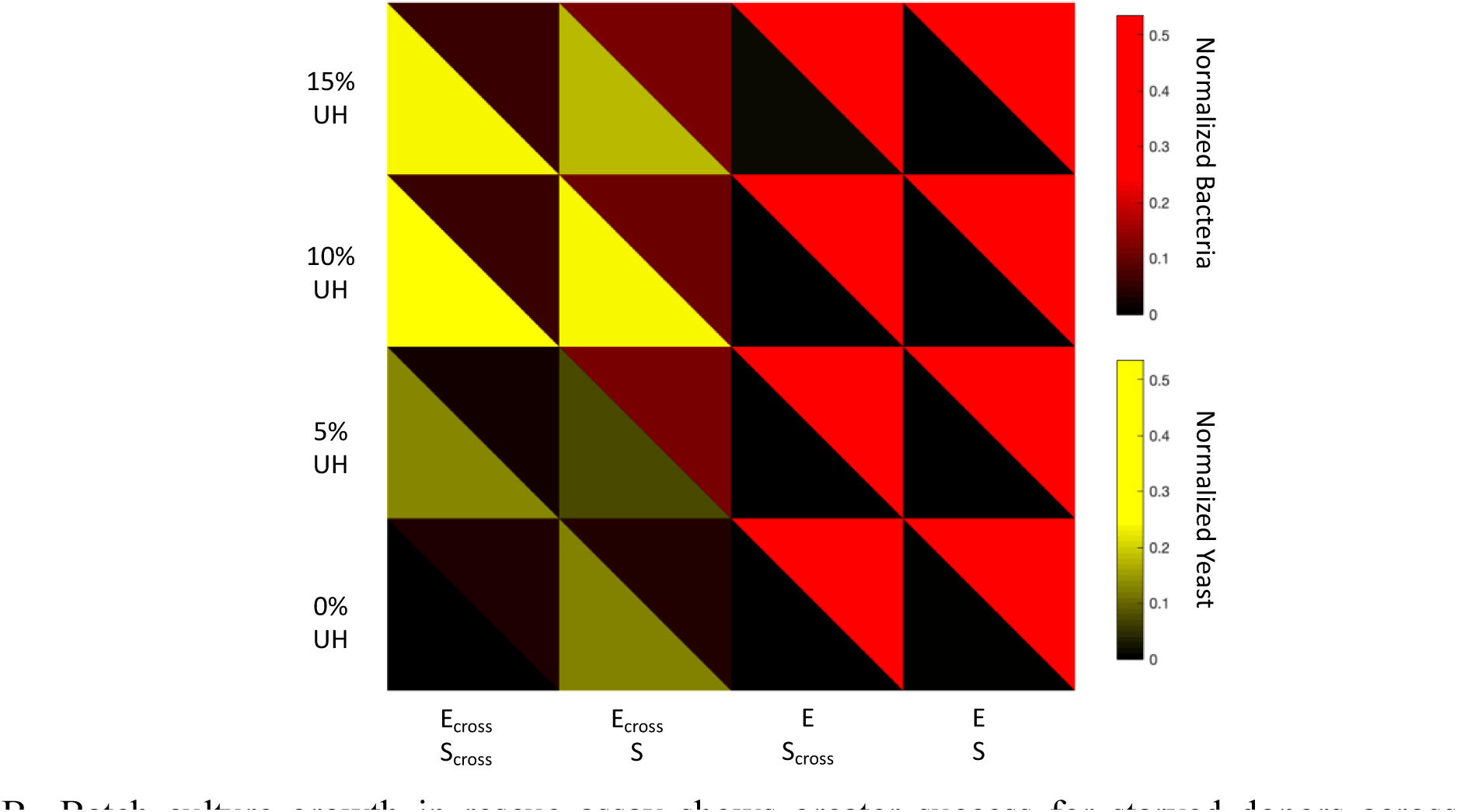
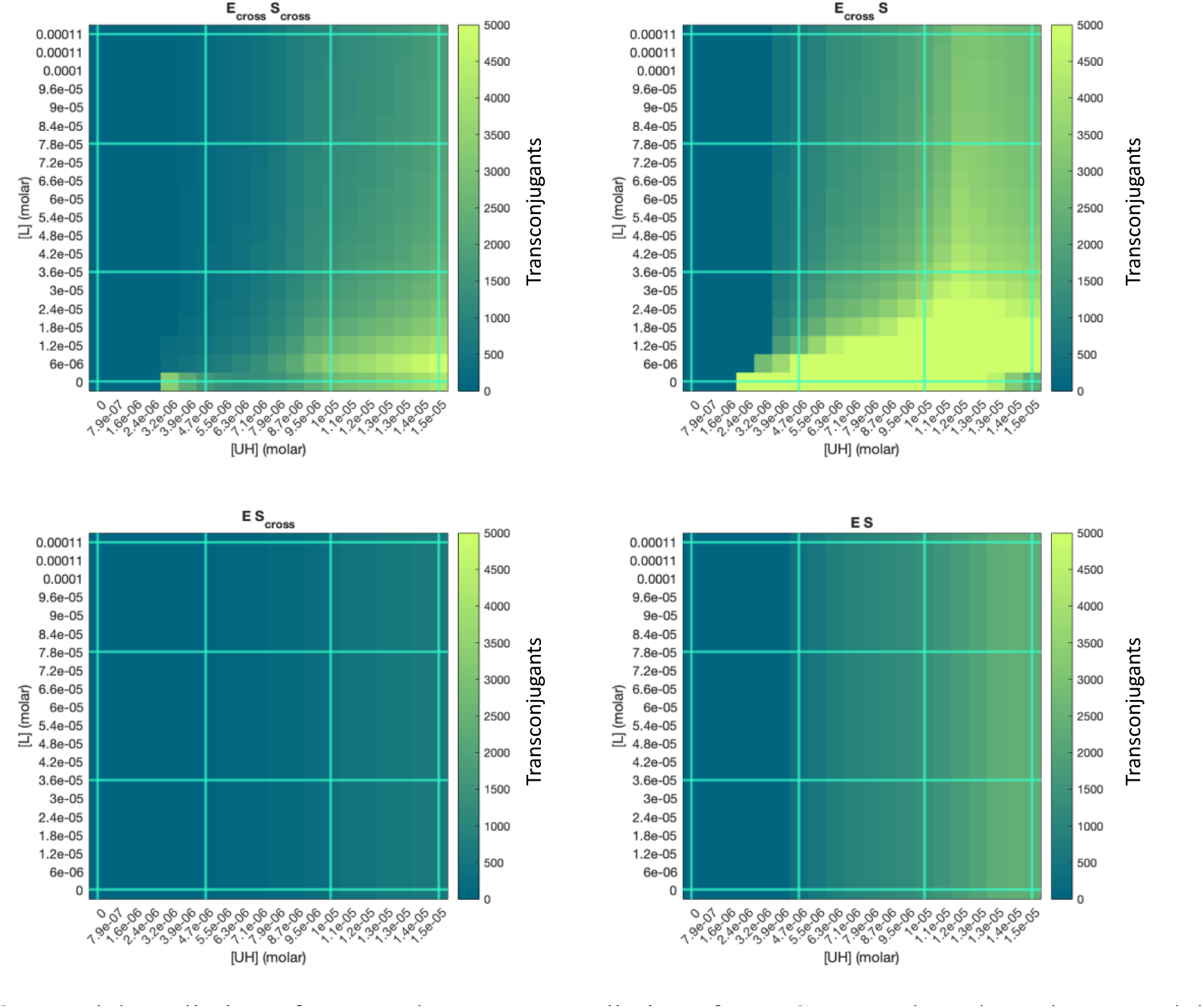
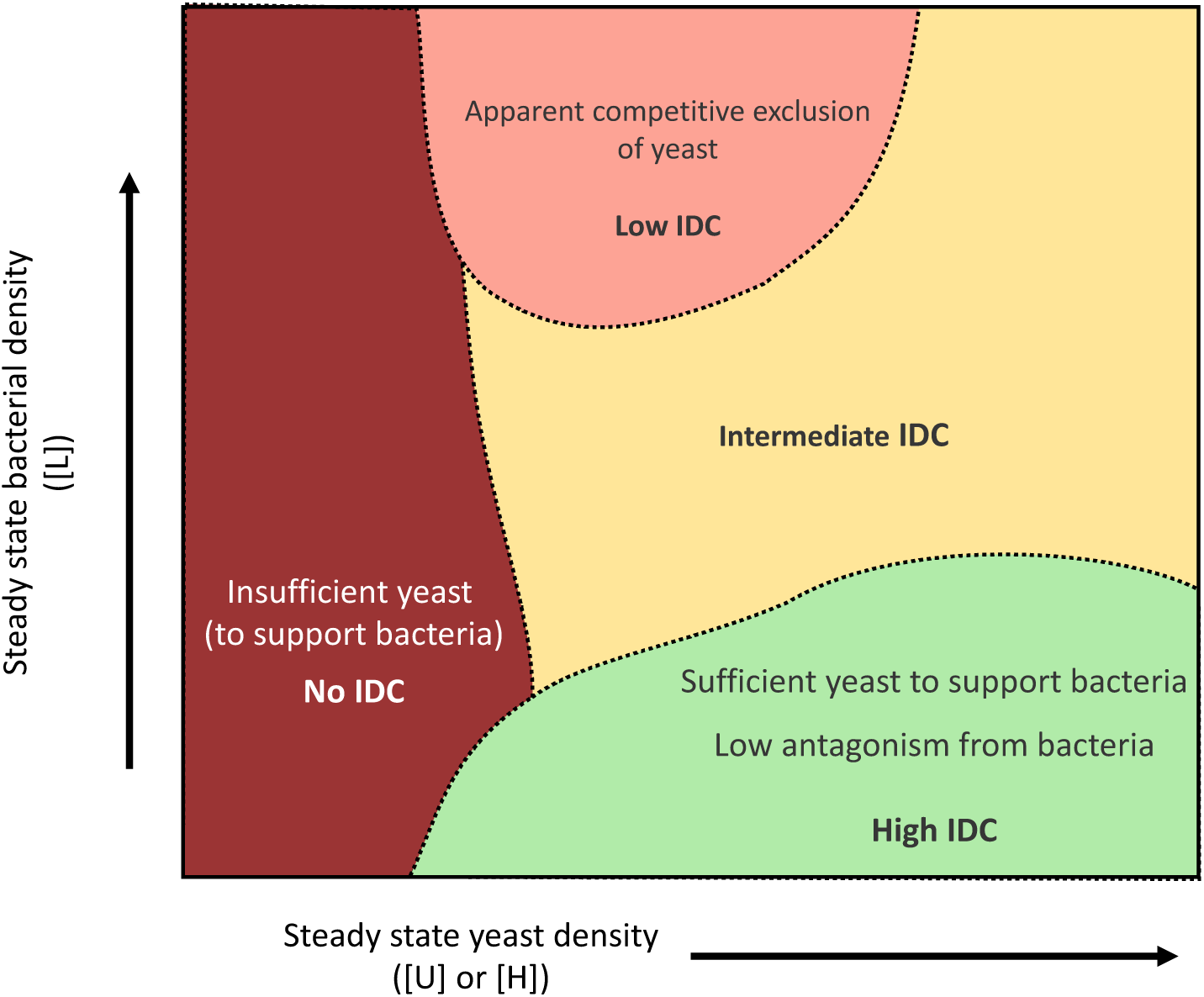
Utilizing population dynamics allows IDC-mediated? rescue of unhealthy recipient populations. A. <u>Yeast growth is rescued by crossfeeding donor IDC</u>. Yeast cell counts from flow cytometry, normalized to max count after day 1, for each cell pairing of S_cross_ at 10% UH, 0% L. In monoculture (dotted line), S_cross_ grows poorly at 10% UH. WT bacterial donors antagonistically depress S_cross_ growth (dot-dash line), despite their ability to transfer IDC plasmid that would rescue recipients. E_cross_ donors, on the other hand, are able to transfer sufficient rescuing plasmid (solid line), allowing full S_cross_ rescue. Means of six replicates over two experiments shown as traces, shading as standard deviation. B. <u>Batch culture growth in rescue assay shows greater success for starved donors across</u> <u>several conditions</u>. Split heatmaps of normalized cell counts from flow cytometry for four cell pairings (columns) and four concentrations of uracil and histidine (rows). All samples grown with 0% leucine to starve E_cross_ (bacteria in red). Yeast (yellow) auxotrophic for URA3 or HIS3, show greater growth upon receiving conjugated pTA-Mob 2.0 (*cis*), which is only significant when paired with E_cross_. Brightness is mean of six replicates across two experiments, normalized to max cell count per species and experiment, multiplied uniformly to visualize low-growing strains. C. <u>Model prediction of rescue phase map.</u> Predictions for IDC counts based on clump model, adapted for rescue assay conditions. Concentrations of L (y-axis), U, and H (x-axis), for each cell pairing in rescue assay shown, with values 0%, 5%, 10% and 15% highlighted with cyan lines. Note that while experimental rescue conditions don’t include a range of leucine concentrations (only 0% L), the model predicts a range of [L] over which E_cross_ could rescue yeast more effectively than WT bacteria (E). D. <u>Conceptual phase map of IDC outcomes.</u> Comparing growth of bacteria (y-axis) and yeast (x-axis), as controlled in rescue assay by amino acid levels. At low enough growth for both species, populations collapse before sufficient IDC can occur. When bacteria are sufficiently supplied with nutrients (or aren’t dependent on them), antagonism suppresses yeast growth, limiting rescue capacity. At low bacterial fitness, but moderately low yeast fitness, enough yeast cells are present to sustain growth of the starved bacteria for long enough to allow IDC, and the lack of antagonism from E_cross_ donors allows for optimal rescue of recipient population.

### IDC-mediated CRISPR killing can be interrupted by mannose addition

We next tested whether we could collapse or depress a recipient yeast population via IDC-mediated killing. We designed a conjugatable CRISPR/Cas9 system that can be transferred from bacteria to yeast, where it targets a blue fluorescent, *URA3*-carrying plasmid in recipients, such that destruction of this plasmid would render recipient cells unable to grow in uracil deficient media. Unlike most Cas9 systems, which utilize a repair sequence to replace the cut DNA, we relied on repeated cutting of the target DNA with no repair, since our goal was simply to suppress the target cells’ growth. Targeting an episomal sequence was also essential, to discern both IDC rates and cutting efficiency separately, without the lethality of cutting genomic DNA in recipient yeast (Fig 7a). After verifying that the IDC-Cas9 plasmid is efficient for cutting its target via both direct yeast transformation and IDC (Fig S17), we batch cultured crossfeeding yeast (W auxotrophs) containing the *BFP-URA3* plasmid at low levels of tryptophan and 0% uracil, along with donor cells that either contained a functional IDC-Cas9 system, or one lacking an *ori^T^* sequence and thus unable to transfer DNA. At 1% W, all yeast cultures died out, while at higher levels of W (5% and 10%), bacterial antagonism resulted in depressed yeast levels relative to monoculture yeast growth (Fig 7b, S18). Donors carrying IDC-Cas9 (“cutters”) significantly depressed yeast growth beyond antagonism-based decreases, especially at 5% W, where yeast growth was decreased several-fold beyond non-transferring control donors (Fig 7c).

**Figure 7:**
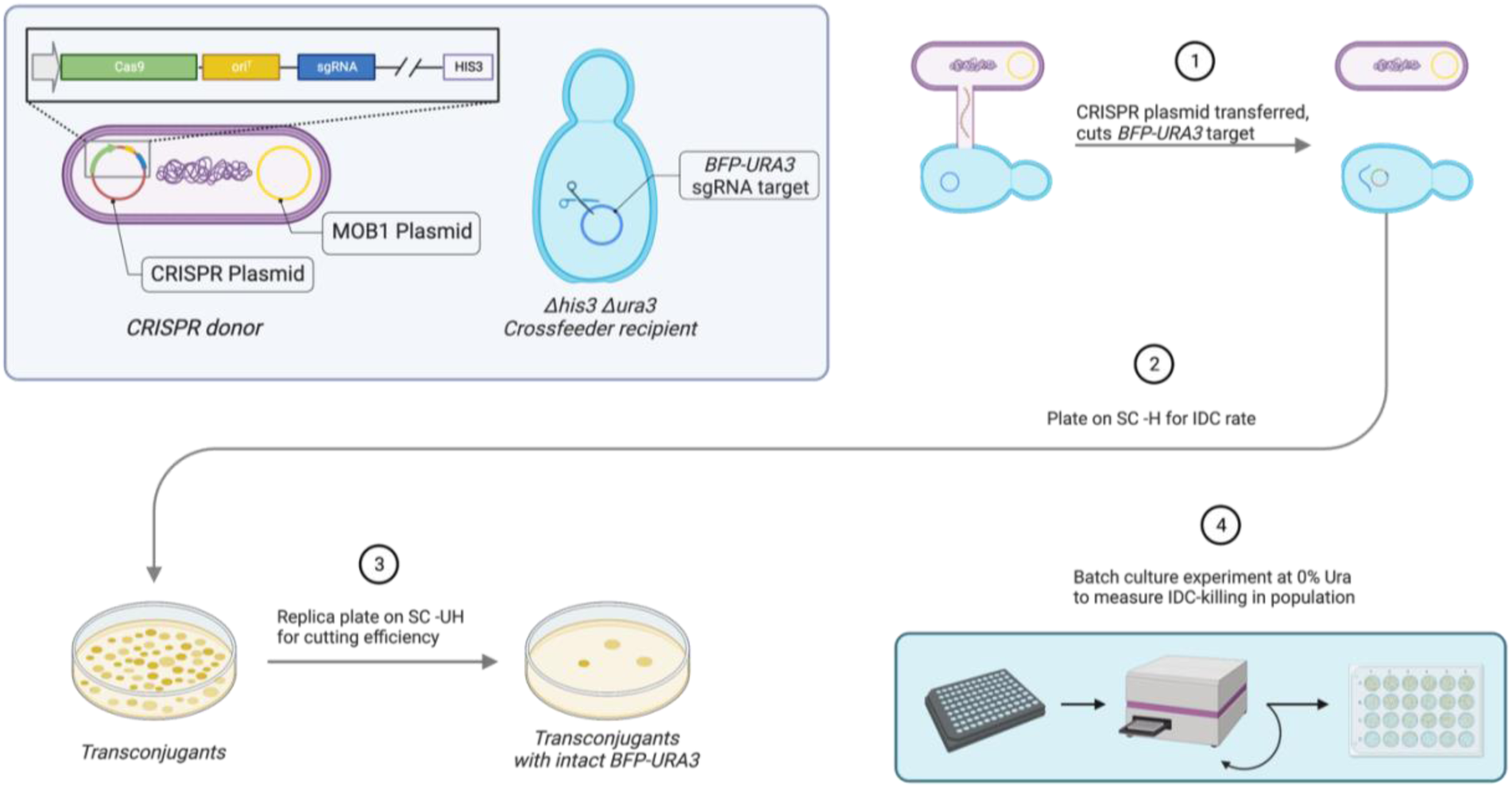
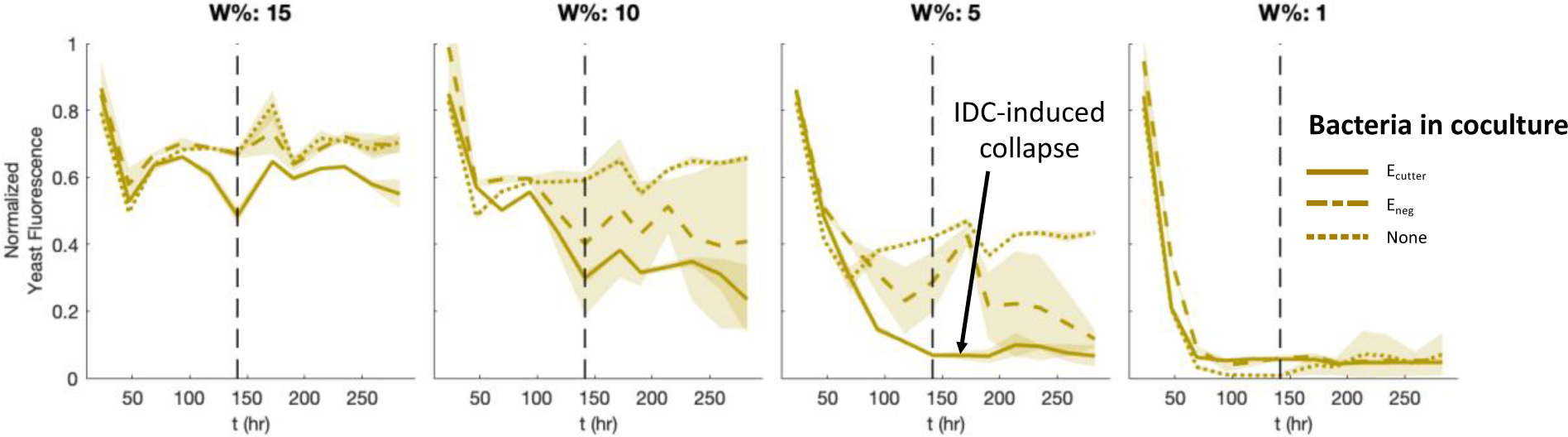
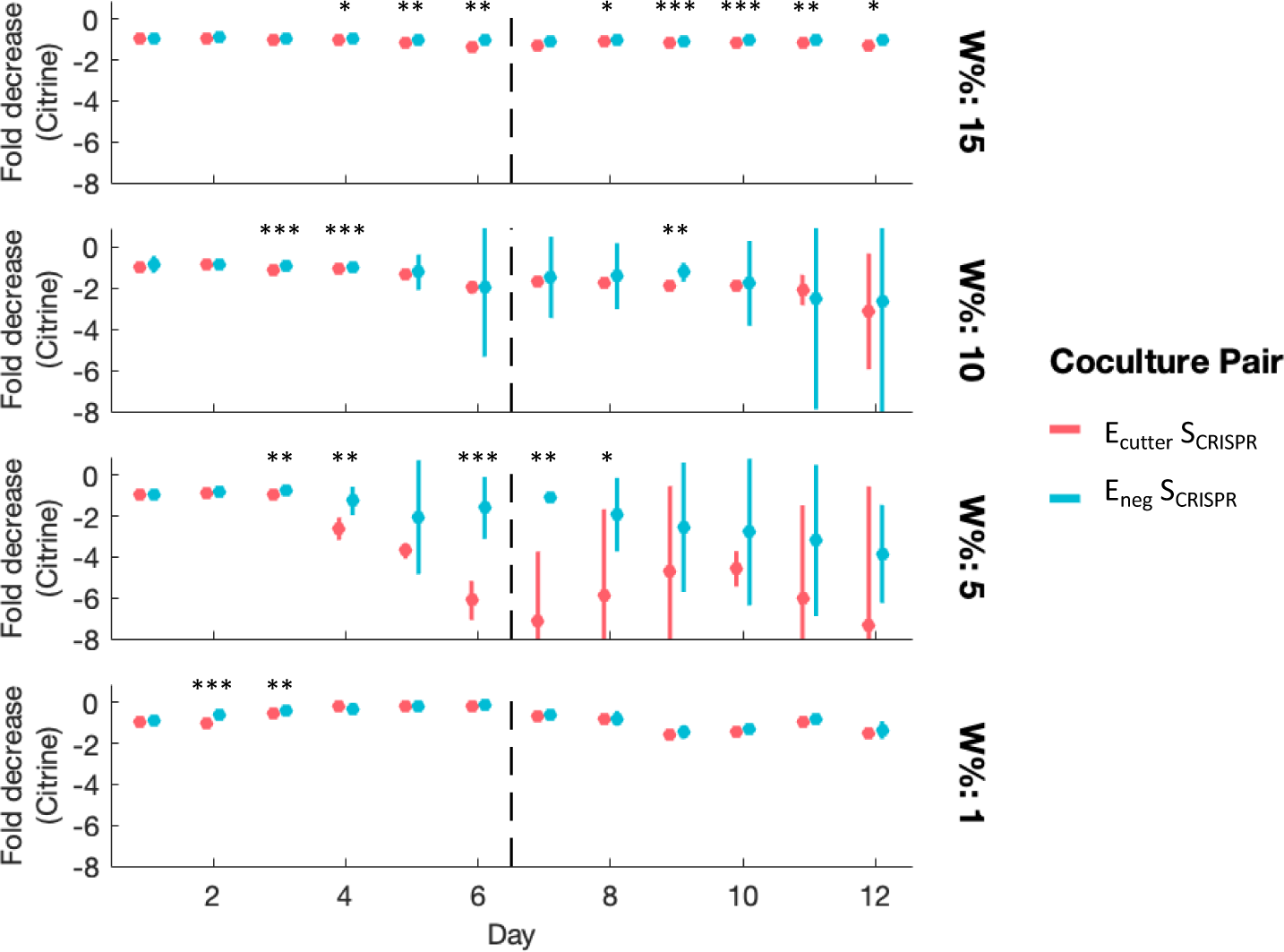
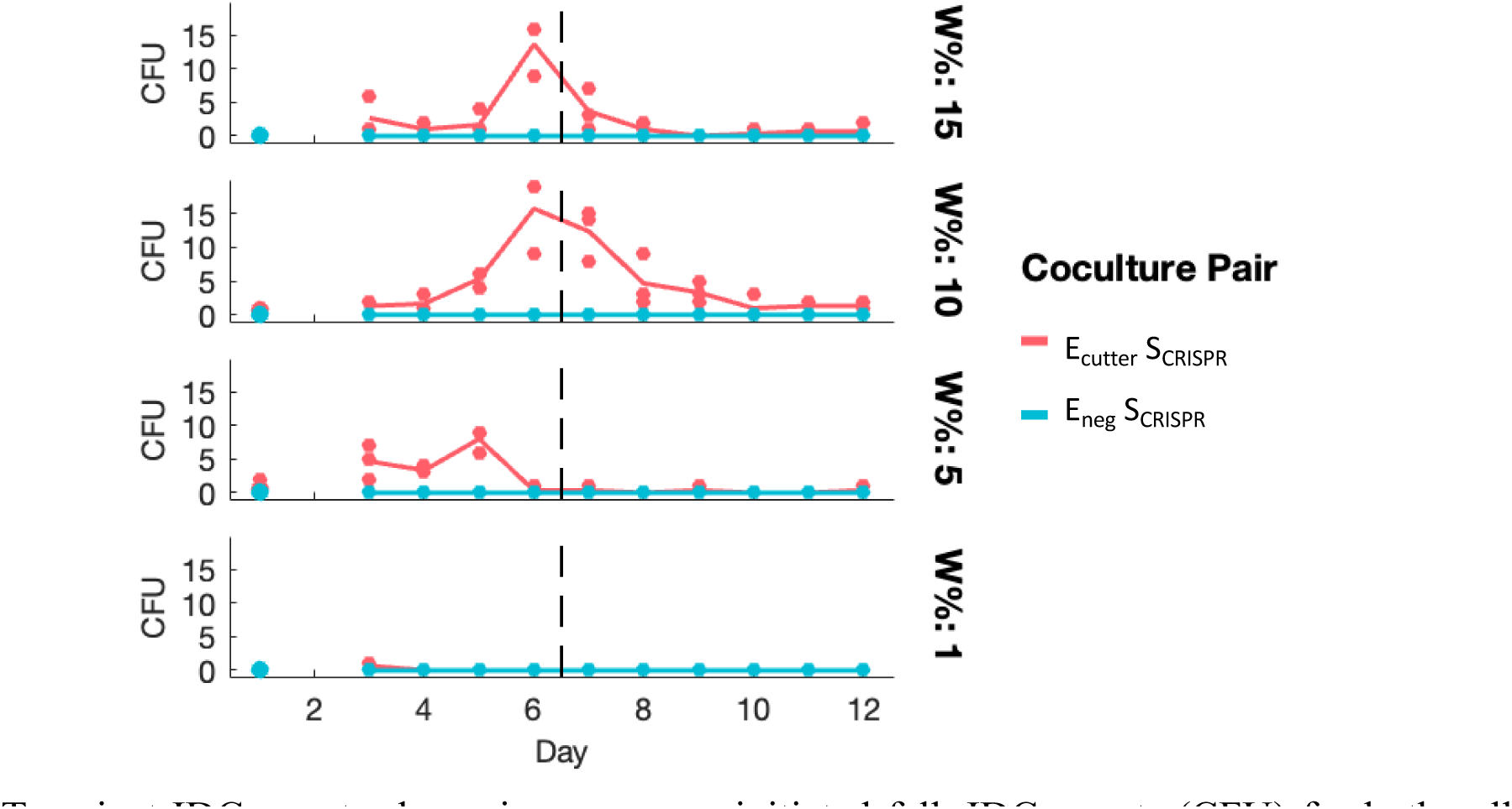
IDC-mediated CRISPR killing is able to drive recipient population extinction, is mannose-interruptible. A. <u>Design of IDC-mediated CRISPR system.</u> pTA-Mob 1.0 T4SS plasmid (*trans*) is paired with a Cas9 plasmid that contains the *ori^T^* sequence (allowing for transfer), HIS3 yeast selection marker, and sgRNA coding for a connector region in BFP-URA plasmid. Recipient yeast are *Δura30, Δhis3::HPHMX6* and carry BFP-URA plasmid. Upon IDC transfer, BFP-URA plasmid is cut via Cas9, with no repair template, but yeast can continue to grow in media supplemented with uracil. In this way, we can measure IDC efficiency independently from CRISPR cutting efficiency, by plating for IDC (SC -H) and then replica plating for cut yeast (SC -UH). Finally, cut-verified donors are grown in batch culture with CRISPR recipient yeast at 0% U to gauge ability to depress recipient population through IDC-killing. A. <u>Growth plots of cocultures show IDC-killing in some conditions</u>. Fluorescence measurements, shown here at the end of each day, from 12 days of batch culturing of cell pairs at four concentrations of tryptophan (rows). Trp-auxotrophic recipients (S_cross_) collapse for both pairings (cutting donor—solid line—and no-*ori^T^* negative control donor—dashed line) at 1% W, but only for the cutting donor at 5% W. Lines are means of three replicates, shaded region standard deviation. Mannose was added to experiment after day 6, to break up cell clumps, shown as vertical dotted lines. A. <u>Comparing recipient population decline between cocultures, monoculture.</u> Fold-decreases in yeast growth, based on normalized fluorescence, comparing yeast in coculture to yeast monoculture. Fold decrease = - (Normalized Monoculture Citrine) / (Normalized Coculture Citrine). Points are means of three replicates, bars 95% CI. Stars represent *p-*value significance from two-sample *t-*test, with no significance for time points lacking stars (*p* > 0.05). Vertical dotted lines designate addition of mannose at day 6, to break up cell clumps. A. <u>Transient IDC counts show rise, mannose-initiated fall</u>. IDC counts (CFU) for both cell pairings (negative control donor, blue, is unable to transfer DNA, all counts = 0), over four W% (rows). After addition of mannose (vertical dotted line), most surviving cocultures drop in IDC counts. Note that 5% W yeast population is coincidentally driven to extinction just before day 6, and thus unaffected by mannose. IDC counts here are “transient” because transconjugants are terminal at 0% U, so transconjugants are unable to persist in coculture across days.

To gauge whether any effects of IDC could be reversed by interrupting cell clumps, we switched batch cultures to mannose-supplemented media after six days of growth and allowed them to grow for another six days (SF19). In both coculture pairings, subsequent yeast growth stopped declining after the media switch and persisted at steady-state levels from day six, ending trends of decline in both coculture pairings, though never recovering recipients completely to previous (higher) levels (Fig 7b, S18). From this data, we were thus able to discern the extent to which recipient populations are depressed by antagonism from coculture with bacteria, versus IDC-mediated cutting, since the positive and negative donors are equivalent for fitness and ability to adhere and form pili to recipients (Fig 7c, S20). IDC counts showed reversibility with mannose addition, with IDC dropping after day 6 (Fig 7d). Since transconjugants are terminal in 0% U, it should be noted that IDC counts are effectively transient “snapshots” of yeasts carrying IDC-Cas9 that have not yet been diluted out of batch-culture or died from starvation.

## Discussion

This work builds upon advances in synthetic microbiology that allow tunable control of microbial consortia and demonstrates the capacity to significantly alter recipient populations through contact dependent IDC. Importantly, we’ve strayed from much of the literature regarding IDC, which considers transfer rate as a percentage of recipient cells^39,43,56,67^, though some examples exist of quantifying conjugation as a rate of cell coincidence, similar to the modeling we’ve done here^62,68,69^. This is likely due to IDC’s use as a substitute for DNA transformation, but for many potential applications, the per-donor IDC rate may be equally relevant, especially if donors are utilized as a temporary probiotic. While our would-be crossfeeding mutations didn’t support steady-state obligate mutualism, auxotrophic *E. coli* were able to form commensal relationships with even WT *S. cerevisiae*, yielding some of the highest IDC values we observed. This presents an opportunity to implement conjugation systems in which only donors are engineered. This is true, too, for mannoprotein-based adhesion between bacteria and yeasts: whereas some research has shown the benefit of engineered adhesion to conjugation among bacteria^70^, the native binding strategy employed in this work avoids the need to engineer recipient cells while allowing further options for engineering donors, e.g. by making the bacterial binding mechanism inducible. Additional work is also needed to discern the short-term dynamics of cell-cell adherence, and the full extent to which mannose can reverse IDC-mediated perturbations. Previous research has demonstrated a wide repertoire of strategies *E. coli* use to derive resources from proximal producer cells, including through cell-cell nanotube connections^54,71,72^. In our case it remains unclear, however, whether E_cross_ cells can acquire nutrients through conjugative pili, via diffusion from closer recipients, or by killing clumped cells and importing lysed metabolites.

Future work should test the generalizability of these our findings, but given the ubiquity of mannoproteins among fungal cells, and the discrepancy in growth rates between prokaryotic and eukaryotic cells—which allow bacteria to adapt quickly and persist in adverse conditions—there’s ample reason to believe that similar results are achievable with other yeasts, including pathogenic fungi such as *Candida glabrata, Malassezia restricta,* and *Aspergillus fumigatus*, for which treatments are limited and in great demand^2,3,73,74^. Similarly, additional work is needed to replicate and characterize IDC rates in mixed colonies and determine the extent to which dependent bacterial donors are able to intermix despite antagonistic interactions with yeasts, especially in mixed biofilms. While we sought to spatiotemporally resolve IDC events by developing a transconjugant fluorescent reporter (see SI discussion), we were only able to determine colony-wide endpoint IDC measurements, which showed higher rates for more intermixed colonies. Further development of such a reporter and experiments using confocal microscopy would greatly benefit this endeavor.

Finally, we demonstrated that our insights relating population dynamics to IDC can be applied to functionally alter recipient yeast populations. We “rescued” a low-growing population via IDC transfer of an essential gene, by depressing donor growth and making it dependent on recipient cells, in keeping with our dynamics findings. We also “killed” recipient cells via IDC-mediated Cas9 cutting of an essential gene. In this latter case, we deliberately added a layer of complexity unnecessary to the aim of killing cells, in that we designed yeast recipients to carry the essential gene *URA3* episomally, so as to verify the cutting efficiency. Any work aiming only to depress recipient populations could just as easily target the genome, and repeatedly introduce blunt-end cut sites with no repair sequence, simultaneously disrupting the genome and draining cell resources in non-homologous end joining (NHEJ). Perhaps more interesting for future work, however, would be to target more complex functions in recipient cells, such as modifying metabolic pathways in a consortia producing a useful product, or disrupting quorum sensing function in virulent cells, for example, by targeting the farnesol pathway in *C. glabrata*. Recent work has furthermore demonstrated the capacity of conjugative DNA transfer to tune intercellular messages in a synthetic *E. coli* consortium^22^, and such a strategy could very feasibly be expanded to a wide range of both DNA programs and recipient species. This gets to the heart of IDC’s power: unlike other perturbation strategies such as Type VI Secretion Systems, which have been used for targeted killing^75,76^, the possibilities for recipient-programming via IDC are only as limited as our ability to engineer the DNA for those functions and express them in recipient populations, as well as the frequency of conjugative delivery. Our work here focuses primarily on the latter hurdle—achieving high enough IDC rates to modify recipient populations, and to control them tunably—but further work expanding the range of functions delivered to recipients could open the door to a vast array of *in situ* microbiome and synthetic consortia engineering.

## Methods

### Strain and plasmid construction

Yeast cells in this study are derived from W303 strains developed by Müller et. al. (*MAT***a** *can1-100 hmlαΔ::*BLE *leu9Δ::KANMX6 his3Δ::prACT1-ymCitrine-tADH::HIS3MX6*, with S288C version of *BUD1*)^49^. Crossfeeding yeast strains (“S_cross_”, yMM1430) have additional mutations to make them auxotrophic for tryptophan and leucine-overproducing: *LEU4^FBR^ trp2Δ::NATMX4 URA3::prACT1yCerulean-tADH1,* with leucine feedback resistance (FBR) resultant from deletion of codon 548 of *LEU4*. S_cross_ is also constitutively fluorescent for ymCitrine and yCerulean, whereas the baseline yeast used here (aka “WT yeast”, “S”, yMM1636) is only ymCitrine-fluorescent. Further mutations were introduced into these strains to make them auxotrophic for uracil and/or histidine, for IDC selection and CRISPR assay. Uracil was knocked out by amplifying a cassette of *URA3* homology arms, transforming into yMM1430, and selecting for growth on 5-Fluoroorotic acid (5FOA). *HIS3* was replaced with either *KANMX6* or *HPHMX6*, depending on the strain (see table S4 for list of strains and related experiments), by amplifying either resistance gene with overlap for *HISMX6* regions.

Bacterial strains in this study are derived from Keio Collection strains of single-gene knockouts, based on BW25113 background (*F-Δ(araD-araB)567 lacZ4787Δ::*rrnB-3 *λ-rph-1 Δ(rhaD-rhaB)568 hsdR514*)^77^. WT *E. coli* (“E”) strains are simply BW25113, or Coli Genetic Stock Center (CGSC) #7636, containing different plasmids depending on the experiment (see table S5 for list of plasmids and corresponding experiments). Crossfeeding mutations were introduced into CGSC #11110 (*ΔtrpR789:kan^R^*), which lacks the trp repressor gene, and has been shown to be tryptophan-overproducing^54^. Briefly, the kanamycin resistance gene at the *trpR* locus was “flipped” out via flippase recognition target (FRT) sequences and flippase-expressing plasmid pMM0821^78^. Leucine auxotrophy was introduced by λred recombination of PCR-amplified *ΔleuA781::kan^R^*, from CGSC #8373, using pMM0820, which expresses genes for λred. *kan^R^* was again flipped out to obtain kMM127, a double knockout of *ΔtrpR, ΔleuA*, with no antibiotic resistance. Note that we originally constructed the crossfeeding *E. coli* (“E_cross_”) from *ΔleuA::kan^R^* (CGSC #8373), but it caused severe aggregation in coculture, such that cells would precipitate out of media immediately, whereas the same mutation introduced from the *ΔtrpR::kan^R^* strain did not produce this result. Moreover, we found that *ΔleuB::kan^R^* (CGSC #11943) proved prototrophic for leucine over long time periods, despite its similar function in the leucine biosynthesis pathway (ref).

IncP-type IDC plasmids^79^ pTA-Mob 1.0 (*trans*-transferring) and pTA-Mob 2.0 (*cis*-transferring) were generously provided to us by the Karas lab^56^. pTA-Mob 2.0 contains gentamicin resistance for bacterial selection, *URA3* and *HIS3* genes for yeast selection, *CEN6/ARSH4* for yeast maintenance, and the *ori^T^* sequence required for conjugative transfer of the plasmid into recipients, whereas pTA-Mob 1.0 only carries gentamicin resistance. Constitutive bacterial reporter pMM0819 contains *pProD:mCherry*, using a synthetic reporter meant to be high-expressing and minimally susceptible to cell phase^80,81^. IDC plasmids for *trans*-transfer were constructed using the Golden Gate-based Yeast MoClo Toolkit^82^ (YTK), to modularly assemble a fluorescent yeast reporter (*pTDH3-yeBFP*), IDC selection (*HIS3*), and yeast replication machinery (*CEN6/ARSH4*). The *ori^T^* sequence was then added to the connector sequence downstream of yeBFP via Gibson assembly.

For IDC-mediated CRISPR killing assay, the S_cross_, *ura3Δ0 hismx6Δ::HPHMX6* strain yMM1786 was transformed with a plasmid containing *pTDH3-yeBFP URA3 CEN/ARS*. sgRNAs were designed to cut within the connector region of this plasmid (ConR1 from YTK), downstream of yeBFP, such that any YTK-assembled plasmid containing the ConR1 sequence could be a target in future experiments. CRISPR plasmids were assembled using Ellis lab plasmids^83,84^. Briefly, oligos for five sgRNA sequences targeting the ConR1 region were designed using Benchling^85^, PNK-phosphorylated and annealed. Annealed oligos were then assembled into sgRNA entry vector pMM1340 via Golden Gate assembly and transformed into bacteria, selecting with carbenicillin. Purified and sequence-verified sgRNA plasmids were then digested with EcoRV to isolate the sgRNA sequences with homology arms matching the insertion site of the Cas9 plasmid. Plasmid pMM1341, which contains Cas9, GFP, and *HIS3*, was digested with BsmBI to remove GFP and leave homology arms for sgRNA at each end of the resultant linear DNA. The two pieces were combined via yeast recombinant cloning. Finally, *ori^T^* was inserted by ligating a modified version of the *ori^T^* sequence with the assembled Cas9-sgRNA after digesting with AatII and SacII.

### Batch culture experiments and IDC counting

Yeast and bacterial cultures used in each batch culture experiment were grown overnight in selective YPD or LB media, at 30°C or 37°C, respectively. After ≥ 16 hours’ growth, bacterial strains were measured for OD600, yeast strains were measured for OD660, and each culture was washed at least 2 times with SC or M9 sans glucose or amino acids. Cells were then combined such that each reaction started with 1E7 cells, based on OD measurements. Growth media was composed of 200 μL of a 75:25 mixture of SC:M9 minimal media (see SI discussion) with 2% glucose, appropriate amino acids, and antibiotics to maintain each bacterial plasmid. Amino acid percentages in the text are based on the following molarities, considered 100%: L = 762 μM, W = 245 μM, U = 178 μM, H = 95.4 μM. For clumping experiment (Fig 3), half of the media was supplemented with 4% mannose. Upon spiking cells into 96-well CellVis back-walled optical glass-bottom plates (cat #P96-1-N), plates were sealed with gas permeable membranes (Fisher Scientific cat #50-550-304) seals to allow air flow for aerobic conditions.

Plates were grown in a customized Tecan Fluent automated plate handling robot, on a Bioshakes heater-shaker, kept at 30°C and rotating at 1000 rpm with a 2 mm orbital. In 15 minute intervals, the Fluent was programmed to transfer each 96-well plate to a connected Tecan Spark fluorimeter, in which each well was measured for OD600, mCherry (Ex=575nm, Em=620nm, 20nm bandwidth, gain=60), ymCitrine (Ex=500nm, Em=545nm, 20nm bandwidth, gain=60), and, for CRISPR experiment (Fig 7), yeBFP (Ex=381nm, Em=445nm, 20nm bandwidth, gain=60). After each plate was measured, it was returned to the Bioshakes, where it grew for another 15 minutes until the next read. Each plate was grown in this way for roughly 18-24 hr., at which time plates were briefly spun (1 min at 1000xG) to remove droplets from plate seal. Each plate was then diluted 1:10 in fresh media (180 μL media + 20 μL previous day’s culture) for that day’s growth, with another 20 μL diluted into a plate of PBS+0.1% Tween for flow cytometry (see below). Tecan data was consolidated in Excel format and imported into MATLAB via a custom script, which parses the Tecan Excel export format based on number of plates and channels measured. All further analyses were performed in MATLAB, including normalization, in which all fluorescence measurements were divided by the max reading of that channel; these normalized reads were used for D:R ratios in Fig 2 and SF3.

An additional 100 μL of each day’s culture was added, undiluted, to a 24-well plate containing IDC-selective SC with 2% agar: for *cis*-transfer experiments (Figs 2, 5, and 6), SC-UH was used, whereas SC-H was used for *trans*-transfer experiments (Figs 3, 5, and 7). IDC plates were then placed in a culture shaker at 30°C for ∼40 minutes, without lids, to dry. Once dried, IDC plates were incubated for ∼3 days to grow countable transconjugant colonies. Individual transconjugant colonies were counted for CFU, unless wells were saturated, for which estimates were generated based on density relative to countable wells, up to 500. Samples with no transconjugants were set to 0.1 in log-scale plots, to account for detection limit. For rescue assay, due to higher counts, cultures after day 2 were serial-diluted up to 1:10,000, in increments of 10x dilutions, and frogged onto SC-UH 2% agar in a 245 mm BioAssay Dish (Corning cat # 431111). Countable microcolonies from frogging dilutions were averaged, based on dilution value; thus, saturated microcolonies were ignored.

### Colony experiments

Each strain was grown, measured, washed, and diluted as in batch culture experiments. Because we wanted 2 μL mixed culture droplets to seed each colony, we had to lower the input cell counts to 1E6 of each cell type. Strains were combined accordingly, then aliquoted into striptubes, from which we were able to multichannel-pipette ≥ 18 identical 2 μL mixed colonies onto 2% agar minimal media plates. Each 60 mm plate (Eppendorf cat #0030701011) contained 75:25 SC:M9 with appropriate bacterial antibiotics for plasmid maintenance, 2% agar, and one concentration of amino acids, such that each plate represented a single experimental condition; molten media was aliquoted to plates in equal (15 mL) portions. Once mixed colonies were added to plates, they were allowed to grow at 30°C for 6 days. Three representative colonies (by eye) were designated after the first day’s growth to be repeatedly imaged over the entire time course, while another three were designated to be imaged that day only, after which they would be scraped, washed, and measured by flow cytometry and IDC-plating. All colonies were numbered, upon being selected, to correlate measurements.

Plates were imaged for fluorescence using a Zeiss AxioZoom V16 dissecting microscope, at UW-Madison’s Newcomb Imaging Center. Each *cis-*donor mixed colony was imaged for mCherry (Zeiss Set 43 BP 545/25, FT 570, BP 605/70, 200 ms exposure) and ymCitrine (Zeiss Set 46 HE, EX BP 500/20, BS FT 515, EM BP 535/30, 600 ms exposure), while *trans-*donor mixed colonies were additionally measured for yeBFP (Zeiss set 49: G365, FT395, BP445/50, 500 ms exposure). All images were taken at 8x zoom. See SI discussion for more information on image processing and analysis.

After imaging, colonies were manually scraped off plates via micropipette tips and diluted into 1.5 mL tubes containing 1 mL water. Each diluted colony was vortexed for ∼30 s to break up colonies and dilute residual agar, then spun at 3000xG for 5 min. 800 μL of water was removed from each tube, cells were resuspended in the remaining ∼200 μL, 100 μL of which was plated for IDC-selection (see Batch Culture methods) and another 20 μL was aliquoted into 180 μL PBS+0.1% Tween for flow cytometry (see below). After six days of growth, the colonies (1-3) designated for continual microscopy imaging were processed and measured in this way.

### Flow cytometry

After diluting cells from culture or colonies (see above) 1:10 into PBS+0.1% Tween (total volume=200 μL) in 96-well round-bottom plates (Fisher Scientific cat #07-200-760), samples were measured for cellular composition using a ThermoFisher Attune NxT V6 Flow Cytometer at UW-Madison’s Carbone Cancer Center, which includes a 96-well compatible autosampler. Because the sizes of bacteria and yeast are so different, each coculture was measured twice, with different forward and side scatter voltages for each cell type (monoculture controls were generally measured using only that species’ voltage settings, though at least two of the other species were included for each to get baseline counts). Each well was measured for ymCitrine (488 nm laser, 530/30 503LP filters, off target fluorescent) and mCherry (561 nm laser, 620/15 600LP filters), in addition to scatter, using a draw volume of 20 μL, at a flow rate of 200 μL/min.

FCS files exported from the Attune were processed via custom MATLAB tools modified for dual-voltage experiments. Gates were drawn per voltage setting to capture all cells of that species based on fluorescence and forward scatter. FCS files were imported, correlated with sample information, and queried for inclusion in each gate. Summary tables for each cell type were consolidated to combine all readings per experiment, upon which noise floors were calculated based on negative controls per voltage setting. Gate-defined cell counts for each species were subtracted by these baselines and converted to total cells per 100 μL, to compare to IDC counts (see IDC prep in Batch Culture methods). Cell counts were further normalized by dividing by the max count for that experiment and cell type; normalized counts were used to generate D:R ratios (Figs 2, 5, 6).

### Microscopy of culture aggregates

Batch culture samples were diluted to various degrees (depending on day and sample density) in media lacking glucose and amino acids, but with mannose for samples grown with it, in a CellVis 96-well back-walled optical glass-bottom plates (cat #P96-1-N). Plates were loaded onto the stage of an inverted fluorescence microscope (Nikon TiE), enclosed by an opaque incubation chamber. A custom Nikon JOBS script was written to image each well of a plate in three random locations distal to the well edges, with a two second wait time before each photo to allow cells to settle after moving the stage. All wells were imaged at 10x objective for mCherry (Chroma 96365, Ex = 560/40x, Em = 630/75m, 200 ms exposure), ymCitrine (Chroma 96363, Ex = 500/20x, Em = 535/30m, 600 ms exposure), and yeBFP (Chroma NC296093, Ex = 350/50x, Em = 460/50m, 500 ms exposure). See SI discussion for image processing and analysis.

### Statistics

Two-sample *t-*tests calculated using MATLAB’s “ttest2” function, with default two tails (Figs 3c, 5c, 7c). 1-way ANOVA of ICQ calculated between coculture pairings in Fig 5c via MATLAB’s “anovan” function, using type II sum of squares, alpha = 0.5 (significance level ≤ 0.5). See Table S6 for F-values and degrees of freedom for 1-way ANOVAs for Fig 5c. Significance stars indicate *p*-values less than 0.05 (*), 0.01 (**), or 0.005 (***), or else are considered not significant. Number of replicates cited in each figure caption.

### Data analysis and figures

Unless otherwise specified, all data processing was performed using custom MATLAB scripts, which can be accessed at the GitHub repository for this paper: https://github.com/mccleanlab/Stindt_2023. Most data plots were generated with the gramm MATLAB toolbox^86^ and flow diagrams were created with BioRender.com.

## Acknowledgments

We would like to thank Taylor Scott and Kieran Sweeney for invaluable feedback on this work and help with coding, and the McClean Lab for many helpful discussions. Conjugative plasmids pTA-Mob were generously provided to us by the Karas Lab, Western University, Ontario; many thanks to Bogumil, Ryan Cochrane, and Maximillian Soltysiak for their generosity in mailing and emailing. This work was supported by NIGMS (MIRA R35GM128873), NIAID (R01AI154940), and the Wisconsin Alumni Research Foundation (WARF 135 AAI9593). Flow cytometry was enabled by the University of Wisconsin Carbone Cancer Center Support Grant (P30CA014520). Megan Nicole McClean, PhD holds a Career Award at the Scientific Interface from the Burroughs Wellcome Fund. Kevin Stindt was supported in part by the Molecular Biophysics Training Grant (T32GM130550).

## Author Contributions

KS and MNM planned experiments. KS performed the experiments. KS and MNM analyzed the data and wrote the paper.

## Data Availability

The data that support the findings of this study are openly available in Dryad at https://doi.org/10.5061/dryad.5mkkwh7c7.

## REFERENCES

1. Huffnagle, G.B., and Noverr, M.C. (2013). The emerging world of the fungal microbiome. Trends Microbiol 21, 334–341. 10.1016/j.tim.2013.04.002.

2. Pappas, P.G., Lionakis, M.S., Arendrup, M.C., Ostrosky-Zeichner, L., and Kullberg, B.J. (2018). Invasive candidiasis. Nat Rev Dis Primers 4, 18026. 10.1038/nrdp.2018.26.

3. Limon, J.J., Tang, J., Li, D., Wolf, A.J., Michelsen, K.S., Funari, V., Gargus, M., Nguyen, C., Sharma, P., Maymi, V.I., et al. (2019). Malassezia Is Associated with Crohn’s Disease and Exacerbates Colitis in Mouse Models. Cell Host Microbe 25, 377–388.e6. 10.1016/j.chom.2019.01.007.

4. Aykut, B., Pushalkar, S., Chen, R., Li, Q., Abengozar, R., Kim, J.I., Shadaloey, S.A., Wu, D., Preiss, P., Verma, N., et al. (2019). The fungal mycobiome promotes pancreatic oncogenesis via activation of MBL. Nature 574, 264–267. 10.1038/s41586-019-1608-2.

5. Casadevall, A. (2018). Fungal Diseases in the 21st Century: The Near and Far Horizons. Pathog Immun 3, 183–196. 10.20411/pai.v3i2.249.

6. Kalan, L., Loesche, M., Hodkinson, B.P., Heilmann, K., Ruthel, G., Gardner, S.E., and Grice, A. A. (2016). Redefining the Chronic-Wound Microbiome: Fungal Communities Are Prevalent, Dynamic, and Associated with Delayed Healing. mBio 7, e01058–16. 10.1128/mBio.01058-16.

7. Doehlemann, G., Ökmen, B., Zhu, W., and Sharon, A. (2017). Plant Pathogenic Fungi. Microbiology Spectrum 5, 10.1128/microbiolspec.funk-0023–2016. 10.1128/microbiolspec.funk-0023-2016.

8. Seyedmousavi, S., Bosco, S. de M.G., de Hoog, S., Ebel, F., Elad, D., Gomes, R.R., Jacobsen, I.D., Jensen, H.E., Martel, A., Mignon, B., et al. (2018). Fungal infections in animals: a patchwork of different situations. Med Mycol 56, S165–S187. 10.1093/mmy/myx104.

9. Cornelison, C.T., Keel, M.K., Gabriel, K.T., Barlament, C.K., Tucker, T.A., Pierce, G.E., and Crow, S.A. (2014). A preliminary report on the contact-independent antagonism of Pseudogymnoascus destructans by Rhodococcus rhodochrous strain DAP96253. BMC Microbiol 14, 246. 10.1186/s12866-014-0246-y.

10. Wu, N.C., Cramp, R.L., Ohmer, M.E.B., and Franklin, C.E. (2019). Epidermal epidemic: unravelling the pathogenesis of chytridiomycosis. Journal of Experimental Biology 222, jeb191817. 10.1242/jeb.191817.

11. Ponomarova, O., Gabrielli, N., Sévin, D.C., Mülleder, M., Zirngibl, K., Bulyha, K., Andrejev, S., Kafkia, E., Typas, A., Sauer, U., et al. (2017). Yeast Creates a Niche for Symbiotic Lactic Acid Bacteria through Nitrogen Overflow. Cell Systems 5, 345–357.e6. 10.1016/j.cels.2017.09.002.

12. Villarreal-Soto, S.A., Beaufort, S., Bouajila, J., Souchard, J.-P., and Taillandier, P. (2018). Understanding Kombucha Tea Fermentation: A Review. Journal of Food Science 83, 580–588. 10.1111/1750-3841.14068.

13. Mittermeier, F., Bäumler, M., Arulrajah, P., García Lima, J. de J., Hauke, S., Stock, A., and Weuster-Botz, D. (2023). Artificial microbial consortia for bioproduction processes. Engineering in Life Sciences 23, e2100152. 10.1002/elsc.202100152.

14. Parapouli, M., Vasileiadis, A., Afendra, A.-S., and Hatziloukas, E. (2020). Saccharomyces cerevisiae and its industrial applications. AIMS Microbiol 6, 1–31. 10.3934/microbiol.2020001.

15. Amobonye, A., Aruwa, C.E., Aransiola, S., Omame, J., Alabi, T.D., and Lalung, J. (2023). The potential of fungi in the bioremediation of pharmaceutically active compounds: a comprehensive review. Front Microbiol 14, 1207792. 10.3389/fmicb.2023.1207792.

16. Sheth, R.U., Cabral, V., Chen, S.P., and Wang, H.H. (2016). Manipulating Bacterial Communities by in situ Microbiome Engineering. Trends in Genetics 32, 189–200. 10.1016/j.tig.2016.01.005.

17. Villa, T.G., Feijoo-Siota, L., Sánchez-Pérez, A., Rama, JL.R., and Sieiro, C. (2019). Horizontal Gene Transfer in Bacteria, an Overview of the Mechanisms Involved. In Horizontal Gene Transfer: Breaking Borders Between Living Kingdoms, T. G. Villa and M. Viñas, eds. (Springer International Publishing), pp. 3–76. 10.1007/978-3-030-21862-1_1.

18. Our Solutions | Atterx Biotherapeutics https://www.atterx.com/gn-4474.

19. López-Igual, R., Bernal-Bayard, J., Rodríguez-Patón, A., Ghigo, J.-M., and Mazel, D. (2019). Engineered toxin–intein antimicrobials can selectively target and kill antibiotic-resistant bacteria in mixed populations. Nat Biotechnol 37, 755–760. 10.1038/s41587-019-0105-3.

20. Razzaq, A., Saleem, F., Kanwal, M., Mustafa, G., Yousaf, S., Imran Arshad, H.M., Hameed, M.K., Khan, M.S., and Joyia, F.A. (2019). Modern Trends in Plant Genome Editing: An Inclusive Review of the CRISPR/Cas9 Toolbox. International Journal of Molecular Sciences 20, 4045. 10.3390/ijms20164045.

21. Brophy, J.A.N., Triassi, A.J., Adams, B.L., Renberg, R.L., Stratis-Cullum, D.N., Grossman, A.D., and Voigt, C.A. (2018). Engineered integrative and conjugative elements for efficient and inducible DNA transfer to undomesticated bacteria. Nature Microbiology 3, 1043–1053. 10.1038/s41564-018-0216-5.

22. Marken, J.P., and Murray, R.M. (2023). Addressable and adaptable intercellular communication via DNA messaging. Nat Commun 14, 2358. 10.1038/s41467-023-37788-z.

23. Ronda, C., Chen, S.P., Cabral, V., Yaung, S.J., and Wang, H.H. (2019). Metagenomic engineering of the mammalian gut microbiome in situ. Nat Methods 16, 167–170. 10.1038/s41592-018-0301-y.

24. Neil, K., Allard, N., Roy, P., Grenier, F., Menendez, A., Burrus, V., and Rodrigue, S. (2021). High-efficiency delivery of CRISPR-Cas9 by engineered probiotics enables precise microbiome editing. Molecular Systems Biology 17, e10335. 10.15252/msb.202110335.

25. Bourras, S., Rouxel, T., and Meyer, M. (2015). Agrobacterium tumefaciens Gene Transfer: How a Plant Pathogen Hacks the Nuclei of Plant and Nonplant Organisms. Phytopathology 105, 1288–1301. 10.1094/PHYTO-12-14-0380-RVW.

26. Karas, B.J., Diner, R.E., Lefebvre, S.C., McQuaid, J., Phillips, A.P.R., Noddings, C.M., Brunson, J.K., Valas, R.E., Deerinck, T.J., Jablanovic, J., et al. (2015). Designer diatom episomes delivered by bacterial conjugation. Nature Communications 6, 6925. 10.1038/ncomms7925.

27. Diner, R.E., Bielinski, V.A., Dupont, C.L., Allen, A.E., and Weyman, P.D. (2016). Refinement of the Diatom Episome Maintenance Sequence and Improvement of Conjugation-Based DNA Delivery Methods. Front. Bioeng. Biotechnol. 4. 10.3389/fbioe.2016.00065.

28. Slattery, S.S., Diamond, A., Wang, H., Therrien, J.A., Lant, J.T., Jazey, T., Lee, K., Klassen, Z., Desgagné-Penix, I., Karas, B.J., et al. (2018). An Expanded Plasmid-Based Genetic Toolbox Enables Cas9 Genome Editing and Stable Maintenance of Synthetic Pathways in Phaeodactylum tricornutum. ACS Synth. Biol. 7, 328–338. 10.1021/acssynbio.7b00191.

29. Sharma, A.K., Nymark, M., Sparstad, T., Bones, A.M., and Winge, P. (2018). Transgene-free genome editing in marine algae by bacterial conjugation – comparison with biolistic CRISPR/Cas9 transformation. Scientific Reports 8, 1–11. 10.1038/s41598-018-32342-0.

30. Waters, V.L. (2001). Conjugation between bacterial and mammalian cells. Nat Genet 29, 375–376. 10.1038/ng779.

31. Heinemann, J.A., and Jr, G.F.S. (1989). Bacterial conjugative plasmids mobilize DNA transfer between bacteria and yeast. Nature 340, 205–209. 10.1038/340205a0.

32. Nishikawa, M., and Yoshida, K. (1998). Trans-kingdom conjugation offers a powerful gene targeting: tool in yeast. Genetic Analysis: Biomolecular Engineering 14, 65–73. 10.1016/S1050-3862(97)10003-1.

33. Nishikawa, M., Suzuki, K., and Yoshida, K. (1992). DNA integration into recipient yeast chromosomes by trans-kingdom conjugation between Escherichia coli and Saccharomyces cerevisiae. Curr Genet 21, 101–108. 10.1007/BF00318467.

34. Inomata, K., Nishikawa, M., and Yoshida, K. (1994). The yeast Saccharomyces kluyveri as a recipient eukaryote in transkingdom conjugation: behavior of transmitted plasmids in transconjugants. J Bacteriol 176, 4770–4773.

35. Suzuki, K., Moriguchi, K., and Yamamoto, S. (2015). Horizontal DNA transfer from bacteria to eukaryotes and a lesson from experimental transfers. Research in Microbiology 166, 753– 763. 10.1016/j.resmic.2015.08.001.

36. Woese, C.R., Kandler, O., and Wheelis, M.L. (1990). Towards a natural system of organisms: proposal for the domains Archaea, Bacteria, and Eucarya. Proc Natl Acad Sci U S A 87, 4576–4579. 10.1073/pnas.87.12.4576.

37. Waksman, G. (2019). From conjugation to T4S systems in Gram-negative bacteria: a mechanistic biology perspective. EMBO reports 20, e47012. 10.15252/embr.201847012.

38. Christie, P.J. (2016). The Mosaic Type IV Secretion Systems. EcoSal Plus 7. 10.1128/ecosalplus.ESP-0020-2015.

39. Cochrane, R.R., Shrestha, A., Severo de Almeida, M.M., Agyare-Tabbi, M., Brumwell, S.L., Hamadache, S., Meaney, J.S., Nucifora, D.P., Say, H.H., Sharma, J., et al. (2022). Superior Conjugative Plasmids Delivered by Bacteria to Diverse Fungi. BioDesign Research 2022. 10.34133/2022/9802168.

40. Getino, M., and Cruz, F. de la (2018). Natural and Artificial Strategies To Control the Conjugative Transmission of Plasmids. Microbiology Spectrum 6. 10.1128/microbiolspec.MTBP-0015-2016.

41. Filutowicz, M., Burgess, R., Gamelli, R.L., Heinemann, J.A., Kurenbach, B., Rakowski, S.A., and Shankar, R. (2008). Bacterial conjugation-based antimicrobial agents. Plasmid 60, 38–44. 10.1016/j.plasmid.2008.03.004.

42. Lawrence, J.G., and Retchless, A.C. (2009). The Interplay of Homologous Recombination and Horizontal Gene Transfer in Bacterial Speciation. In Horizontal Gene Transfer: Genomes in Flux Methods in Molecular Biology., M. B. Gogarten, J. P. Gogarten, and L. C. Olendzenski, eds. (Humana Press), pp. 29–53. 10.1007/978-1-60327-853-9_3.

43. Moriguchi, K., Yamamoto, S., Ohmine, Y., and Suzuki, K. (2016). A Fast and Practical Yeast Transformation Method Mediated by Escherichia coli Based on a Trans-Kingdom Conjugal Transfer System: Just Mix Two Cultures and Wait One Hour. PLOS ONE 11, e0148989. 10.1371/journal.pone.0148989.

44. Gietz, R.D., and Schiestl, R.H. (2007). High-efficiency yeast transformation using the LiAc/SS carrier DNA/PEG method. Nat Protoc 2, 31–34. 10.1038/nprot.2007.13.

45. Hamilton, T.A., Pellegrino, G.M., Therrien, J.A., Ham, D.T., Bartlett, P.C., Karas, B.J., Gloor, G.B., and Edgell, D.R. (2019). Efficient inter-species conjugative transfer of a CRISPR nuclease for targeted bacterial killing. Nature Communications 10, 1–9. 10.1038/s41467-019-12448-3.

46. Tolker-Nielsen, T., and Molin, S. (2000). Spatial Organization of Microbial Biofilm Communities. Microb Ecol 40, 75–84. 10.1007/s002480000057.

47. Freese, P.D., Korolev, K.S., Jiménez, J.I., and Chen, I.A. (2014). Genetic Drift Suppresses Bacterial Conjugation in Spatially Structured Populations. Biophys J 106, 944–954. 10.1016/j.bpj.2014.01.012.

48. Blanchard, A.E., and Lu, T. (2015). Bacterial social interactions drive the emergence of differential spatial colony structures. BMC Systems Biology 9, 59. 10.1186/s12918-015-0188-5.

49. Muller, M.J.I., Neugeboren, B.I., Nelson, D.R., and Murray, A.W. (2014). Genetic drift opposes mutualism during spatial population expansion. Proceedings of the National Academy of Sciences 111, 1037–1042. 10.1073/pnas.1313285111.

50. Scarinci, G., and Sourjik, V. (2023). Impact of direct physical association and motility on fitness of a synthetic interkingdom microbial community. ISME J 17, 371–381. 10.1038/s41396-022-01352-2.

51. Miozzari, G., Niederberger, P., and Hütter, R. (1978). Tryptophan biosynthesis in Saccharomyces cerevisiae: control of the flux through the pathway. J Bacteriol 134, 48–59.

52. Cavalieri, D., and Polsinelli, M. Trīuoroleucine resistance and regulation of a-isopropyl malate synthase in Saccharomyces cerevisiae. 9.

53. Kohlhaw, G., Leary, T.R., and Umbarger, H.E. (1969). Alpha-isopropylmalate synthase from Salmonella typhimurium. Purification and properties. J Biol Chem 244, 2218–2225.

54. Pande, S., Shitut, S., Freund, L., Westermann, M., Bertels, F., Colesie, C., Bischofs, I.B., and Kost, C. (2015). Metabolic cross-feeding via intercellular nanotubes among bacteria. Nature Communications 6, 6238. 10.1038/ncomms7238.

55. Bianchi, F., van’t Klooster, J.S., Ruiz, S.J., and Poolman, B. (2019). Regulation of Amino Acid Transport in Saccharomyces cerevisiae. Microbiology and Molecular Biology Reviews 83, 10.1128/mmbr.00024-19. 10.1128/mmbr.00024-19.

56. Soltysiak, M.P.M., Meaney, R.S., Hamadache, S., Janakirama, P., Edgell, D.R., and Karas, B.J. (2019). Trans-Kingdom Conjugation within Solid Media from Escherichia coli to Saccharomyces cerevisiae. International Journal of Molecular Sciences 20, 5212. 10.3390/ijms20205212.

57. Gow, N.A.R., Latge, J.-P., and Munro, C.A. (2017). The Fungal Cell Wall: Structure, Biosynthesis, and Function. Microbiology Spectrum 5, 10.1128/microbiolspec.funk-0035–2016. 10.1128/microbiolspec.funk-0035-2016.

58. Jann, K., Schmidt, G., Blumenstock, E., and Vosbeck, K. (1981). Escherichia coli adhesion to Saccharomyces cerevisiae and mammalian cells: role of piliation and surface hydrophobicity. Infect Immun 32, 484–489.

59. Ofek, I., and Beachey, E.H. (1978). Mannose Binding and Epithelial Cell Adherence of Escherichia coli. Infection and Immunity 22, 247–254.

60. Hoek, T.A., Axelrod, K., Biancalani, T., Yurtsev, E.A., Liu, J., and Gore, J. (2016). Resource Availability Modulates the Cooperative and Competitive Nature of a Microbial Cross-Feeding Mutualism. PLOS Biology 14, e1002540. 10.1371/journal.pbio.1002540.

61. Wu, F., Lopatkin, A.J., Needs, D.A., Lee, C.T., Mukherjee, S., and You, L. (2019). A unifying framework for interpreting and predicting mutualistic systems. Nat Commun 10. 10.1038/s41467-018-08188-5.

62. Levin, B.R., Stewart, F.M., and Rice, V.A. (1979). The kinetics of conjugative plasmid transmission: Fit of a simple mass action model. Plasmid 2, 247–260. 10.1016/0147-619X(79)90043-X.

63. Volkova, V.V., Lu, Z., Lanzas, C., Scott, H.M., and Gröhn, Y.T. (2013). Modelling dynamics of plasmid-gene mediated antimicrobial resistance in enteric bacteria using stochastic differential equations. Scientific Reports 3, 2463. 10.1038/srep02463.

64. Hallatschek, O., Hersen, P., Ramanathan, S., and Nelson, D.R. (2007). Genetic drift at expanding frontiers promotes gene segregation. Proc Natl Acad Sci U S A 104, 19926– 19930. 10.1073/pnas.0710150104.

65. Fusco, D., Gralka, M., Kayser, J., Anderson, A., and Hallatschek, O. (2016). Excess of mutational jackpot events in expanding populations revealed by spatial Luria–Delbrück experiments. Nat Commun 7, 12760. 10.1038/ncomms12760.

66. Li, Q., Lau, A., Morris, T.J., Guo, L., Fordyce, C.B., and Stanley, E.F. (2004). A Syntaxin 1, Gαo, and N-Type Calcium Channel Complex at a Presynaptic Nerve Terminal: Analysis by Quantitative Immunocolocalization. J Neurosci 24, 4070–4081. 10.1523/JNEUROSCI.0346-04.2004.

67. Moriguchi, K., Edahiro, N., Yamamoto, S., Tanaka, K., Kurata, N., and Suzuki, K. (2013). Transkingdom Genetic Transfer from Escherichia coli to Saccharomyces cerevisiae as a Simple Gene Introduction Tool. Appl Environ Microbiol 79, 4393–4400. 10.1128/AEM.00770-13.

68. Simonsen, L., Gordon, D.M., Stewart, F.M., and Levin, B.R. (1990). Estimating the rate of plasmid transfer: an end-point method. Microbiology, 136, 2319–2325. 10.1099/00221287-136-11-2319.

69. Zhong, X., Droesch, J., Fox, R., Top, E.M., and Krone, S.M. (2012). On the meaning and estimation of plasmid transfer rates for surface-associated and well-mixed bacterial populations. Journal of Theoretical Biology 294, 144–152. 10.1016/j.jtbi.2011.10.034.

70. Robledo, M., Álvarez, B., Cuevas, A., González, S., Ruano-Gallego, D., Fernández, L.Á., and de la Cruz, F. (2022). Targeted bacterial conjugation mediated by synthetic cell-to-cell adhesions. Nucleic Acids Research 50, 12938–12950. 10.1093/nar/gkac1164.

71. Germerodt, S., Bohl, K., Lück, A., Pande, S., Schröter, A., Kaleta, C., Schuster, S., and Kost, C. (2016). Pervasive Selection for Cooperative Cross-Feeding in Bacterial Communities. PLOS Computational Biology 12, e1004986. 10.1371/journal.pcbi.1004986.

72. Preussger, D., Giri, S., Muhsal, L.K., Oña, L., and Kost, C. (2020). Reciprocal Fitness Feedbacks Promote the Evolution of Mutualistic Cooperation. Current Biology 30, 3580–3590.e7. 10.1016/j.cub.2020.06.100.

73. Rodrigues, C.F., Rodrigues, M.E., Silva, S., and Henriques, M. (2017). Candida glabrata Biofilms: How Far Have We Come? J Fungi (Basel) 3, 11. 10.3390/jof3010011.

74. Dagenais, T.R.T., and Keller, N.P. (2009). Pathogenesis of Aspergillus fumigatus in Invasive Aspergillosis. Clinical Microbiology Reviews 22, 447–465. 10.1128/cmr.00055-08.

75. Singh, R.P., and Kumari, K. (2023). Bacterial type VI secretion system (T6SS): an evolved molecular weapon with diverse functionality. Biotechnol Lett 45, 309–331. 10.1007/s10529-023-03354-2.

76. Kreitz, J., Friedrich, M.J., Guru, A., Lash, B., Saito, M., Macrae, R.K., and Zhang, F. (2023). Programmable protein delivery with a bacterial contractile injection system. Nature 616, 357–364. 10.1038/s41586-023-05870-7.

77. Baba, T., Ara, T., Hasegawa, M., Takai, Y., Okumura, Y., Baba, M., Datsenko, K.A., Tomita, M., Wanner, B.L., and Mori, H. (2006). Construction of Escherichia coli K-12 in-frame, single-gene knockout mutants: the Keio collection. Mol Syst Biol 2, 2006.0008. 10.1038/msb4100050.

78. Datsenko, K.A., and Wanner, B.L. (2000). One-step inactivation of chromosomal genes in Escherichia coli K-12 using PCR products. Proc Natl Acad Sci U S A 97, 6640–6645.

79. Sen, D., Van der Auwera, G.A., Rogers, L.M., Thomas, C.M., Brown, C.J., and Top, E.M. (2011). Broad-Host-Range Plasmids from Agricultural Soils Have IncP-1 Backbones with Diverse Accessory Genes▿. Appl Environ Microbiol 77, 7975–7983. 10.1128/AEM.05439-11.

80. Davis, J.H., Rubin, A.J., and Sauer, R.T. (2011). Design, construction and characterization of a set of insulated bacterial promoters. Nucleic Acids Res 39, 1131–1141. 10.1093/nar/gkq810.

81. pCDF-mcherry1 was a gift from Michael Lynch (Addgene plasmid # 87144 ; http://n2t.net/addgene:87144 ; RRID:Addgene_87144).

82. Lee, M.E., DeLoache, W.C., Cervantes, B., and Dueber, J.E. (2015). A Highly Characterized Yeast Toolkit for Modular, Multipart Assembly. ACS Synth. Biol. 4, 975–986. 10.1021/sb500366v.

83. Ryan, O.W., Skerker, J.M., Maurer, M.J., Li, X., Tsai, J.C., Poddar, S., Lee, M.E., DeLoache, W., Dueber, J.E., Arkin, A.P., et al. (2014). Selection of chromosomal DNA libraries using a multiplex CRISPR system. eLife 3, e03703. 10.7554/eLife.03703.

84. Horwitz, A.A., Walter, J.M., Schubert, M.G., Kung, S.H., Hawkins, K., Platt, D.M., Hernday, A.D., Mahatdejkul-Meadows, T., Szeto, W., Chandran, S.S., et al. (2015). Efficient Multiplexed Integration of Synergistic Alleles and Metabolic Pathways in Yeasts via CRISPR-Cas. Cell Syst 1, 88–96. 10.1016/j.cels.2015.02.001.

85. Design guide RNAs (gRNAs) (2022). Benchling. https://help.benchling.com/hc/en-us/articles/9684282104333-Design-guide-RNAs-gRNAs-.

86. Morel, P. (2018). Gramm: grammar of graphics plotting in Matlab. Journal of Open Source Software 3, 568. 10.21105/joss.00568.

