## Supplemental Information for "Tuning Interdomain Conjugation Toward *in situ* Population Modification in Yeast"

#### Coculture condition screening

Because we hoped to tune nutrients (especially amino acids) in batch culture media, we sought to use minimal media to grow cocultures containing *E. coli* and *S. cerevisiae*. We performed initial experiments in equal parts M9 bacterial minimal media and synthetic complete (SC) yeast media, with glucose levels at the higher SC-media concentration of 2%. These experiments resulted in cocultures heavily dominated by bacteria, with no stable obligate mutualism at 0% leucine and tryptophan.

Thus, we screened four experimental conditions for steady-state maintenance of each strain over several days, via Tecan Spark fluorescence plate reader, especially looking for maintenance of obligate mutualism at 0% LW (Fig S1). 1) A range of L and W concentrations at 50:50 SC:M9 were used to get a full sense of whether any benefit between crossfeeding strains could be found, even at LW > 0%. We saw minimal benefits for crossfeeder *E. coli* ($E_{cross}$) grown in cocultures, but they were inconsistent over time. 2) We increased initial cell densities, with each strain combined in equal proportions, from 2-fold overall density (2E7 each cell type per well) to 5-fold density (5E7 cells of each strain). We grew all samples in this and subsequent conditions at 0% LW. At each increased density, crossfeeder bacteria showed an ability to survive off of yeasts, especially crossfeeder yeasts ($S_{cross}$), but crossfeeder yeasts weren't able to survive off bacteria. 3) We varied the input ratio of cells from 10:1 yeast-to-bacteria to 1000:1 yeast-to-bacteria, all up from the previous 1:1. These results showed varying levels of $E_{cross}$ survival in coculture, but as before, no corresponding $S_{cross}$ survival. 4) We changed the proportion of minimal media from equal parts SC-to-M9, to a range of SC:M9 ratios: 60:40, 75:25, 90:10, and 100:0. All ratios showed an improved ability for $S_{cross}$ to survive in culture at 0% LW, though all bacteria were unable to survive in 100% SC; this latter observation proved very useful in selecting for yeasts, as IDC-selection plates didn't require any bacterial antibiotics at 100% SC agar. Of the media ratios tested, the 75:25 SC:M9 results showed the best ratio of crossfeeder paired growth vs. pairings with a WT strain, and so that media type was used in all subsequent experiments. Importantly, though, this ability of $S_{cross}$ to survive at 0% W was never seen again, though the steady state concentrations of yeast cells in other conditions were much more favorable to yeast and allowed better population tuning than 50:50 SC:M9.

#### Image Analyses

##### Identifying clumps in mannose assay

Microscopy images from batch cocultures with or without supplemented mannose were taken after each day's growth. Initially, we diluted samples 1:50 into corresponding media (mannose +/- ) without glucose for imaging, though as day-end cell densities diverged between samples, additional dilutions were imaged, from 1:1 to 1:500, depending on the sample and day. A custom Nikon "JOBS" script imaged each well of dilution plates at three random locations, using filter cubes for mCherry, ymCitrine, and yeBFP, at 10x objective. All .nd2 files were converted to lossless .tif files using a custom ImageJ macro. Sample information for TIFF files were imported into a MATLAB table, including sample information, dilution, and whether mannose was included.

We binarized each .tif file via MATLAB's adaptive thresholding, using thresholding values of 0.4 for both mCherry (bacteria) and ymCitrine (yeast). Binarized contiguous events were made solid by "filling" regions, and region properties were calculated via MATLAB's regionprops function: for yeast, event areas and bounding boxes (minimum quadrangle that would fit event); for bacteria, centroids (center coordinates of each event). Yeast event areas less than 10 px^2 were filtered out as noise, and images with fewer than three such events, fewer than three bacteria, or more than 1000 yeast events were ignored. In this way, we only analyzed events that were truly yeast, filtered out samples with too sparse or too concentrated cells, and corrected for optical defects which occasionally appeared as apparent single bright spots in a channel.

We calculated coincident bacteria by comparing red-channel centroids and yellow-channel bounding boxes. For each yeast event bounding box, we queried all bacterial positions for that image for inclusion in the bounding box, with detectably proximal bacterial getting saved as "coincident" per each yeast event. Importantly, this technique depends on distinguishing individual bacterial cells to be counted accurately, which at 10x objective, probably isn't always the case, and so the coincident bacteria metric is likely a slight undercount in all cases.

To impute the number of yeast cells in each ymCitrine event, for use in modeling, we estimated a baseline single-cell size based on mannose-supplemented day 1 data. Because many yeasts in clumps appear rather small, we chose a value on the low end of the single-yeast distribution: 20 px^2. We categorized each yeast event per image into number of predicted yeasts based on the square multiplier of this baseline area. E.g. events less than $2^2$*20 px^2 were considered one cell, events greater $2^2$*20 px^2 but less than $3^2$*20 px^2 considered two cells, etc. Because of variability in yeast cell sizes, and because images don't account for yeast stacking in the z-plane (i.e. any cells obscured by those at the bottom of a plate well), we expect these yeast counts to mostly be underestimates. We defined clumps as yeast events ≥ two cells, and numbers of clumps and numbers of clumped cells, along with their frequencies per total yeast, were calculated from that designation. Similarly, we determined coincident bacteria per clump by comparing coincident bacteria to whether a yeast event was categorized as a clump.

#### Colony analysis

We imaged colonies with the AxioZoom V16 dissecting microscope at 8x objective, using filter cubes for mCherry and ymCitrine for *cis*-donor mixed colonies; we additionally imaged *trans*-donor mixed colonies with a DAPI cube (see Methods). We converted output .czi files, each of which included all fluorescence images, to .tif files via a custom ImageJ macro with no brightness correction. We wrote a custom MATLAB scripts to import file information, read .tif files, background-correct images, identify colony circles, scan intensities of each channel, calculate radial metrics, and measure Li colocalization.

For red (bacteria) and yellow (yeast) channels, we binarized binarized via Otsu's method[1]. The blue channel corresponding to IDC reporter yeBFP was imported using adaptive sensitivity of 0.5. We used "filled" versions of bacterial and yeast channels—for which colony interiors were uniformly filled in—to automatically determine circular colony dimensions. We added together filled, binarized channels and calculated region properties for the combined binary image, as well as binarized bacterial and yeast samples, using MATLAB's regionprops function. If we determined none of the three "images" to have an event with length greater than 300 px and less than 1500 px, they were filtered out and defined as not having grown. Otherwise, we chose whichever of the three showed the largest diameter for circle definition, based on its centroid and major axis length. We visually screed all circle definitions, and any images requiring circle modification—either due to inaccurate circles or lack of circles—we manually drew and saved.

We background corrected images by setting all pixels beyond the boundary of the circle definition to "nan". We then used background corrected images to generate colony-wide metrics, e.g., averages for each channel across the colony. We calculated Li colocalization[2] intensity correlation quotient (ICQ) colony-wide by the formula

$$ICQ = \frac{\sum_{N>0} (B_i - \bar{B})(Y_i - \bar{Y})}{\sum_{N} (B_i - \bar{B})(Y_i - \bar{Y})} - 0.5$$

Where $X_i$ pertains to individual pixel intensities ($B$ = bacteria, $Y$ = yeast), from which we subtracted the mean intensity for that channel. For each pixel, we calculated a product of each channel's difference from the mean, and took the sum of these products. ICQ calculates the proportion of these products that are positive (N>0), based on the idea that, for random distributions of intensities in each channel, the sum of products of differences should near zero, and thus how positive the sum is corresponds to how colocalized the two channels are. If both channels deviate from the mean similarly in space, more products of differences will be positive, and ICQ will be higher. Conversely, if the channels are highly segregated—e.g., if the bacterial signal is well above its average at the same location that the yeast signal is well below its average—more products will be negative, and the ICQ will be lower. The subtraction of 0.5 is an arbitrary way to get the metric to straddle 0 (corresponding to "random" distributions), with negative values corresponding to mostly segregated channels (-0.5 = perfectly segregated), and positive values mostly colocalized (0.5 = perfectly colocalized). In our case, all colonies presented as mostly segregated—ranging from -0.5 to -0.2—which agrees with both subjective assessment of the images and dynamics experiments that demonstrate strong antagonism between the species, which should result in spatial exclusion.

We also calculated metrics radially, by defining a circumference along the colony circle's outer edge and taking intensity profiles for each unique radius from the circle center to pixels along the circumference. This yielded a matrix of pixel values in terms of radius $r$ angle $\theta$, but rings closer to the center were overrepresented due to being measured as many times as there were circumference pixels. Thus, we introduced "nan" values in proportion to that $r$'s fraction of the total radius $R$. Then we calculated metrics along each radial ring, including mean intensities and radial ICQ. Plots of these metrics showed that, for most cases, bacterial growth was localized to the outer ring of the colony, where the greatest abundance of nutrients could be found[3], while most of the yeast intensity was found within these bacterial boundaries (Fig S13). This trend was subverted somewhat by $S_{cross}$-pairings in low amino acid concentrations, for which bacteria were better able to grow within the colony center apparently due to lowered occupancy of the yeast. Because yeast signals mostly dropped off near the edge of the colony, where bacterial signals rose, the greatest colocalization between the channels could be found near the edge, too, though the exact radius of this spike depended on the conditions, with more divergence at lower amino acids, and the highest amino acid concentrations all showing similar colocalization at $r \sim= 3$mm.

From this, we might infer that IDC occurred primarily near the border of colonies, especially for more segregated populations. Indeed, in one case this was observed explicitly, with a subpopulation of transconjugants detectable fluorescently via the BFP channel (Fig S14). This subpopulation seemed to grow from the bacterial-dense outer ring and extend beyond it, and the colony's IDC counts corresponded to a "jackpot" IDC event (IDC $\geq$ 500 CFU)[4]. In general, the *trans*-IDC plasmid carrying yeBFP ("IDC reporter") proved unusable in most experiments due to its low signal, high autofluorescence, and infrequent IDC events. In colonies, the blue signal was indistinguishable from a mixture of bacterial and yeast autofluorescence, with the former appearing stronger; thus, in general, the blue signal was brightest wherever the red signal was brightest. In this case, however, it's clear that the bright blue subpopulation along the outer edge didn't align with the red signal, and thus was a true transconjugant population. More work on this reporter scheme might unlock the possibility of better tracking IDC events spatially and allow much better tracking of IDC distribution.

### Modeling

#### "Free" (non-clumping) growth model

We derived growth equations for bacterial and yeast growth from Pearl-Verhulst[5] logistic growth, for which cells' growth was determined by its (monoculture) growth rate $R$, carrying capacity $K$, and death rate $D$. Modifications for coculture conditions included deviations from monoculture carrying capacity term and amino acid secretion terms[6,7]. For the effect of one species on limiting the carrying capacity of the other, we used a multiplier $c$ to account for incomplete ecological niche overlap. The global concentration of amino acid supplemented to media $G$ and amino acid secretion $\alpha$—dependent on secreting cell's concentration—together modified the growth rate of each species. Monod term $k$ determined a strain's susceptibility to amino acid changes.

Transconjugants grew similarly to non-conjugated yeast, and thus had a nearly identical growth equation. Transconjugants were also added to the system by bacterial and yeast collisions, as modified by IDC rate term $\gamma$. Note that because we compared the growth equation for yeast $dY/dt$ to yeast fluorescence data (see below), $Y$ was a representation of *all yeast*, including transconjugants, and thus had no term depleting cells proportionate to IDC rate $\gamma$, as is occasionally seen in other equations[8–10].

Growth equations, for bacteria ($B$), yeasts ($Y$), and transconjugants ($T$)

$$\frac{dB}{dt} = R_b * B\left(\frac{\alpha_b Y + G_b}{\alpha_b Y + G_b + k_b}\right)\left(1 - \frac{B}{K_b} - \frac{c_y Y}{K_y}\right) - D_b\left(\frac{B}{K_b}\right)$$

$$\frac{dY}{dt} = R_y * Y\left(\frac{\alpha_y B + G_y}{\alpha_y B + G_y + k_y}\right)\left(1 - \frac{Y}{K_y} - \frac{c_b B}{K_b}\right) - D_y\left(\frac{Y}{K_y}\right)$$

$$\frac{dT}{dt} = R_y * T\left(\frac{\alpha_y B + G_y}{\alpha_y B + G_y + k_y}\right)\left(1 - \frac{T}{K_y} - \frac{c_b B}{K_b}\right) - D_y\left(\frac{T}{K_y}\right) + \gamma\left(\frac{B * Y}{B + Y}\right)$$

We first estimated growth rates R and carrying capacities K by fitting integrated versions of simplified monoculture growth equations (lacking amino acid terms, death rates, and coculture modifications) to monoculture fluorescence data at 100% amino acids in solution. Global amino acid concentrations G were known (see Methods for molar values), and we based initial guesses for cell secretion α on literature values for similar strain mutants—0.022 gTrp/CDW*hr for ΔtrpR bacteria[11], and 7*105 moleculesLeu/cell*s for Leu++ yeasts[3]—taken for 1 hour, with an assumed bacterial CDW = 3*10-13 g/cell. Using these, we arithmetically derived Monod terms k for crossfeeder monoculture fits using the equation Rmax = R * [AA]/([AA] + k), where [AA] = limited amino acid concentration, over a range of supplemented values.

For all subsequent terms and fits, we used Latin Hypercube Sampling (LHS) to fit a MATLAB ODE solver. We set upper and lower bounds for each parameter, between which 100,000 random parameter guesses were generated, each of which was put through the ODE solver. Each model output, determined by that guess's randomized parameters, was compared to fluorescence data, error between the two was calculated, and random model guesses were ranked by lowest calculated error.

We then estimated niche overlap terms c using fluorescence of WT cocultures at 100% amino acids, for each WT donor variant (plasmids carried). Original parameter ranges were set widely for simplified conditions—e.g. c was initially sampled between -1 and 2 for WT pairings (Fig S8)—then tightened for full-model fitting.

For batch coculture fitting, we compared fluorescence values to a modified version of ODE solver, in which we fed model outputs from each day into the next day's initial conditions. In this way, we were able to emulate batch culture dilutions at the times experimentally performed and keep the ODE solver in time units of hours. We ignored day 1 growth for both measurements and model, as variation in cell counts among strains initially adapting to batch culture conditions proved extremely unpredictable and resulted in errant model fits. To allow for experimental variation, we incorporated noise into day-end outputs at a rate of +/- 20%, before feeding them fed back into the solver as next-day initial conditions. Initially, we only calculated error between model and measured data for bacterial and yeast growth signals (not IDC values), and we ranked fits according to those errors. We manually assessed means of each model's 1000 best fits (top 1%) for each cell pairing based on three criteria: 1) susceptibility of each strain to changes in amino acid concentration, 2) steady-state persistence of each strain, and 3) approximate D:R ratio of cells. We manually modified means to best reflect these priorities, and then fed them back into the solver. We repeated this process several times, until model outputs no longer improved upon experimental matches (Fig S9). Model thus represents a local minimum for fitting parameters and is not meant to represent a unique fit.

Once a representative parameter set was acquired for each species' growth in coculture, we fixed all parameters besides IDC rate γ, and we tested γ across a range of possible outcomes (10-6 – 10-2) for each cell pairing and concentration of added leucine and tryptophan. Note that, unlike bacterial and yeast growth equations, each of which maintains a single fluorescence unit (mCherry or ymCitrine), transconjugant equations include a combination of each fluorescence type. To account for this, the transconjugant model converts each cell type from fluorescence to cell count before multiplying by γ—using conversion factors derived by comparing flow cytometry data (cell counts) to fluorescence data—after which the entire frequency term is multiplied again by ymCitrine/cell, to maintain units of ymCitrine for transconjugants. Because transconjugants were modeled in units of ymCitrine, and because half of each growth well's volume was plated for IDC counts, we divided model outputs for IDC by 2*Citrine/cell. We incorporated averages from each condition's IDC data into IDC-sweep plots by altering heatmap colormaps according to where color (row-fraction of total output range) matched IDC counts.

Table S1: Free-cell model parameters

| Var | Parameter | Unit | Model Fits per Cell Pairing | | | |
|---|---|---|---|---|---|---|
| | | | $E_{cross}$ $S_{cross}$ | $E_{cross}$ S | E $S_{cross}$ | E S |
| $R_b$ | Bacterial growth rate | $hr^{-1}$ | 0.75 | 0.66 | 0.75 | 0.75 |
| $R_y$ | Yeast growth rate | $hr^{-1}$ | 0.60 | 0.59 | 0.58 | 0.58 |
| $K_b$ | Bacterial carrying capacity | mCherry | 3500 | 2987 | 3201 | 3500 |
| $K_y$ | Yeast carrying capacity | ymCitrine | 2600 | 3044 | 3039 | 2998 |
| $c_b$ | Ecological niche overlap (effect of bacteria on yeast) | Unitless | 0.80 | 0.84 | 0.80 | 0.80 |
| $c_y$ | Ecological niche overlap (effect of yeast on bacteria) | Unitless | 0.95 | 0.99 | 0.90 | 0.90 |
| $G_b$ | Global amino acid (for dependent bacteria) | Molar | 100% leucine = $7.622*10^{-4}$ M | | | |
| $G_y$ | Global amino acid (for dependent yeast) | Molar | 100% tryptophan = $2.449*10^{-4}$ M | | | |
| $\alpha_b$ | Secreted amino acid (for dependent bacteria) | Molar/ mCitrine | 1E-4 | 1E-10 | 1E-4 | 1E-10 |
| $\alpha_y$ | Secreted amino acid (for dependent yeast) | Molar/ mCherry | 1E-9 | 1E-9 | 1E-12 | 1E-12 |
| $k_b$ | Monod term for dependent bacteria | Molar | 2E-6 | 2E-6 | 0 | 0 |
| $k_y$ | Monod term for dependent yeast | Molar | 2.5E-5 | 0 | 2.5E-5 | 0 |
| $D_b$ | Bacterial death rate | $hr^{-1}$ | 0.455 | 0.455 | 0.455 | 0.455 |
| $D_y$ | Yeast death rate | $hr^{-1}$ | 0.50 | 0.547 | 0.570 | 0.498 |
| $\gamma$ | IDC rate | Unitless | 4E-5 | Unknown | 7E-6 | Unknown |

#### Clumping growth model

After testing the free cell model against experimental data, we modified deterministic equations to capture aggregation ("clumping") dynamics between the species. Specifically, three equations were added to the system, tracking 1) formation of clumps, as determined by some constant rate of aggregation for every random free-cell collision; 2) total clumped bacteria, based on both growth of already-clumped bacteria, or additional collisions between clumps and free bacteria; 3) total clumped yeast, based on growth of already-clumped yeast or additional collisions between clumps and free yeast. The latter clumped-cell growth equation terms were similar to free-cell growth, except for modified growth rates (starting values ~1/3 free cell values) and proximity terms P, which allow for altered amino-acid feeding from opposing cell type in a clump. We additionally modified free-cell growth equations to be carrying capacity-limited by summing clumped and free cells in the total per species. Finally, we modified the transconjugant equation to include a second IDC rate γc, based on total number of clumped-bacteria and clumped-yeast interactions. We also changed the growth rate for transconjugants to the clumped-yeast rate, to reflect the expectation that most transconjugants require clumping at some point.

<u>Growth equations, for bacteria (*B*), yeasts (*Y*), transconjugants (*T*), total clumps (*C*), clumped bacteria ($C_b$), and clumped yeast ($C_y$)</u>

$$\frac{dB}{dt} = (R_b * B)\left(\frac{\alpha_b Y + G_b}{\alpha_b Y + G_b + k_b}\right)\left(1 - \frac{(B + C_b)}{K_b} - \frac{c_y(Y + C_y)}{K_y}\right) - D_b\left(\frac{B}{K_b}\right)$$

$$\frac{dY}{dt} = (R_y * Y)\left(\frac{\alpha_y B + G_y}{\alpha_y B + G_y + k_y}\right)\left(1 - \frac{(Y + C_y)}{K_y} - \frac{c_b(B + C_b)}{K_b}\right) - D_y\left(\frac{Y}{K_y}\right)$$

$$\frac{dT}{dt} = R_{cy} * T\left(\frac{\alpha_y B + G_y}{\alpha_y B + G_y + k_y}\right)\left(1 - \frac{Y + C_y}{K_y} - \frac{c_b(B + C_b)}{K_b}\right) - D_y\left(\frac{T}{K_y}\right) + \gamma\left(\frac{B * Y}{B + Y}\right) + \gamma_c\left(\frac{C_b * C_y}{C_b + C_y}\right)$$

$$\frac{dC}{dt} = R_c\left(\frac{B * Y}{B + Y}\right)$$

$$\frac{dC_b}{dt} = (R_{cb} * C_b)\left(\frac{P_b\alpha_b Y + G_b}{P_b\alpha_b Y + G_b + k_b}\right)\left(1 - \frac{C_b}{K_b} - \frac{c_y C_y}{K_y}\right) - D_b\left(\frac{C_b}{K_b}\right) + R_c\left(\frac{C * B}{C + B}\right)$$

$$\frac{dC_y}{dt} = (R_{cy} * C_y)\left(\frac{P_y\alpha_y B + G_y}{P_y\alpha_y B + G_y + k_y}\right)\left(1 - \frac{C_y}{K_y} - \frac{c_b C_b}{K_b}\right) - D_y\left(\frac{C_y}{K_y}\right) + R_c\left(\frac{C * Y}{C + Y}\right)$$

We fit clumped-cell dynamics similarly to free-cell equations, using LHS and the ODE batch solver. We additionally modified model outputs in two ways. We limited clumped-bacteria outputs to 10 bacteria per clump (based on image analyses, see above), beyond which clumped bacteria counts were subtracted from clumped-bacteria model outputs for each day and added to free-bacteria counts, before the next day's growth was modeled. We also added final clumped-cell model outputs to total free-cell outputs before comparing to fluorescence data, which doesn't distinguish between the two.

To use image analyses of batch culture clumping as comparative data for the model, dynamic information on number of clumps, total clumped bacteria, and total clumped yeast were acquired by the following means (see Image Analysis for determination of clump numbers and number of cells). The total number of clumps was taken as a fraction of all yeast, based on determination of number of yeast in each clump. Thus, a fractional term of clumps/total yeast could be multiplied by the ymCitrine fluorescence signal to get the number of clumps in units of ymCitrine, regardless of the fraction of cell culture was imaged to determine clump count. The number of clumped yeast was calculated similarly, but tracking clumped yeast per total yeast, and multiplying by total yeast ymCitrine signal. For clumped bacteria, coincident bacteria on all clumped yeast (>2 yeast per event) was taken as a fraction of total clumps, to get clumped bacteria per clumps. Upon deriving number of clumps in terms of ymCitrine, this could be multiplied by clumped bacteria per clumps term to get clumped bacteria in terms of ymCitrine, which was then converted to mCherry by conversions described in the discussion of IDC sweeps for the free-cell model. All three metrics—each only determined at day-ends—were imputed for intermediate times as linear increases from $1/10^{th}$ the previous day's metric (day 1 assumed = 0).

We compared batch coculture LHS model outputs similarly to free-cell LHS fitting, using error between the model and experimental data, but here we calculated error between each experimental result: total bacteria, total yeasts, IDC counts, number of clumps, clumped bacteria, and clumped yeast (Fig S11). Upon deriving fits that recapitulated main experimental outcomes, we swept key clumping parameters P and γc across a range of values to find approximate viable values. IDC rate sweeps of $\gamma$ and $\gamma_c$ yielded several values of each that were able to recapitulate the data, though they roughly fell into two categories: low $\gamma_c$ with $\gamma$ in the range of $3*10^{-4} – 6*10^{-4}$, or low $\gamma$ with $\gamma_c$ in the range of $2*10^{-5} – 4*10^{-5}$. *P* value sweeps show an apparent amino-acid secretion increase on the order of 50x from S, for $E_{cross}$ cells to be able to grow in 0% leucine. While this model assumes that the vast majority of transconjugants result from clumped interactions, it's not clear how transient clumps are, so the free-cell IDC transfer rate here accounts for IDC from cells measured to be "free" despite having previously been clumped at some time between measurements.

Table S2: Clumped-cell model parameters

| Var | Parameter | Unit | Model Fits per Cell Pairing | | | |
|---|---|---|---|---|---|---|
| | | | $E_{cross}$ $S_{cross}$ | $E_{cross}$ S | E $S_{cross}$ | E S |
| $R_b$ | Free bacterial growth rate | $hr^{-1}$ | 0.75 | 0.75 | 0.75 | 0.75 |
| $R_y$ | Free yeast growth rate | $hr^{-1}$ | 0.58 | 0.58 | 0.58 | 0.58 |
| $R_{cb}$ | Clumped bacterial growth rate | $hr^{-1}$ | 0.25 | 0.25 | 0.25 | 0.25 |
| $R_{cy}$ | Clumped yeast growth rate | $hr^{-1}$ | 0.18 | 0.18 | 0.18 | 0.18 |
| $R_c$ | Clumping rate | Unitless | 0.03 | 0.05 | 0.03 | 0.05 |
| $K_b$ | Bacterial carrying capacity | mCherry | 7566 | 7269 | 5590 | 6790 |
| $K_y$ | Yeast carrying capacity | ymCitrine | 2822 | 2700 | 2637 | 2412 |
| $c_b$ | Ecological niche overlap (effect of bacteria on yeast) | Unitless | 0.80 | 0.69 | 0.90 | 0.70 |
| $c_y$ | Ecological niche overlap (effect of yeast on bacteria) | Unitless | 0.91 | 0.95 | 0.90 | 0.93 |
| $G_b$ | Global amino acid (for dependent bacteria) | Molar | 100% leucine = $7.622*10^{-4}$ M | | | |

| | | | | | | |
|---|---|---|---|---|---|---|
| $G_y$ | Global amino acid (for dependent yeast) | Molar | 100% tryptophan = 2.449*10$^{-4}$ M | | | |
| $\alpha_b$ | Secreted amino acid (for dependent bacteria) | Molar/ mCitrine | 9.2E-5 | 1E-8 | 9.2E-5 | 1E-8 |
| $\alpha_y$ | Secreted amino acid (for dependent yeast) | Molar/ mCherry | 1E-9 | 1E-9 | 1E-12 | 1E-12 |
| $P_b$ | Proximity multiplier for $\alpha_b$ | Unitless | 50 | 50 | 1 | 1 |
| $P_y$ | Proximity multiplier for $\alpha_y$ | Unitless | 1 | 1 | 1 | 1 |
| $k_b$ | Monod term for dependent bacteria | Molar | 2E-6 | 2E-6 | 0 | 0 |
| $k_y$ | Monod term for dependent yeast | Molar | 1.2E-5 | 0 | 1.2E-5 | 0 |
| $D_b$ | Bacterial death rate | hr$^{-1}$ | 0.46 | 0.50 | 0.52 | 0.46 |
| $D_y$ | Yeast death rate | hr$^{-1}$ | 0.46 | 0.39 | 0.45 | 0.48 |
| $\gamma$ | Free IDC rate | Unitless | ~ 2E-4 – 1E-3 (see Fig 4a, SF10) | | | |
| $\gamma_c$ | Clumped IDC rate | Unitless | ~1E-4 – 6E-4 (see Fig 4b, SF12) | | | |

#### Rescue growth model

To apply insights from the clumping model, rescue conditions were tested within the clumping model via the following modifications. Here, yeast cells are selected for IDC events, and there's no limitation of tryptophan (required externally by $S_{cross}$). To account for this experimental change, the entire amino acid term for yeast growth was based upon limitations of uracil and histidine (required externally for both yeast strains, and carried by MOB1 plasmid)—a change accounted for in changes to input parameters *G*—but the entire amino acid term was removed from the transconjugant ODE, making them agnostic to terms *G*, *α*, and *P*. Moreover, because we assume rescued yeast to not stay primarily clumped over a long period, the growth rate for transconjugants was assumed to fall somewhere between that of clumped yeast and free yeast. Finally, unlike previous transconjugant equations, in which transconjugants were primarily carrying-capacity limited by non-transconjugant yeast, here the opposite is likelier true, for any rescue conditions (that would allow yeast numbers to approach carrying capacity), so *T* is entered into carrying capacity limitation for free bacteria, free yeast, and transconjugants.

Growth equation for transconjugants (*T*) (other equations match those from clumped model)

$$\frac{dT}{dt} = \left(\frac{R_{ly} + R_y}{4}\right) * T\left(1 - \frac{(Y + T + C_y)}{K_y} - \frac{c_b B}{K_b}\right) - D_y\left(\frac{T}{K_y}\right) + \gamma\left(\frac{B * Y}{B + Y}\right) + \gamma_c\left(\frac{C_b * C_y}{C_b + C_y}\right)$$

To model the phase space of transconjugant outcomes for a range of bacterial and yeast fitness, based on limited amino acids, concentrations of leucine ($E_{cross}$-dependent) and uracil/histidine (all yeast-dependent) *G* were tested against other fixed parameter outcomes from the clumped-cell model. Because molar concentrations of uracil ($S_{cross}$-dependent) and histidine (S-dependent) are similar for 100% KS solution (0.0954 mM histidine, 0.178 mM uracil), both were assumed equal at 100% (0.100 mM). Monod terms *k* were modified to reflect amino-acid sensitivity differences from crossfed coculture experiments, with $E_{cross}$ maintaining its value for leucine dependence, E maintaining its lack of sensitivity ($k_b = 0$), and setting both yeast strains to the value found for $S_{cross}$ sensitivity to tryptophan (see Table S2), making both $S_{cross}$ and S equally sensitive to U or H.

Concentrations of amino acids were swept over the range of 0-15%, as per many experimental conditions, even though the rescue assay kept [L] at 0% (uracil and histidine ranged from 0% to 15%). Model anomalies arose when setting all amino acids at or near 0, in which IDC values far surpassed possible ranges (>$10^{10}$), presumably due to small denominators in cell collision equation terms. To account for this, model outputs for which either bacterial or yeast counts dropped below 1 (after converting from fluorescence, for both clumped- and free-cells) were zeroed out for IDC at those times. This modification had no perceptible changes for amino acids not near 0%.

Table S3: Rescue model parameters (those not listed are the same as in Table S2)

| Var | Parameter | Unit | Model Fits per Cell Pairing | | | |
|---|---|---|---|---|---|---|
| | | | $E_{cross}$ $S_{cross}$ | $E_{cross}$ S | E $S_{cross}$ | E S |
| $G_y$ | Global amino acid (for dependent yeast) | Molar | 100% Uracil/Histidine = 1.0*$10^{-4}$ M | | | |
| $\alpha_y$ | Secreted amino acid (for dependent yeast) | Molar/ mCherry | 1E-20 | 1E-20 | 1E-20 | 1E-20 |
| $\gamma$ | Free IDC rate | Unitless | 1E-8 | | | |
| $\gamma_c$ | Clumped IDC rate | Unitless | 2.5E-4 | | | |

#### Strains and Plasmids

Table S4: Strains used in this study

| ID | Species | Genetic features | Fluorescence | Source | Figures |
|---|---|---|---|---|---|
| yMM1585 | Yeast | $Leu^{++}$, $Trp^{-}$, $Ura^{-}$ | ymCitrine, yCerulean | [3], this work | 2,5,6,SF1-4, |
| yMM1636 | Yeast | $His^{-}$ | ymCitrine | [3], this work | All but 7, SF17-20 |
| yMM1720 | Yeast | $Leu^{++}$, $Trp^{-}$, $His^{-}$ | ymCitrine, yCerulean | [3], this work | 3,4,SF3-12 |
| yMM1786 | Yeast | $Leu^{++}$, $Trp^{-}$, $Ura^{-}$, $His^{-}$ | ymCitrine, yCerulean | [3], this work | 7,SF17-20 |
| kMM0011 | Bacterium | None | None | [12] | All but 7, SF17-20 |
| kMM0127 | Bacterium | $Trp^{++}$, $Leu^{-}$ | None | [12], this work | All |

Table S5: Plasmids used in this study

| ID | Species | Function / Features | Fluorescence | Source | Figures |
|---|---|---|---|---|---|
| pMM0819 | Bacterium | pProD-mCherry | mCherry | Addgene 87144 | All |
| pMM0820 | Bacterium | $\lambda_{red}$ genes | None | [13] | None |
| pMM0821 | Bacterium | Flippase | None | [13] | None |
| pMM0892 | Bacterium | T4SS genes, $Gent^{R}$ | None | pTA-Mob 1.0[14] | 3,4,7,SF3-14,17-20 |

| | | | | | |
|---|---|---|---|---|---|
| pMM0893 | Both | T4SS genes, Gent$^R$, URA3, HIS3, CEN/ARS, $ori^T$ | None | pTA-Mob 2.0, Addgene 149662 | 1,2,5,6,SF1-4,15-16 |
| pMM1340 | Bacterium | sgRNA assembly vector, sfGFP | GFP | Addgene 90516 | None |
| pMM1341 | Bacterium | Cas9, sfGFP | GFP | Addgene 90519 | None |
| pMM1342 | Bacterium | sgRNA assembly, YTK target | None | This work | None |
| pMM1360 | Yeast | yeBFP, URA3, CEN/ARS | BFP | [15], this work | 7,SF17-20 |
| pMM1438 | Both | yeBFP, $ori^T$, HIS3, CEN/ARS | BFP | [15], this work | 3,4,SF3-14 |
| pMM1440 | Both | Cas9, $ori^T$, sgRNA (YTK), HIS3 | None | This work | 7,SF17-20 |
| pMM1354 | Both | Cas9, sgRNA (YTK), HIS3 | None | This work | 7,SF17-20 |

#### Statistics

See Methods and relevant figures for statistical methodology.

Table S6: 1-way ANOVA results for colony colocalization (ICQ, see Fig 5c)

| **LW%** | **Day** | ***F* value** | ***p* value** | **df** |
|---|---|---|---|---|
| 100 | 1 | 7.3163 | 0.0017 | 3 |
| 100 | 2 | 3.6916 | 0.0290 | 3 |
| 100 | 3 | 1.2825 | 0.3075 | 3 |
| 100 | 4 | 6.0689 | 0.0041 | 3 |
| 100 | 5 | 1.8477 | 0.2168 | 3 |
| 100 | 6 | 0.5891 | 0.6392 | 3 |
| 0 | 1 | 28.0473 | $2.29*10^{-7}$ | 3 |
| 0 | 2 | 53.9723 | $9.0556*10^{-10}$ | 3 |
| 0 | 3 | 37.5045 | $2.1079*10^{-8}$ | 3 |
| 0 | 4 | 26.4395 | $3.6668*10^{-7}$ | 3 |
| 0 | 5 | 23.9480 | $2.3787*10^{-4}$ | 3 |
| 0 | 6 | 26.5237 | $1.6502*10^{-4}$ | 3 |

### Supplemental Figure Captions

#### SI Figure 1: Condition screening for batch cocultures

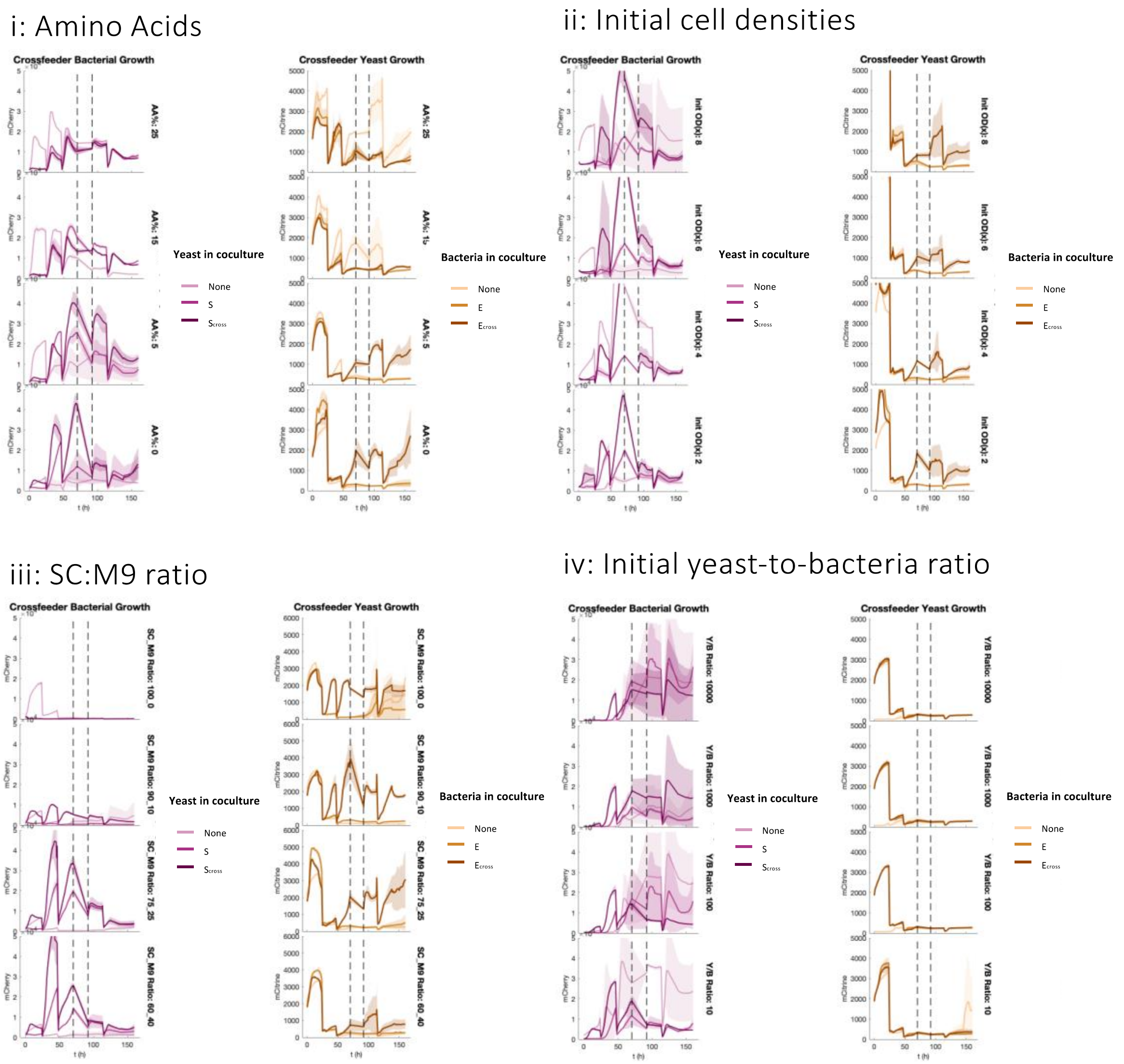


Fluorescence readings of crossfeeder bacteria (red traces) and crossfeeder yeast (brown traces) across several conditions during six days of batch culture. In each case, darkest traces pertain to coculture with paired crossfeeder strain, mid-darkness traces with WT pair, and lightest traces monoculture (e.g. dark red is bacteria grown with crossfeeder yeast, mid-red trace is bacteria grown with WT yeast, light red trace is bacteria grown in monoculture); lines are means of three replicates, shading is 95% CI. (i) Crossfeeding strains grown across several limited amino acid concentrations (LW%, rows). (ii) Initial total cell density (rows), all 1:1 ratio between cell types, 0% LW. (iii) Ratios of SC (yeast minimal media) to M9 (bacteria minimal media, rows), all 0% LW. (iv) Initial ratio of yeast to bacteria (rows), all at 0% LW. Across all conditions of 0% LW, 75:25 SC:M9 (iii, 2nd row) showed the most promise for survival of both crossfeeders by the end of time course (dark traces). Vertical dotted lines represent gap in data.

#### SI Figure 2: Growth trajectories for all *cis*-cell pairings at various amino acid concentrations

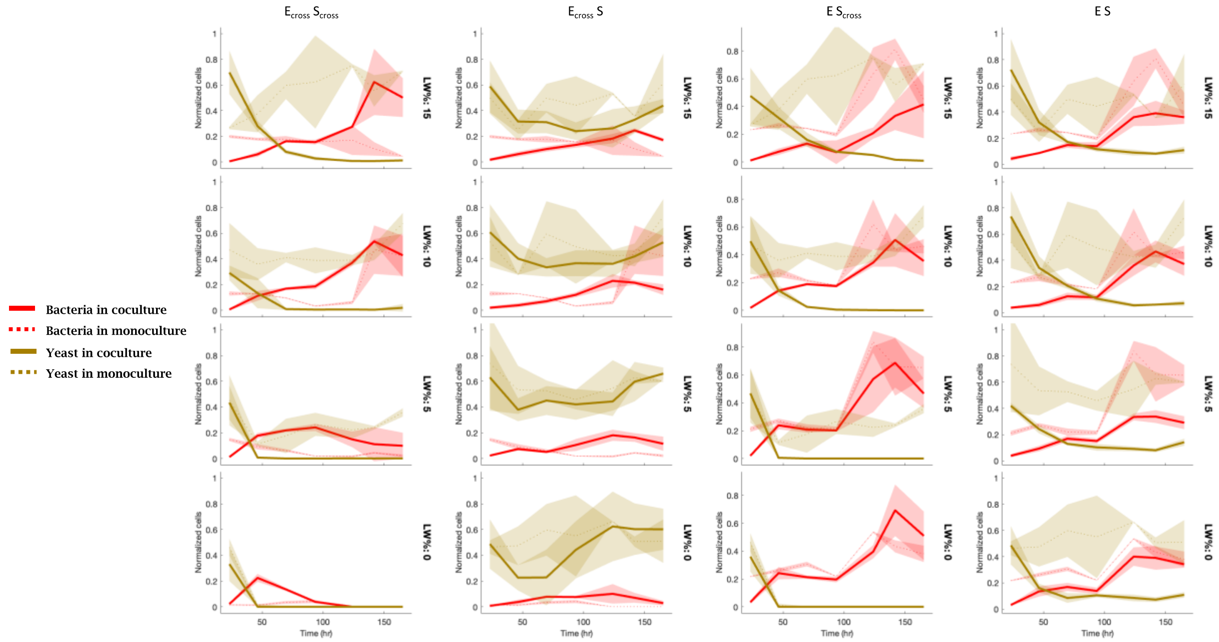


Normalized flow cytometry cell counts for each coculture pairing (columns) at four crossfed amino acid concentrations (LW%, rows), for seven days of batch culturing. Bacterial cell counts shown in red, yeast counts in yellow. Solid lines represent coculture traces, which are plotted with each cell's monoculture traces (dotted) for comparison; shading is standard deviation (four replicates coculture, two replicates monoculture). Representative outcomes at 0% LW range from apparent parasitism ($E_{cross}$ $S_{cross}$), commensalism ($E_{cross}$ S), and competitive exclusion (E $S_{cross}$), though the mechanistic causes of each interaction type wasn't specifically probed.

### SI Figure 3: D:R ratios vs. IDC counts

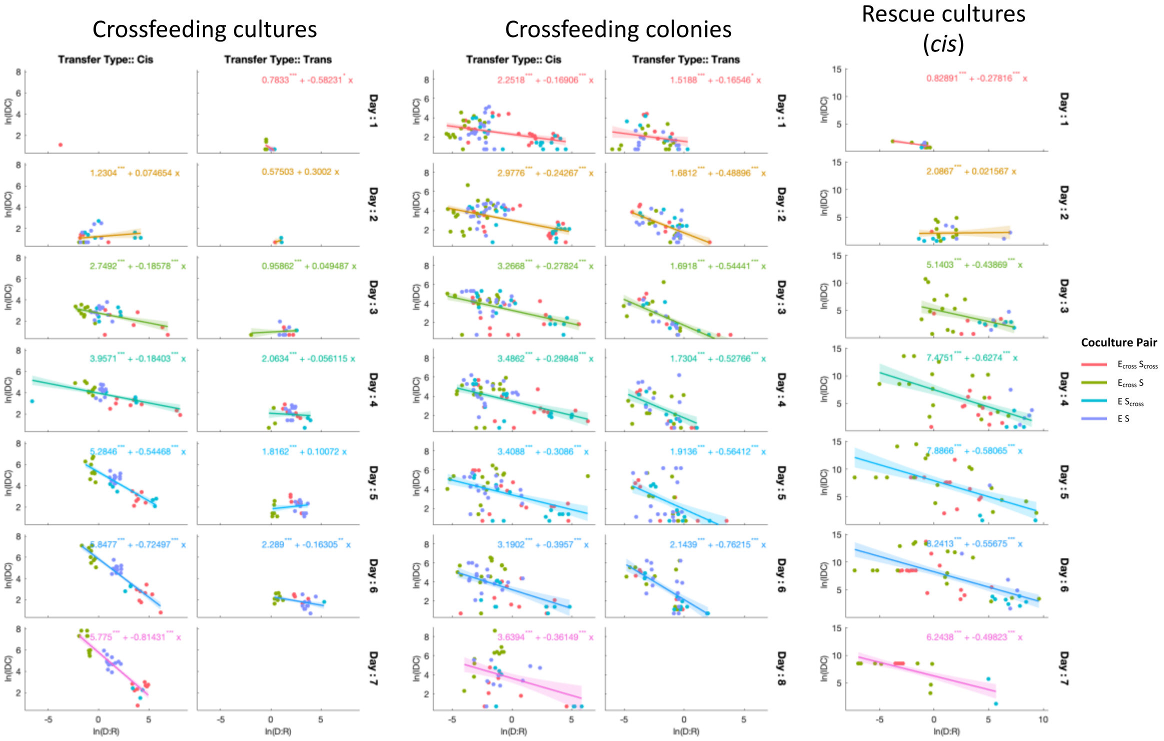


Log-log plots of donor-to-recipient ratios (D:R) and IDC, for *cis*- and *trans*-donor cultures (left 2 columns), *cis*- and *trans*-donor colonies (middle 2 columns), and rescue assay (*cis*, right column), for each day's measurements (rows). Generalized linear model fit, with normal distribution, shown in solid red line, with 95% CI in shaded region. Fit equation displayed with stars denoting *p*-value significance for each term. Note that while most conditions and days show a similar negative correlation between D:R and IDC, *trans*-cultures don't explicitly follow this trend, possibly due to low IDC near the detection limit.

#### SI Figure 4: Transconjugants as fractions of yeast populations over time

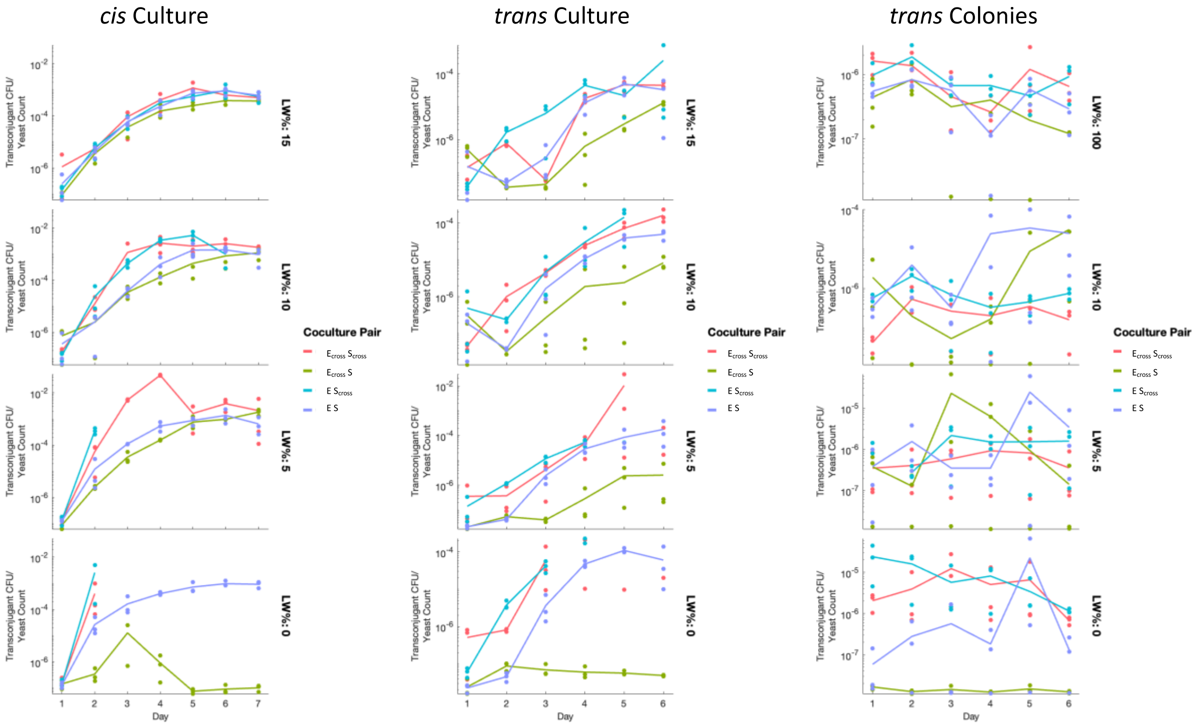


Fraction of transconjugants per total yeast for *cis*-donor cultures, *trans*-donor cultures, and *trans*-donor colonies over 6-7 days. Transconjugants measured as CFU per 100 μL culture or CFU per colony. Yeast counts from culture flow cytometry were back-calculated to represent count per 100 μL. Note that, for both culture examples, transconjugant fractions increase for all pairings until leveling off at ~day 5, suggesting that increased IDC for certain pairings is not solely a function of higher yeast populations. Also note that *trans*-donor pairings show lower IDC in general. Colony fractions, on the other hand, are mostly constant and noisy, which could either result from a spatial population-stabilizing effect (i.e. there are few new interactions over time that could result in additional IDC events) or because each colony is measured separately—e.g. day 4's results don't relate to day 5, etc.—and counts are highly variable due to jackpot events.

#### SI Figure 5: Clump sizes and coincident bacteria

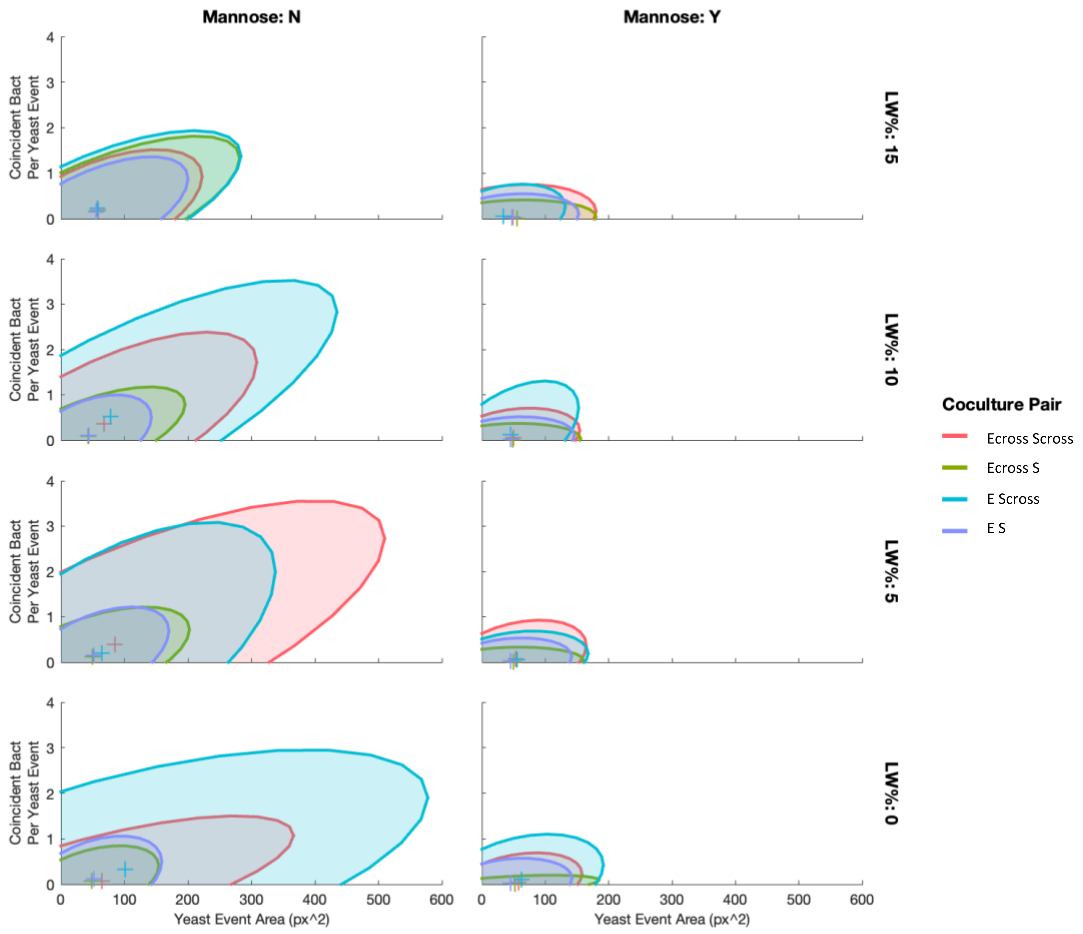


Sizes of optical yeast events, as determined by image analysis of ymCitrine fluorescence, plotted against number of coincident bacteria (number of distinct bacteria identifiable in proximity to yeast events) after six days of batch culture. Ellipses are fit to include 95% of points for each cell pairing (color) and LW% (rows), for samples without mannose (left) or with mannose (right). Colors pertain to coculture pairings.

#### SI Figure 6: Histograms of clump sizes and coincident bacteria

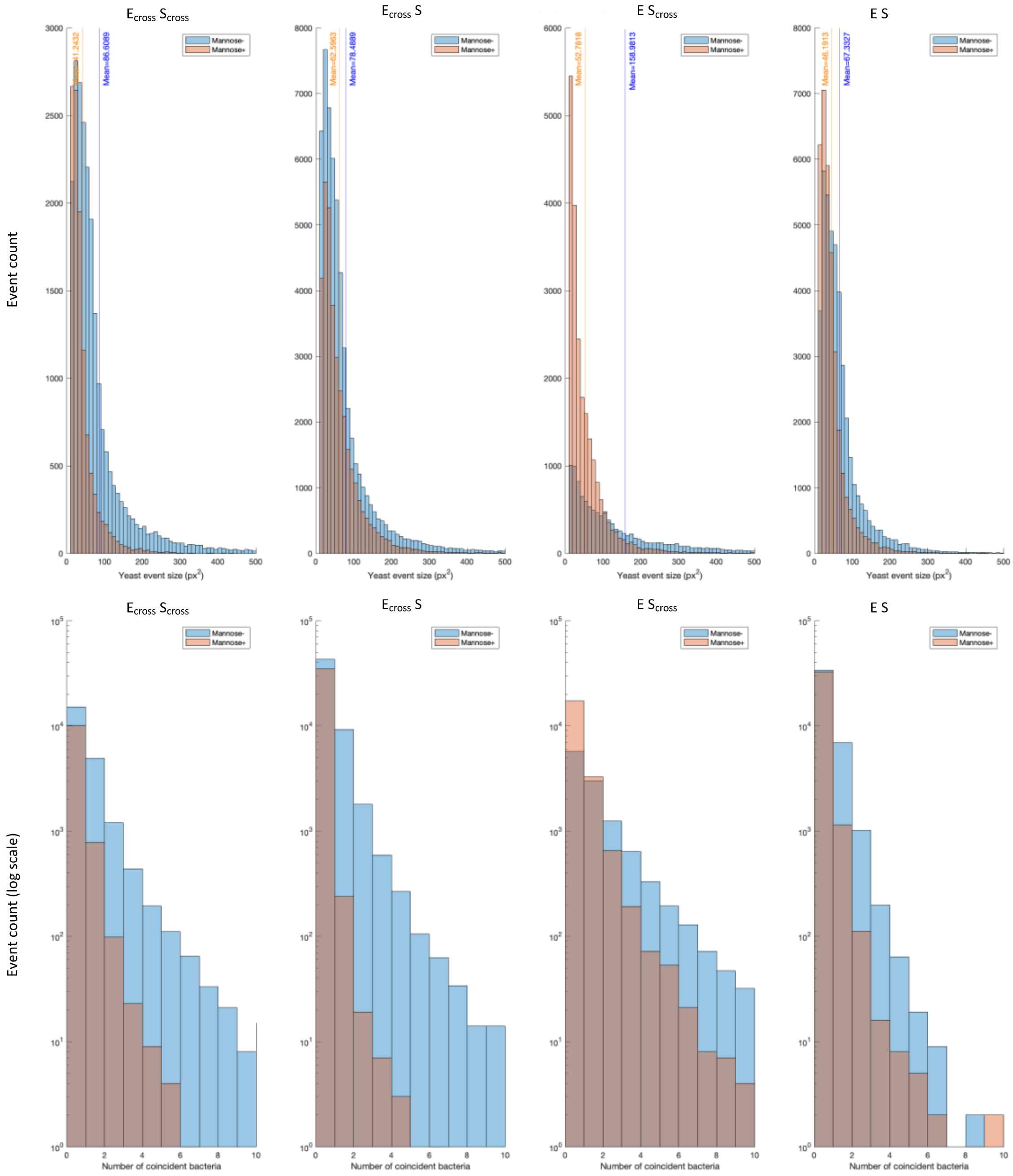


Histograms of optical yeast event sizes (top), as determined by image analysis of ymCitrine fluorescence, after six days of batch culture, either with mannose (orange) or without it (blue). Means of each distribution is shown in vertical lines. Log-frequency histograms of coincident bacteria (number of distinct bacteria identifiable in proximity to yeast events) shown at bottom for the same samples.

#### SI Figure 7: D:R and IDC difference in clumping

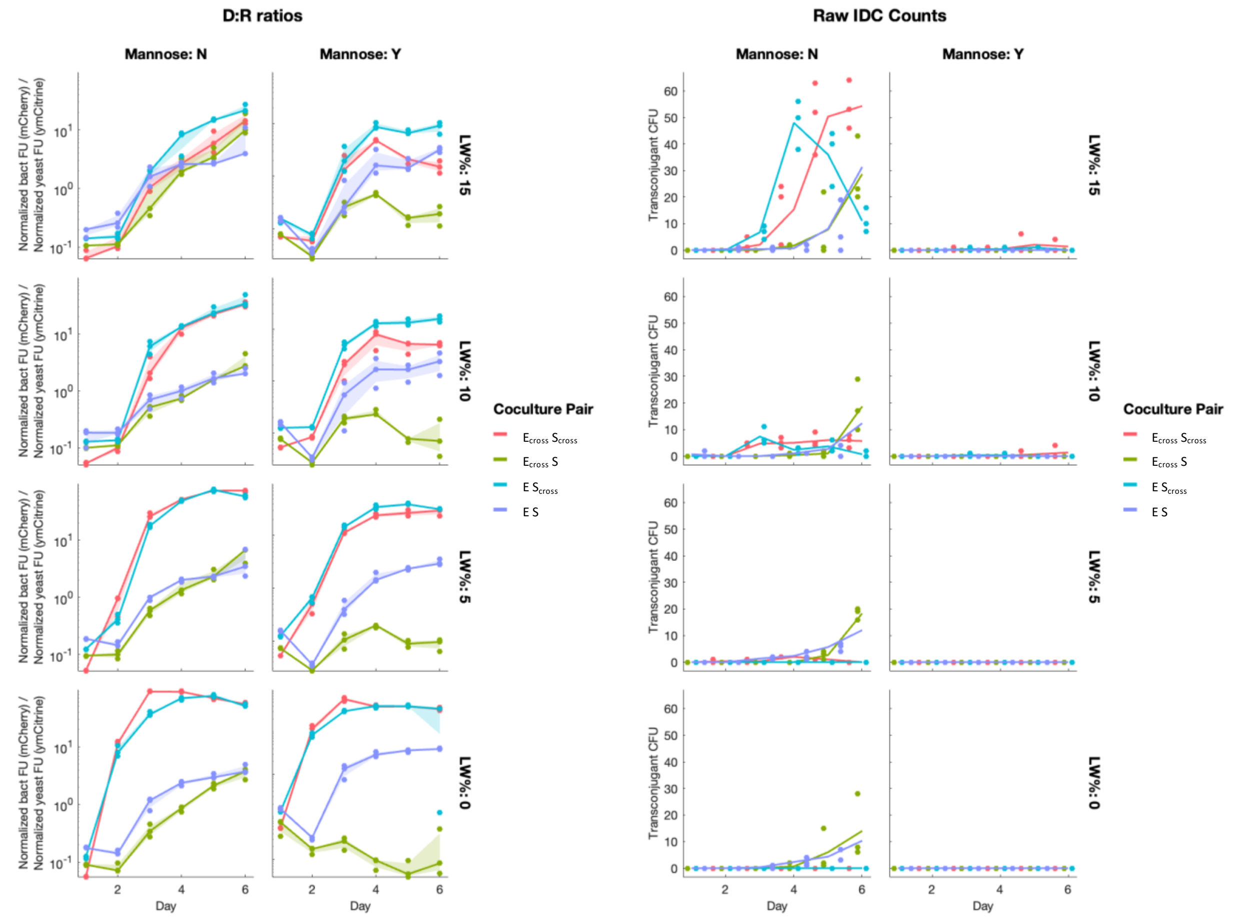


Donor-to-recipient ratios (left) for each cell pairing (color) and LW% (row), for samples with or without mannose (columns) over six days of batch culture. Note how the $E_{cross}$-S pairing (green) ratio stays fairly consistent without mannose—wherein cells can clump—but drops with LW% in samples with mannose supplemented, preventing clumping. IDC counts (right) for the same conditions show markedly higher conjugation activity when cells are allowed to clump (mannose-minus, left column). Lines represent means of three replicates, shaded regions in D:R plots standard deviation.

#### SI Figure 8: Example parameter distributions for niche overlap terms *c*

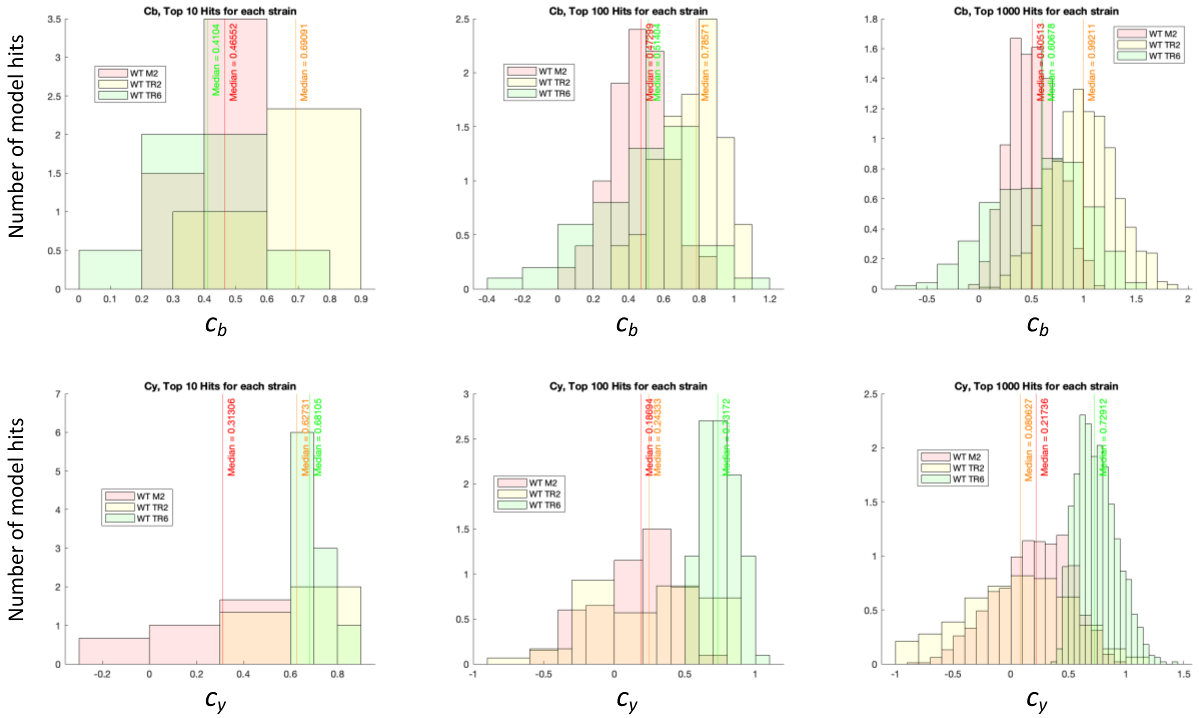


Example of parameter exploration (see SI discussion on modeling strategy) using Latin Hypercube Sampling of ecological overlap terms $c$ for each WT cell pairing at 100% LW. Each histogram color represents a different plasmid combination for WT donor cells (E), to account for slight fitness differences. Columns show top 10, 100, and 1000 parameter guesses for $c_b$ (top) and $c_y$ (bottom), after ranking for lowest error compared to experimental data, based on a simplified version of free-cell model that omits amino-acid feeding, IDC, and death rates. Means of these distributions served as initial guesses in the full free-cell model, and are meant to demonstrate the wider range of parameters explored before the full model was fit.

#### SI Figure 9: Free-cell model fits for bacteria and yeast in coculture

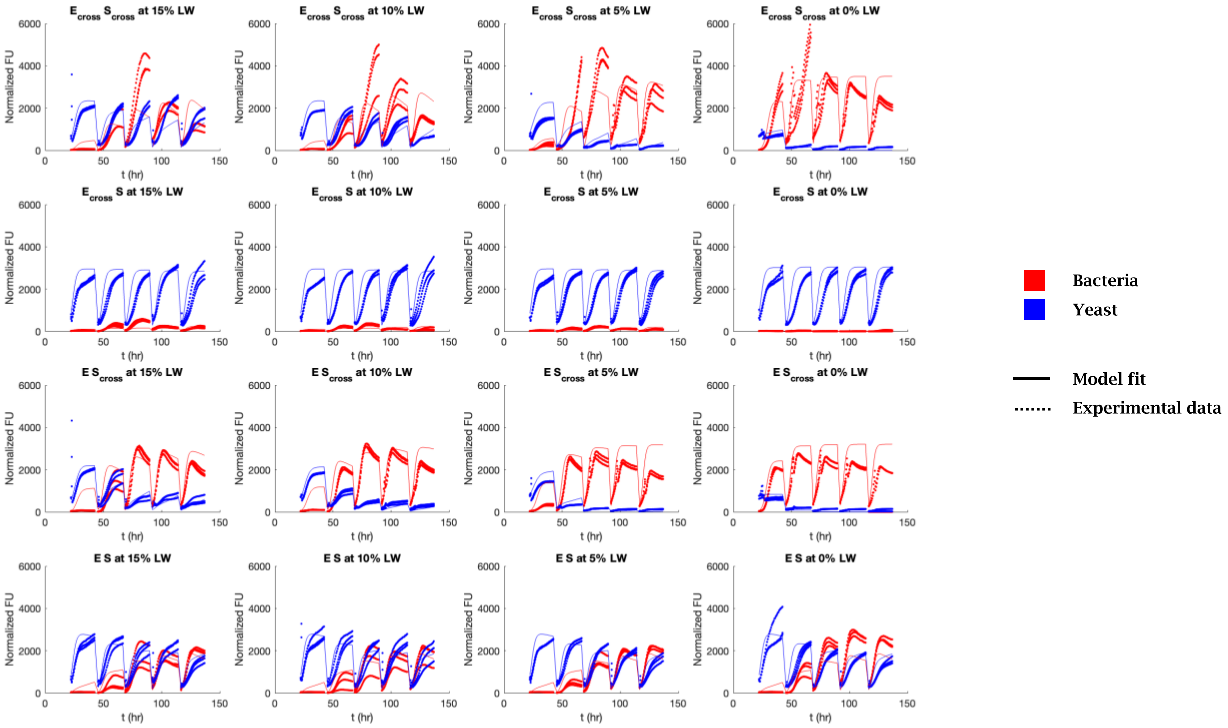


Fluorescent data (dots) of bacteria (red) and yeast (blue) with mannose is compared to predictions (lines) from free-cell model, given parameters listed in Table S1. Each cell pairing (rows) and LW% (columns) are shown over six days of batch culturing. Free-cell model was fit to prioritize three specific outcomes: 1) susceptibility of each strain to changes in amino acid concentration, 2) steady-state persistence of each strain, and 3) approximate D:R ratio of cells.

#### SI Figure 10: IDC rate-term sweep for “free” cell model

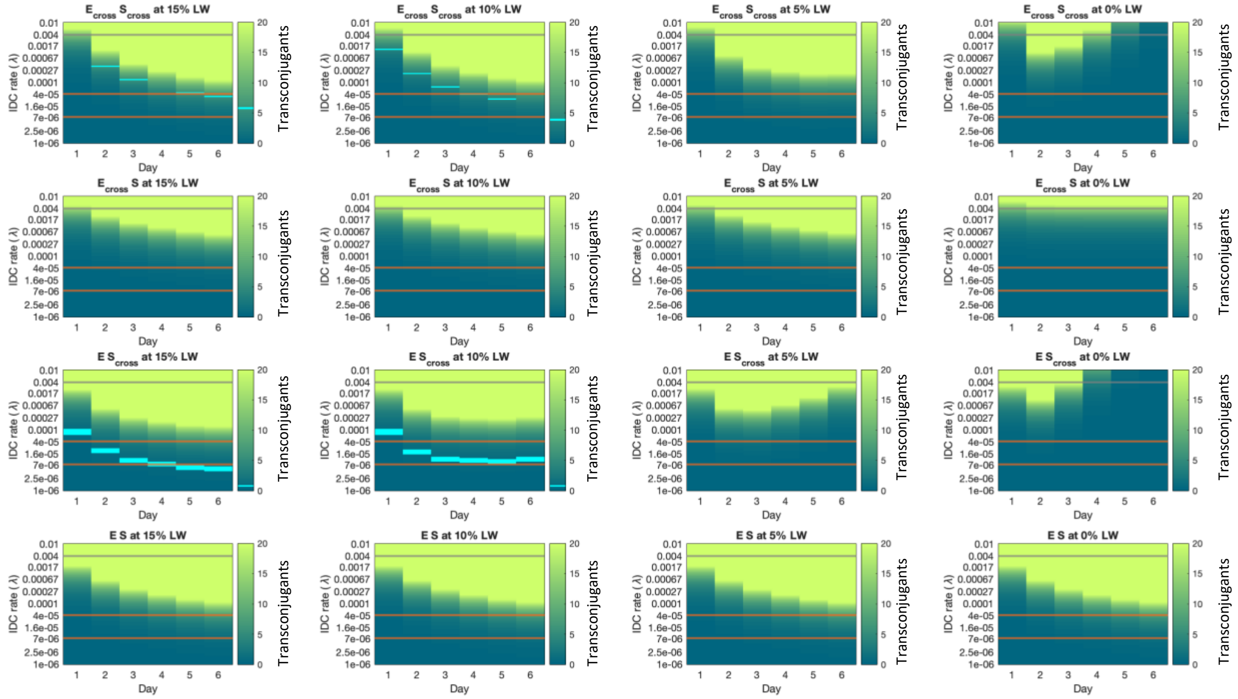


Heatmaps showing predicted number of transconjugants (color) for a range of IDC rates $\gamma$ (y-axis) over six days of batch culturing, for all experimental conditions, assuming cells were unable to clump, and thus conjugated via random collisions. Cyan heat markers represent experimental IDC counts for the four conditions that had counts above zero with mannose, i.e. in the unclumped samples. Gray line at $\gamma$=0.004 represents literature prediction for enteric E. coli IDC rate, orange lines represent range of IDC-rate values matching data, roughly between 7*10-6 and 4*10-5.

#### SI Figure 11: Clumped-cell model fits for free cells, clumped cells, and number of clumps

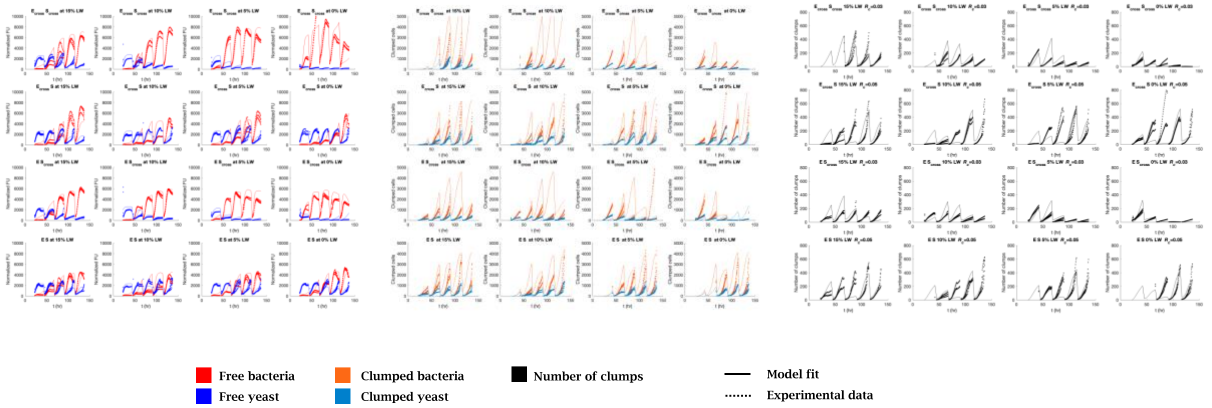


Left: fluorescent data (dots) of free bacteria (red) and yeast (blue) without mannose is compared to predictions (lines) from clumped model, given parameters listed in Table S2. Each cell pairing (rows) and LW% (columns) are shown over six days of batch culturing. Middle: image analysis estimates for clumped bacteria (orange) and clumped yeast (blue) with overlain model fits (lines) for each condition. Right: image analysis estimates for number of clumps (dots) with model predictions overlain (lines) for each condition.

#### SI Figure 12: IDC rate-term sweep for "clumped" cell model.

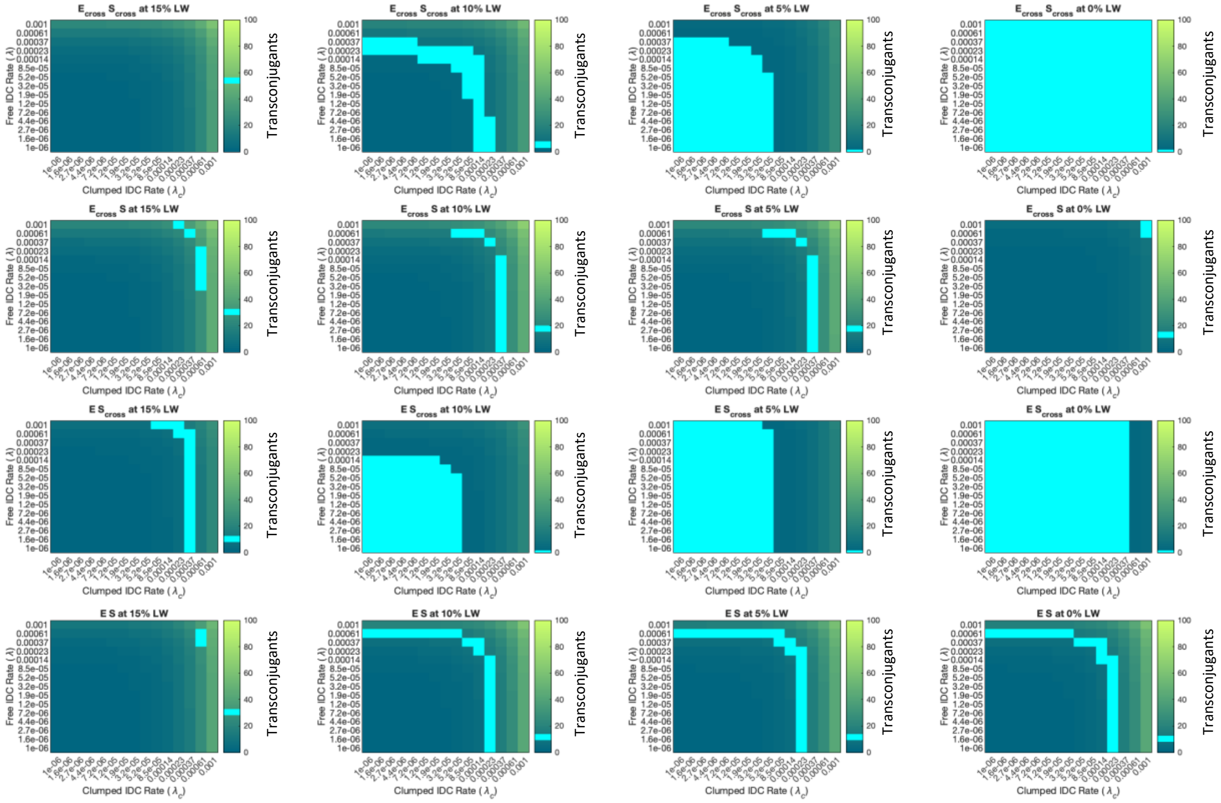


Heatmaps showing predicted number of transconjugants (color) for a range of "free" IDC rates $\gamma$ (y-axis) and "clumped" IDC rates $\gamma_c$ (x-axis), for all experimental conditions at day six of batch culturing. Cyan boxes represent mean experimental IDC count of per 100 uL culture (three replicates), with several values of $\gamma$ and $\gamma_c$ resulting in these numbers of transconjugants in many cases. Two main categories of rates yield the experimental IDC results for many experimental conditions: low $\gamma_c$ with $\gamma$ above $5*10^{-4}$, or low $\gamma$ with $\gamma_c$ near $3*10^{-4}$. Because free-model results showed $\gamma$ below $5*10^{-5}$, it's likely that the latter case is true, with most conjugation resulting from clumped interactions.

#### SI Figure 13: Radial fluorescence measurements, mixed colony experiment

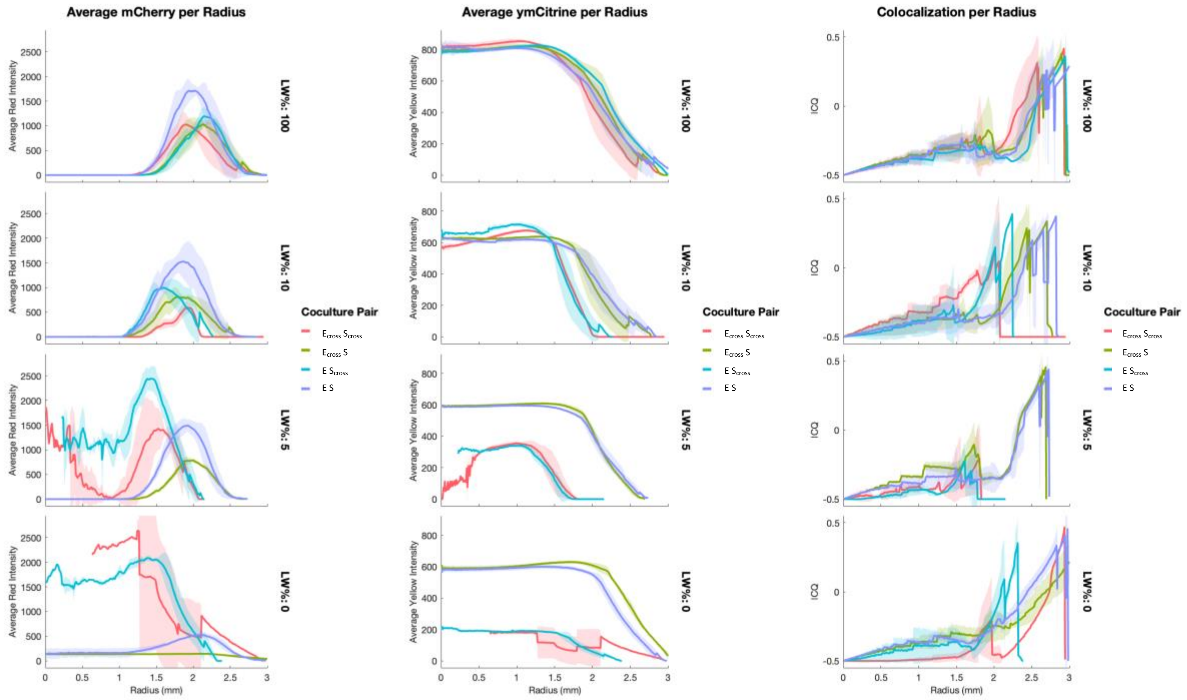


Radial measurements of mixed colony bacteria (via mCherry, left column), yeast (via ymCitrine, middle column), and colocalization between the two cell signals (right column), for each pairing (color) and LW% (rows) after 6 days of growth. Lines are means of all pixels measured at a given radial shell (number of pixels varied per colony image and radius), shading is standard deviation. Colocalization is measured by intensity correlation quotient (ICQ, see SI discussion) for each radius.

#### SI Figure 14: Jackpot transconjugant population along expansion front in mixed colony

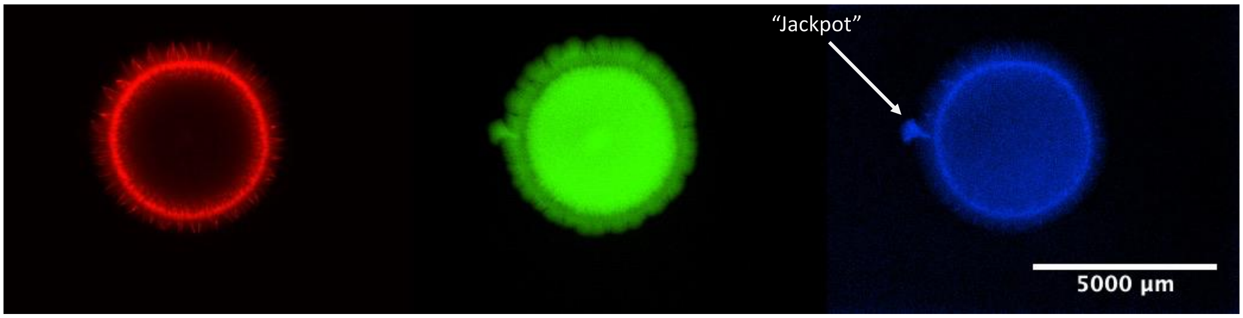


Bacteria (mCherry) Yeast (ymCitrine) Transconjugants (yeBFP)

Fluorescence images of WT (E S) mixed colony at 10% LW, after six days of growth, with a *cis*-transfer donor. Bacterial (mCherry, left), yeast (ymCitrine, middle), and IDC (yeBFP, right. See SI discussion on colony analysis for IDC-reporter details) channels are shown. Note that while yeBFP signal is largely convoluted by autofluorescence from bacteria (especially) and yeast, a protrusion on the left side of the colony shows some of the brightest yeBFP and doesn't correspond to bacterial signal. This colony showed "jackpot" IDC counts > 500 CFU. Scale bar = 5 mm.

### SI Figure 15: Growth trajectories for rescue assay

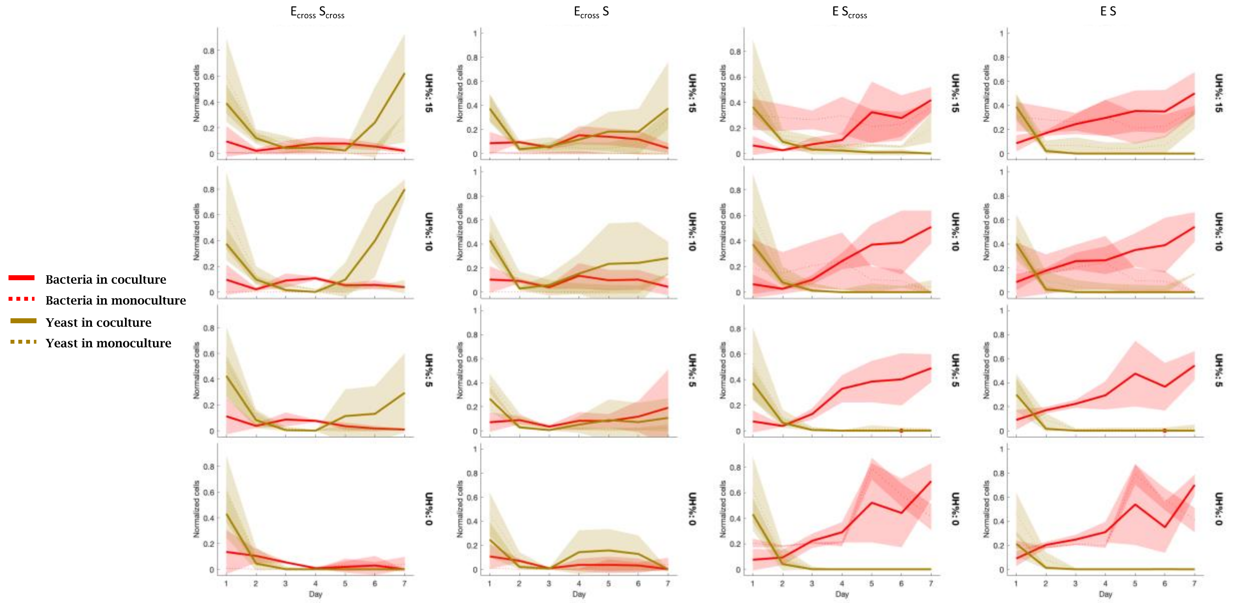


Normalized flow cytometry data for each cell pairing (columns) at four yeast-dependent amino acid concentrations (UH%, rows), for seven days of batch culturing; all samples have 0% L to make $E_{cross}$ cells dependent on yeast. Bacterial cell counts shown in red, yeast counts in yellow. Solid lines represent coculture traces, which are matched with each cell's monoculture traces (dotted) for comparison; shading is standard deviation (six replicates coculture, five replicates monoculture).

#### SI Figure 16: Stochasticity in rescue outcomes at apparent critical point

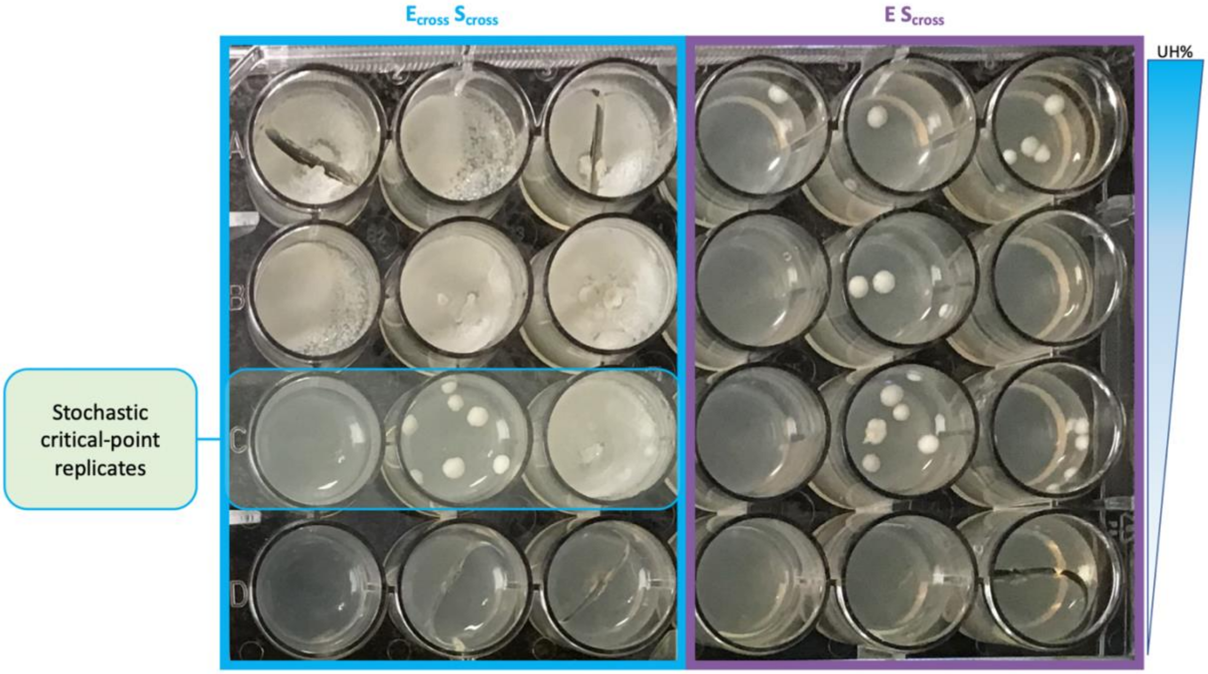


IDC plate for one rescue experiment at day six. Left three columns are $E_{cross}$-$S_{cross}$, right three columns E-$S_{cross}$, with each row a different UH% (top to bottom: 15%, 10%, 5%, 0%); each condition shown has three replicates, and all samples have 0% L. Note that at 5% UH, three $E_{cross}$-$S_{cross}$ replicates give drastically different outcomes, ranging from collapse (well C1) to full rescue (well C3). We predict this corresponds to the boundary between red and green conditions in the phase map (Fig 6d).

#### SI Figure 17: sgRNA testing for IDC-mediated killing

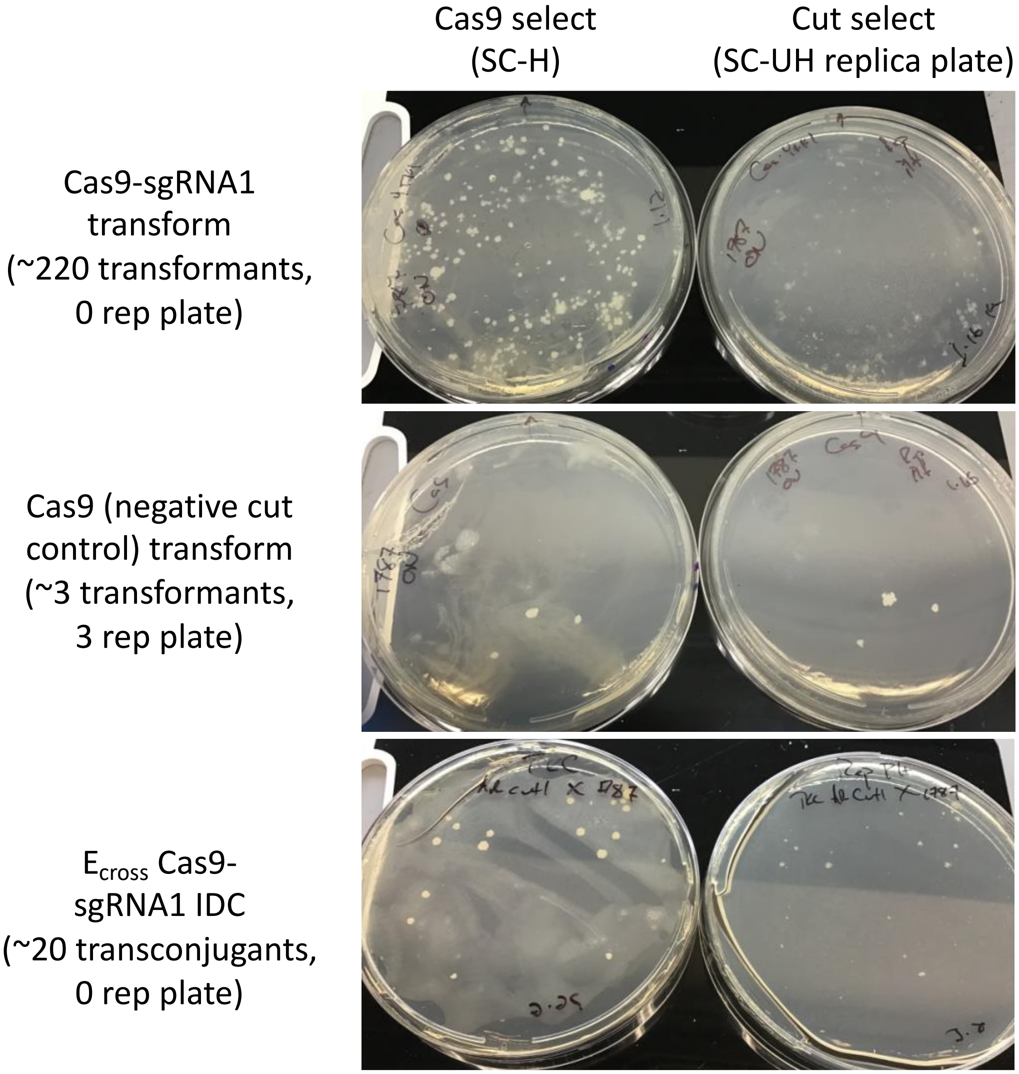


Transformant or IDC selection plates (SC-H, left) and corresponding cut selection replica plates (SC-UH, right). Top row: Cas9-sgRNA was transformed directly into yMM1787, resulting in ~220 colonies on SC-H, none of which replica plated on SC-UH, demonstrating complete cutting of *BFP-URA3* plasmid. Middle row: Cas9 (no sgRNA) negative control was transformed directly into yMM1787, showing only three colonies on SC-H, all of which replica-plated on SC-UH, demonstrating no cutting. Bottom row: Cas9-$ori^T$-sgRNA was conjugated into yMM1787 via $E_{cross}$ IDC donor, showing ~20 transconjugants, none of which replica plated on SC-UH, demonstrating complete cutting of *BFP-URA3* plasmid via IDC.

#### SI Figure 18: Growth trajectories for IDC-killing assay

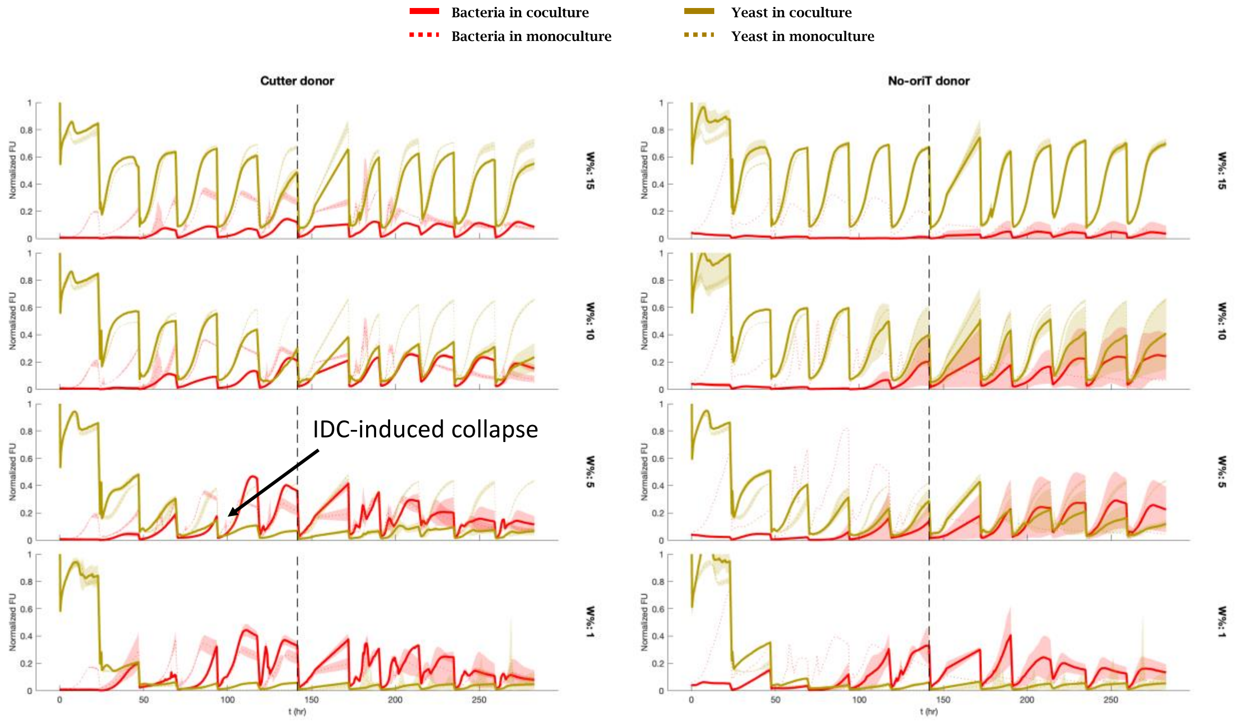


Fluorescence data for each cell pairing (columns), normalized to max per channel after day 1, at four yeast-dependent amino acid concentrations (W%, rows), for 12 days of batch culturing. All samples shown are grown in 0% U such that yeasts that receive the IDC-Cas9 plasmid are cut and can no longer grow. Bacterial (normalized mCherry) traces shown in red, yeast (normalized ymCitrine) counts in yellow. Solid lines represent coculture traces, which are matched with each cell's monoculture traces (dotted) for comparison; lines are mean of three replicates, shading is standard deviation. Vertical dotted lines represent addition of mannose to media, which prevents clumping, after which yeast growth appears to level off.

#### SI Figure 19: Microscopy images of IDC-killing cocultures pre and post mannose addition

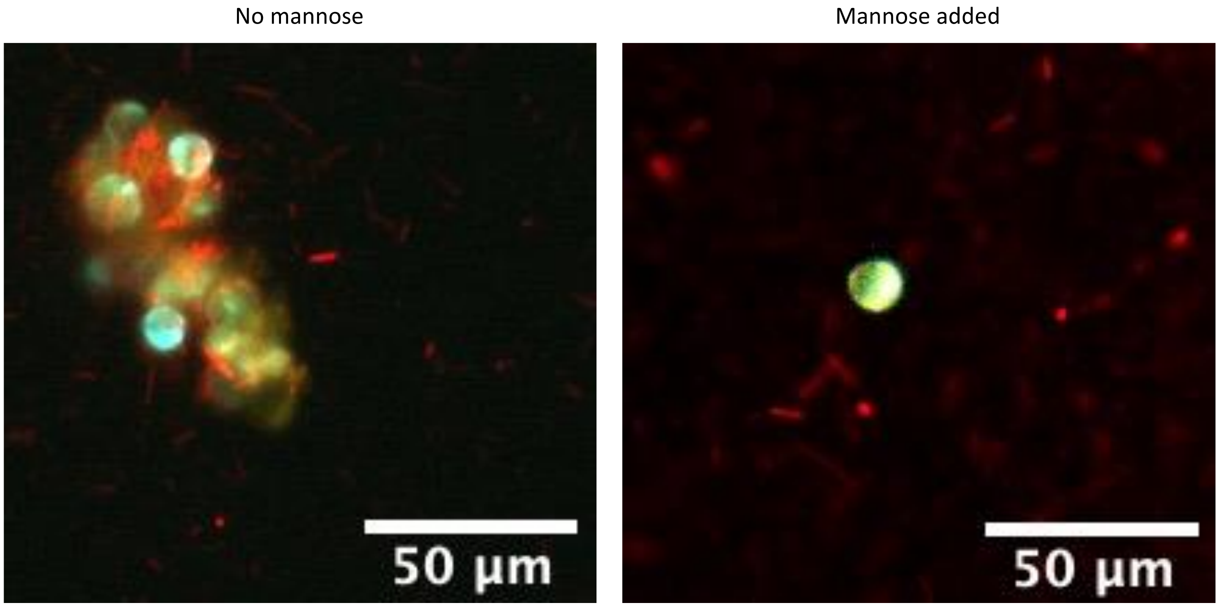


10x images of cocultures from IDC-killing assay, showing clumping before mannose addition (left, day 5), vs. non-clumped cells after mannose addition (right, day 7); mannose was added after day 6. Cells shown are cut-donors and yMM1787, at 5% W 100% U. Brightness for each channel (red, yellow, and blue) was manually scaled to visualize cell distribution. Scale bars = 50 µm.

#### SI Figure 20: Plasmid loss for IDC-killing assay without cut selection conditions

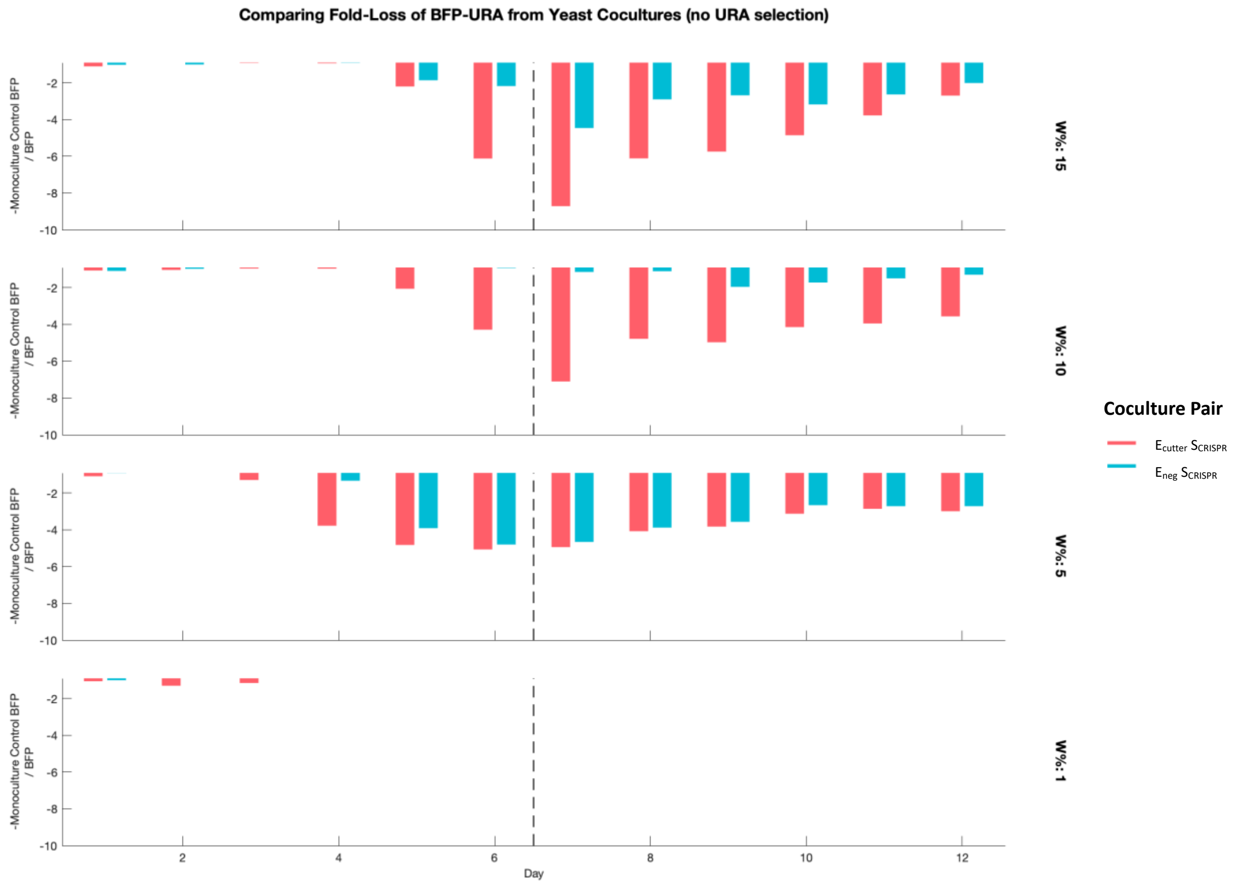


Fold-loss of *BFP-URA3* plasmid in 100% U—in which cut yeasts can still grow—for each coculture relative to yMM1787 (SCRISPR) monoculture, at four different W% (rows) over 12 days of batch culture. Because yeasts are prone to lose the *BFP-URA3* plasmid without uracil selection, these demonstrate additional loss of this plasmid due to cutting. Fold decrease measured by – (Monoculture BFP signal) / (Coculture BFP signal). Note that cut-donors (red) drive yeast to extinction in 5% W at around day four, and so parity with negative-control donors (blue) after that is likely an artifact of overall low BFP signal. Vertical dotted lines denote addition of mannose, which prevents clumping.

### REFERENCES


1. Kurita, T., Otsu, N., and Abdelmalek, N. (1992). Maximum likelihood thresholding based on population mixture models. Pattern Recognition *25*, 1231–1240. 10.1016/0031-3203(92)90024-D.

2. Li, Q., Lau, A., Morris, T.J., Guo, L., Fordyce, C.B., and Stanley, E.F. (2004). A Syntaxin 1, Gαo, and N-Type Calcium Channel Complex at a Presynaptic Nerve Terminal: Analysis by Quantitative Immunocolocalization. J Neurosci *24*, 4070–4081. 10.1523/JNEUROSCI.0346-04.2004.

3. Muller, M.J.I., Neugeboren, B.I., Nelson, D.R., and Murray, A.W. (2014). Genetic drift opposes mutualism during spatial population expansion. Proceedings of the National Academy of Sciences *111*, 1037–1042. 10.1073/pnas.1313285111.

4. Fusco, D., Gralka, M., Kayser, J., Anderson, A., and Hallatschek, O. (2016). Excess of mutational jackpot events in expanding populations revealed by spatial Luria–Delbrück experiments. Nat Commun *7*, 12760. 10.1038/ncomms12760.

5. Zhang, B., DeAngelis, D.L., and Ni, W.-M. (2021). Carrying Capacity of Spatially Distributed Metapopulations. Trends in Ecology & Evolution *36*, 164–173. 10.1016/j.tree.2020.10.007.

6. Hoek, T.A., Axelrod, K., Biancalani, T., Yurtsev, E.A., Liu, J., and Gore, J. (2016). Resource Availability Modulates the Cooperative and Competitive Nature of a Microbial Crossfeeding Mutualism. PLOS Biology *14*, e1002540. 10.1371/journal.pbio.1002540.

7. Wu, F., Lopatkin, A.J., Needs, D.A., Lee, C.T., Mukherjee, S., and You, L. (2019). A unifying framework for interpreting and predicting mutualistic systems. Nat Commun *10*. 10.1038/s41467-018-08188-5.

8. Levin, B.R., Stewart, F.M., and Rice, V.A. (1979). The kinetics of conjugative plasmid transmission: Fit of a simple mass action model. Plasmid *2*, 247–260. 10.1016/0147-619X(79)90043-X.

9. Simonsen, L., Gordon, D.M., Stewart, F.M., and Levin, B.R. (1990). Estimating the rate of plasmid transfer: an end-point method. Microbiology, *136*, 2319–2325. 10.1099/00221287-136-11-2319.

10. Zhong, X., Droesch, J., Fox, R., Top, E.M., and Krone, S.M. (2012). On the meaning and estimation of plasmid transfer rates for surface-associated and well-mixed bacterial populations. Journal of Theoretical Biology *294*, 144–152. 10.1016/j.jtbi.2011.10.034.

11. Zhao, Z.-J., Zou, C., Zhu, Y.-X., Dai, J., Chen, S., Wu, D., Wu, J., and Chen, J. (2011). Development of l-tryptophan production strains by defined genetic modification in Escherichia coli. J Ind Microbiol Biotechnol *38*, 1921–1929. 10.1007/s10295-011-0978-8.

12. Baba, T., Ara, T., Hasegawa, M., Takai, Y., Okumura, Y., Baba, M., Datsenko, K.A., Tomita, M., Wanner, B.L., and Mori, H. (2006). Construction of Escherichia coli K-12 in-frame, single-gene knockout mutants: the Keio collection. Mol Syst Biol *2*, 2006.0008. 10.1038/msb4100050.

13. Datsenko, K.A., and Wanner, B.L. (2000). One-step inactivation of chromosomal genes in Escherichia coli K-12 using PCR products. Proc Natl Acad Sci U S A *97*, 6640–6645.

14. Soltysiak, M.P.M., Meaney, R.S., Hamadache, S., Janakirama, P., Edgell, D.R., and Karas, B.J. (2019). Trans-Kingdom Conjugation within Solid Media from Escherichia coli to Saccharomyces cerevisiae. International Journal of Molecular Sciences *20*, 5212. 10.3390/ijms20205212.

15. Lee, M.E., DeLoache, W.C., Cervantes, B., and Dueber, J.E. (2015). A Highly Characterized Yeast Toolkit for Modular, Multipart Assembly. ACS Synth. Biol. *4*, 975–986. 10.1021/sb500366v.
